# Tailored chemical reactivity probes for systemic imaging of aldehydes in fibroproliferative diseases

**DOI:** 10.1101/2023.04.20.537707

**Authors:** Hua Ma, Iris Y. Zhou, Y. Iris Chen, Nicholas J. Rotile, Ilknur Ay, Eman Akam, Huan Wang, Rachel Knipe, Lida P. Hariri, Caiyuan Zhang, Matthew Drummond, Pamela Pantazopoulos, Brianna F. Moon, Avery T. Boice, Samantha E. Zygmont, Jonah Weigand-Whittier, Mozhdeh Sojoodi, Romer A. Gonzalez-Villalobos, Michael K. Hansen, Kenneth K. Tanabe, Peter Caravan

**Affiliations:** Athinoula A. Martinos Center for Biomedical Imaging, Institute for Innovation in Imaging (i^3^), Department of Radiology, Massachusetts General Hospital, Harvard Medical School, Boston, USA; Division of Pulmonary and Critical Care Medicine and the Center for Immunology and Inflammatory Diseases, Massachusetts General Hospital, Boston, MA, USA; Department of Pathology, Massachusetts General Hospital, Harvard Medical School, Boston, USA; Division of Gastrointestinal and Oncologic Surgery, Massachusetts General Hospital, Harvard Medical School, Boston, Massachusetts 02114, United States; Cardiovascular and Metabolism Discovery, Janssen Research and Development LLC, Boston, MA, USA

## Abstract

During fibroproliferation, protein-associated extracellular aldehydes are formed by the oxidation of lysine residues on extracellular matrix proteins to form the aldehyde allysine. Here we report three Mn(II)-based, small molecule magnetic resonance (MR) probes that contain α-effect nucleophiles to target allysine in vivo and report on tissue fibrogenesis. We used a rational design approach to develop turn-on probes with a 4-fold increase in relaxivity upon targeting. The effects of aldehyde condensation rate and hydrolysis kinetics on the performance of the probes to detect tissue fibrogenesis noninvasively in mouse models were evaluated by a systemic aldehyde tracking approach. We showed that for highly reversible ligations, off-rate was a stronger predictor of in vivo efficiency, enabling histologically validated, three-dimensional characterization of pulmonary fibrogenesis throughout the entire lung. The exclusive renal elimination of these probes allowed for rapid imaging of liver fibrosis. Reducing the hydrolysis rate by forming an oxime bond with allysine enabled delayed phase imaging of kidney fibrogenesis. The imaging efficacy of these probes, coupled with their rapid and complete elimination from the body, make them strong candidates for clinical translation.

## Main

Electrophilic carbonyls such as ketones and aldehydes are generally considered rare in mammalian biology, existing at low concentrations in homeostasis and only transiently due to their reactivity in a nucleophilic environment. This is especially true in the extracellular space^1, 2^. Indeed, the scarcity of extracellular carbonyls has led to the development of bioorthogonal conjugation chemistry involving aldehydes. For example, Bertozzi’s early work on modifying cell surfaces relied on the integration of keto sialic acid to cell-surface glycans, which then can be covalently ligated by hydrazine or oxyamine under physiological conditions^3, 4^. Paulson and co-workers introduced aldehydes into cell-surface sialic acid residues and then captured the modified glycoproteins by reaction with aminooxybiotin followed by streptavidin chromatography^5^. Taking advantage of the small size and compatibility in living systems, aldehyde/ketone have been utilized as labeling blocks or targeting warheads for bioorthogonal ligation^6-8^.

An important aldehyde in the extracellular matrix (ECM) is the amino acid allysine which is formed by the oxidation of lysine residues by the enzyme lysyl oxidase (LOX) and its paralogs^9^. Allysine formed on collagens (or elastin) undergoes a series of condensation and rearrangement reactions with other collagens that result in irreversible bonds that crosslink the proteins and stabilize the ECM. Outside of development and wound healing, tissue levels of LOX and allysine are low in healthy, mature mammals. However, if the tissue is damaged then LOX is upregulated while the tissue is being remodeled^10^. We have found that lung and liver tissue allysine concentrations can be in the hundreds of micromolar during such pathological processes^11, 12^.

About half of mortality in the industrialized world is caused by a disease that has a fibroproliferative component^13, 14^. Fibrosis is characterized by tissue scarring, and as functional tissue is replaced by scar, the organ becomes less compliant and dysfunctional. When progressive, fibrosis leads to organ failure and/or death. In organs like liver, fibrosis has been shown to promote tumor formation^15^. Many cancers also have a fibrotic component that is believed to promote tumor growth and shield cancer from chemo-or immunotherapy^16^. Common diseases associated with fibrosis are non-alcoholic steatohepatitis (NASH)^17^, chronic kidney disease (CKD)^18^, heart failure (Fig. 1a)^19^, but there are also less common fibrotic diseases such as idiopathic pulmonary fibrosis (IPF)^20^, and all of them cause significant morbidity and mortality. A major unmet need across all of these diseases is the ability to measure disease activity noninvasively^21, 22^, which could help identify the early onset of disease, could distinguish between an active disease that is progressing from stable scar, could provide prognostic information, or could be used to monitor response to therapy.

**Fig. 1.**
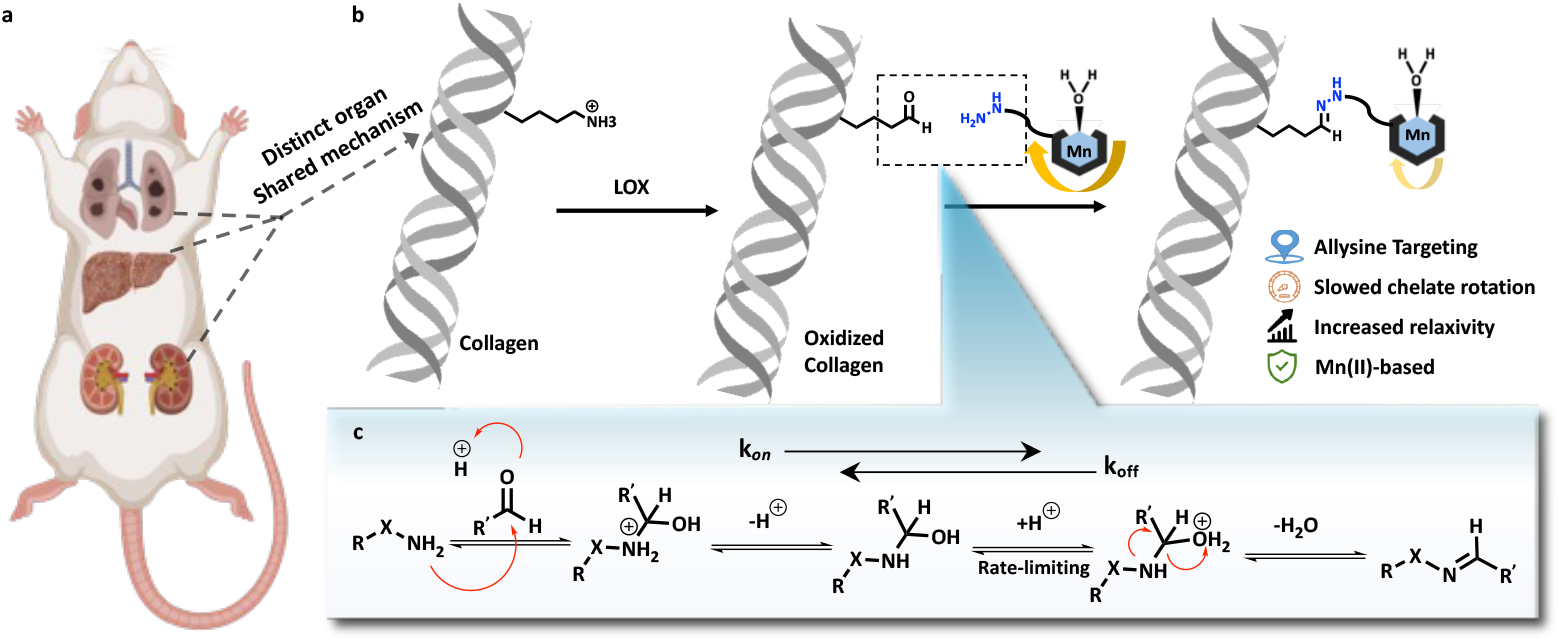
Chemical reactivity probes can systemically target aldehydes for quantitative molecular imaging of fibrogenesis. **a**, Fibrosis can occur in any organ and is characterized by spatially heterogeneous fibrotic foci, wherein lysine residues are converted to the aldehyde allysine while the adjacent tissue remains normal. Precise monitoring of fibrotic disease activity calls for fine-tuning of targeting kinetics. Created with BioRender.com. **b**, Taking advantage of the condensation reaction between hydrazine (or oxyamine) and aldehyde, allysine-targeted Mn(II)-based molecular MR imaging probes can be rationally designed with relaxivity “turned on” upon binding for increased detection sensitivity. **c**. Breakdown of the tetrahedral intermediate is the rate-limiting step of condensation reaction between nucleophile (hydrazine or oxyamine) and aldehyde, providing a large variation in reaction kinetics to meet the organ-specific physiological demands.

The high concentration of allysine during fibrogenesis makes it amenable to detection by a targeted magnetic resonance (MR) probe molecule^12, 23-26^. The general absence of aldehydes in the ECM of healthy tissue suggests that an extracellular, aldehyde-targeted probe would have high specificity for fibroproliferative disease activity. There are several considerations in designing a molecular probe for noninvasive sensing of extracellular aldehydes in the body by MRI^27^. The probe should produce a strong MR signal change and requires an appropriate MR signal generator like a Gd^3+^ or Mn^2+^ chelate. The probe should be metabolically stable in vivo such that the MR signal reflects the distribution of the probe and not metabolites. For liver imaging applications, it is important that the probe not accumulate in the liver or undergo hepatobiliary elimination, as this would increase the background signal. Since MR relaxation probes function by changing the existing MR signal, i.e. they are contrast agents, it is necessary to acquire an image before and after administering the probe to see the signal change induced by the probe. The ideal probe would accumulate rapidly and be retained at its target immediately after injection, but the unbound probe would be rapidly eliminated to minimize nonspecific signal enhancement. Because the dose required for molecular MRI is fairly high (mg metal ion per kg body weight), it is also important that the probe is ultimately eliminated from the body. Finally, since we are targeting extracellular aldehydes, it is important that the probe have an extracellular distribution to minimize any background signal from intracellular aldehydes.

In this study, we describe three novel, extracellular aldehyde targeting probes that comprise a Mn^2+^ chelate conjugated to an aldehyde reactive moiety. As an aldehyde reactive group, we used either alkyl hydrazine, an alpha-carboxylate alkyl hydrazine, and an alpha-carboxylate oxyamine. These designs enabled us to examine the effects of aldehyde condensation rate and condensation product hydrolysis rate on the in vivo performance of the probes for aldehyde sensing in mouse models of lung, liver, and kidney fibrosis.

## Results

### Design and characterization of aldehyde-targeting MR probes

We designed our probes, Fig. 2a, on the symmetric macrocyclic Mn(PC2A)(H_2_O) core which contains one rapidly exchanging coordinated water co-ligand and is thermodynamically stable and kinetically inert with respect to Mn^2+^ dissociation in vivo^28^. The PC2A chelator was readily derivatized at the N7 position to introduce an aldehyde reactive group, either an alkyl hydrazine or oxyamine moiety, as these groups undergo fast reversible condensation reactions with aldehydes. Hydrazines typically have faster aldehyde condensation rates than oxyamines at neutral pH, but the hydrolysis rate of the resultant hydrazone is much faster than that of an oxime. For the condensation reaction, dehydration of the tetrahedral intermediate is typically the rate-limiting step and the reaction proceeds under general acid catalysis^29^. Substitution of an acidic moiety adjacent to the hydrazine/oxyamine will accelerate the condensation reaction but may also catalyze the back reaction. MnL1 and MnL2 allow for the direct comparison between a hydrazine and an oxyamine. MnL1 and MnL3 allow for the direct comparison of the effect of the α-carboxylate of reaction kinetics and subsequent in vivo performance. We also compare MnL2 to the historic compound GdOA to assess the effect of the α-carboxylate on oxyamine condensation/hydrolysis kinetics. MnL4 was synthesized as a structurally matched negative control, which should have similar pharmacokinetic properties as MnL1, MnL2, and MnL3, but is incapable of undergoing condensation with allysine. All the complexes are very hydrophilic which should minimize non-specific protein binding and reduce the fraction of hepatobiliary elimination.

**Fig. 2.**
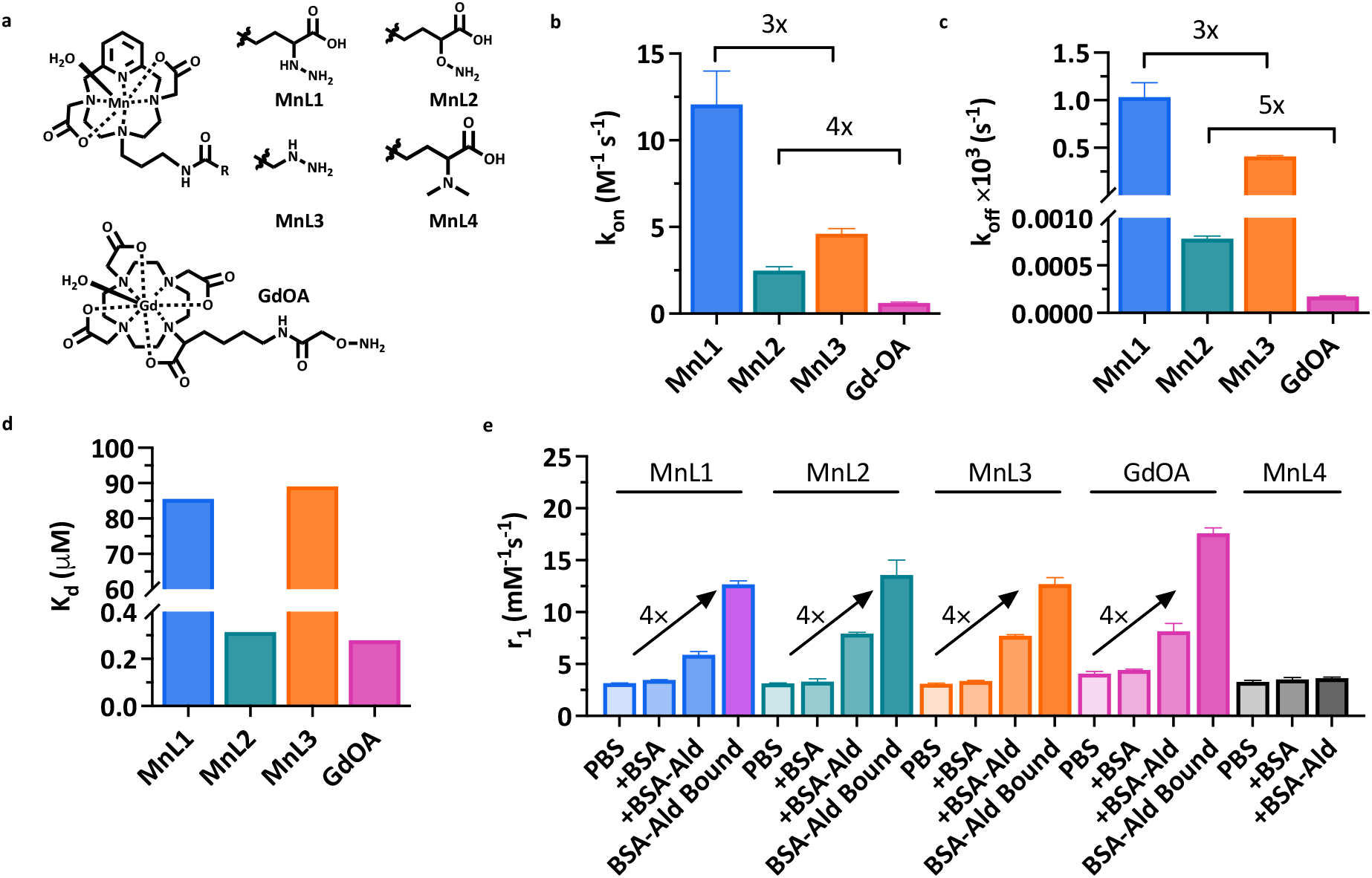
Design and characterization of extracellular aldehyde-targeted MR probes. **a**, Structure of novel MnPC2A-based complexes and the known compound GdOA. **b**, Second order rate constant (k_on_) for the reaction of MnL1, MnL2, MnL3, and GdOA with butyraldehyde (pH 7.4, PBS, 25 °C; data presented as mean ± s.d., n = 3 independent samples). **c**, Hydrolysis rate constant (k_off_) of condensation products (data presented as mean ± s.d., n = 3 independent samples). **d**, Dissociation constant (K_d_) of condensation reaction between hydrazine/oxyamine-based probes with butyraldehyde calculated as the ratio k_off_/k_on_. **e**, Relaxivities of MnL1, MnL2, MnL3, MnL4, and GdOA in PBS, in BSA, in allysine-modified BSA-Ald, and of the BSA-Ald bound species (1.4 T, 37°C. Data presented as mean ± s.d., n = 3 independent samples).

Each of the Mn(II) complexes was synthesized starting from the macrocycle pyclen, followed by regioselective alkylation of the N4 and N10 positions using *tert*-butyl-bromoacetate at pH 8. Next, the N7-position was alkylated with *N*-(3-Bromopropyl)phthalimide, and the primary amine liberated by reaction with hydrazine. This common intermediate was used to prepare each of the Mn(II) probes by conventional amide bond formation with the appropriate synthon to introduce α-carboxylate substituted hydrazine, α-carboxylate substituted oxyamine, terminal hydrazine, or dimethylamine group. After TFA-promoted deprotection and subsequent Mn(II) chelation, we accessed the four Mn(II) probes. The synthetic procedures are fully described in Supplementary Information. Each compound was purified and characterized using standard methods (preparative HPLC, ^1^H NMR, ^13^C NMR, LC-MS, ICP-MS, and LC-ICP-MS; please see Methods and Supplementary Information for details).

The condensation reaction rate constants with butyraldehyde were measured by UV spectroscopy under pseudo-first-order conditions with respect to each probe (Supplementary Information). As shown in Fig. 2b and Supplementary Fig. 1, introduction of the α-carboxylate moiety led to 3 – 4 fold higher condensation reaction rates: MnL1 had a second-order rate constant of 12.1 ± 1.9 M^-1^s^-1^ compared to 4.6 ± 0.3 M^-1^s^-1^ for MnL3; MnL2 had a second-order rate constant of 2.5 ± 0.2 M^-1^s^-1^ compared to 0.6 ± 0.1 M^-1^s^-1^ for GdOA. This result confirmed the hypothesis that intramolecular acidic condition would facilitate the condensation reaction between hydrazine/oxyamine and aldehyde. Then, the hydrolysis of hydrazone and oxime products with butyraldehyde were investigated by HPLC. Following Kalia and Raines^30^, we used an excess of formaldehyde to trap the liberated nitrogen base and thereby push the hydrolysis reaction to completion without interference from the reverse (condensation) reaction. As shown in Fig. 1c and Supplementary Fig. 2, MnL1-hydrazone hydrolysis is 3-times faster than MnL3-hydrazone, and MnL2-oxime hydrolysis is 5-times faster than GdOA. In other words, the α-carboxylate moiety accelerates condensation and hydrolysis to a similar extent. Measurement of the on- and off-rate constants allowed us to compute the dissociation constant (Kd) as the ratio k_off_/k_on_ (Fig. 1d). The much slower off-rate of the oxime hydrolysis resulted in 2 orders of magnitude lower K_d_ for MnL2 and GdOA compared to MnL1 and MnL3. However, since the α-carboxylate moiety accelerates condensation and hydrolysis equally, there was little difference in K_d_ between MnL1 and MnL3.

MR probes are characterized by their relaxivity, r_1_, which is the change in solvent 1/T_1_ caused by adding the probe, normalized to the probe concentration. We measured relaxivity (1.4 T, 37 °C, pH 7.4) of each complex in phosphate-buffered saline (PBS) and then again in PBS containing bovine serum albumin (BSA). The four Mn complexes exhibited similar r_1_ values (about 3.1 mM^-1^s^-1^) in PBS, consistent with one coordinated water ligand (Fig. 1e). The presence of a single coordinated water co-ligand was confirmed by temperature-dependent H_2_^17^O transverse relaxation rates using the method of Gale et al. (Supplementary Fig. 3)^31^. In the presence of excess BSA there was little to no relaxivity enhancement, indicating low nonspecific protein binding. We next measured the relaxivity of each complex in the presence of oxidized BSA wherein about 4 lysine side chains on BSA were oxidized to allysine, and we used this allysine-modified BSA (BSA-Ald, 43.8 mg mL^-1^, 1.2 mM aldehyde) as water-soluble model aldehyde bearing protein^32, 33^. In the presence of BSA-Ald (3 h incubation, 37°C), the relaxivity increased by 90%, 150%, 150%, and 100% (5.9 mM^-1^s^-1^, 7.9 mM^-1^s^-1^, 7.7 mM^-1^s^-1^, and 8.2 mM^-1^s^-1^) for MnL1, MnL2, MnL3, and GdOA, respectively, relative to their relaxivity values in PBS. We then separated the unbound probe from the BSA-Ald bound probe by ultrafiltration with a molecular weight cutoff filter (Supplementary Fig. 4) and then measured the relaxivity of the protein-bound species. The protein-bound relaxivities were 4 times higher than the relaxivity of the unbound probe (12.7 mM^-1^s^-1^, 13.6 mM^-1^s^-1^, 12.7 mM^-1^s^-1^, and 17.6 mM^-1^s^-1^ for MnL1, MnL2, MnL3, and GdOA respectively). In comparison, the relaxivities of non-reactive control probe MnL4 were unchanged from PBS to BSA or BSA-Ald, indicating a lack of protein binding for the control probe. At a higher magnetic field of 4.7 T where the imaging studies were performed, the relaxivity of BSA-Ald bound probes was also significantly enhanced compared to the unbound form (113% increase for MnL1, 115% increase for MnL2, 115% increase for MnL3, Supplementary Fig. 5).

The thermodynamic stability constant of Mn^2+^ with the unmodified PC2A ligand was reported to be log K^MnPC2A^ = 17.09^28^, and we expect similar stability with the complexes here. We assessed the kinetic inertness of the 4 Mn complexes described here by transmetalation experiments with an excess of Zn^2+^, which should form more stable complexes than Mn^2+^ based on its position in the Irving-Williams series^34^. The Mn(II) complexes were incubated with 25 equivalents of Zn^2+^ at 37 °C, in pH 6.0 MES buffer, and changes in the paramagnetic longitudinal relaxation rate (%R_1_^P^) were used as readout. Under this condition, any free Mn^2+^ released by chelator will cause an increase in R_1_^P^ owing to the higher relaxivity of the Mn^2+^ aqua ion. Mn-PyC3A, which is currently being developed as an MRI probe and was shown to be excreted from animals in its intact form, was measured as a comparative benchmark^35^. Measured pseudo-first-order rate constants were 2.5 ± 0.1 × 10^−4^, 2.4 ± 0.3 ×10^−4^, 2.3 ± 0.3 ×10^−4^, 2.2 ± 0.4 ×10^−4^, and 5.5 ± 0.4 ×10^−4^ s^-1^ for MnL1, MnL2, MnL3, MnL4, and Mn-PyC3A, respectively (Supplementary Fig. 6). Since the Mn-PC2A derivatives reported here are twice as inert as Mn-PyC3A, we expect these complexes to also be stable in vivo with respect to Mn^2+^ dissociation.

The stabilities of MnL1, MnL2, MnL3, and MnL4 in human plasma were also assessed using HPLC coupled to ICP-MS (LC-ICP-MS) to measure the formation of any new Mn-containing species (Supplementary Fig. 7). Incubation of 1.0 mM complex in human plasma at 37 °C for 1 h resulted in the formation of up to 18 % of new Mn-containing species for MnL1, up to 5% for MnL2, and up to 5% for MnL3, but no new Mn-containing species were observed when MnL4 was incubated with plasma. Since no new species were observed with the control probe, it is likely that the species formed in plasma are products of reactions with endogenous aldehyde or ketone-containing molecules.

### Distribution and pharmacokinetics in mice

We next used MRI and positron emission tomography (PET) to evaluate the distribution, pharmacokinetics, and whole-body elimination of the probes in naïve C57Bl/6 mice. As shown in Supplementary Fig. 8a, intravenous administration of 0.1 mmol•kg^-1^ of either MnL1, MnL2, MnL3, or MnL4 resulted in an immediate and marked increase in MR signal in the blood pool (aorta). We measured the percentage change in signal intensity (%SI) over time in the aorta, liver, and kidney and the data were fit with a monoexponential decay function to obtain the half-life of the probe in each tissue (Supplementary Fig. 8 b-d). All four probes displayed rapid and almost identical blood clearance with blood half-lives of 7.1 ± 1.2 min (MnL1), 7.0 ± 1.4 min (MnL2), 7.1 ± 1.0 min (MnL3), and 7.2 ± 0.9 min (MnL4). All probes were eliminated from the body via the kidneys. Only minimal and transient liver enhancement was observed and matched the blood pool kinetics. To measure whole-body elimination, we radiolabeled each probe with the Mn-52 isotope and performed PET imaging and ex vivo biodistribution analysis 24 h post-injection of each probe in naïve C57Bl/6 mice. The high sensitivity of PET allows us to detect any trace levels of injected Mn-probe in the body and to distinguish the distribution and elimination of injected probe from endogenous Mn. Mice were injected a dose of MnL1, MnL2, MnL3 or MnL4 (0.1 mmol•kg^-1^) mixed with ^52^MnL1, ^52^MnL2, ^52^MnL3, or ^52^MnL4 respectively (1.0 – 1.2 MBq, Supplementary Fig. 9) by intravenous injection and simultaneous PET-MRI was performed 24 h post-injection (Supplementary Fig. 10a). Finally, the organs were harvested for ex vivo biodistribution analysis. Data are presented as percentage injected dose per gram organ (%ID/g, Supplementary Fig. 10b). At 24 h post-injection, >96% of the injected Mn is eliminated from the mice (98.5% for MnL1, 96.3% for MnL2, 98.6% for MnL3, and 99.0% for MnL4). PET-MRI 24 h post-injection showed that most of the remaining activity was localized in the kidneys for each probe, except for MnL2 where we also observed signal in the skin. The skin is an organ with elevated LOX family expression as a result of the rapid turnover^36, 37^, given the much higher hydrolytic stability of the oxime bond, the higher retained activity from MnL2 is most likely due to binding to intrinsic allysine. An important observation in the PET study for all compounds was the absence of Mn-52 activity in the liver, lymph nodes, bone, brain, and salivary glands of the mice, which would have been indicative of dechelated Mn-52, suggesting that these complexes remain intact in vivo.^38^

### Comparison of noninvasive aldehyde sensing in a mouse model of pulmonary fibrogenesis

We next sought to compare the effects of differing on- and off-rates on noninvasive aldehyde sensing in vivo. We first used a bleomycin-induced lung fibrosis mouse model (BM). A single intratracheal (i.t.) dose of bleomycin results in rapid development of pulmonary fibrosis in mice, and we have previously shown that fibrogenesis and lung allysine levels peak at 14 days post injury^39^. Compared to naïve mice, bleomycin injury resulted in distorted lung architecture and increased cellular infiltration on hematoxylin and eosin (H&E) staining (Fig. 3a), while Masson’s trichrome staining revealed large regions of fibrosis (blue staining of collagen, Fig. 3b). Biochemical quantification of allysine showed a 2-fold increase in the content of this aldehyde in the BM-injured lungs (Fig. 3c) and collagen lung concentration as quantified by hydroxyproline (Fig. 3d), was also significantly elevated 1.3-fold.

**Fig. 3.**
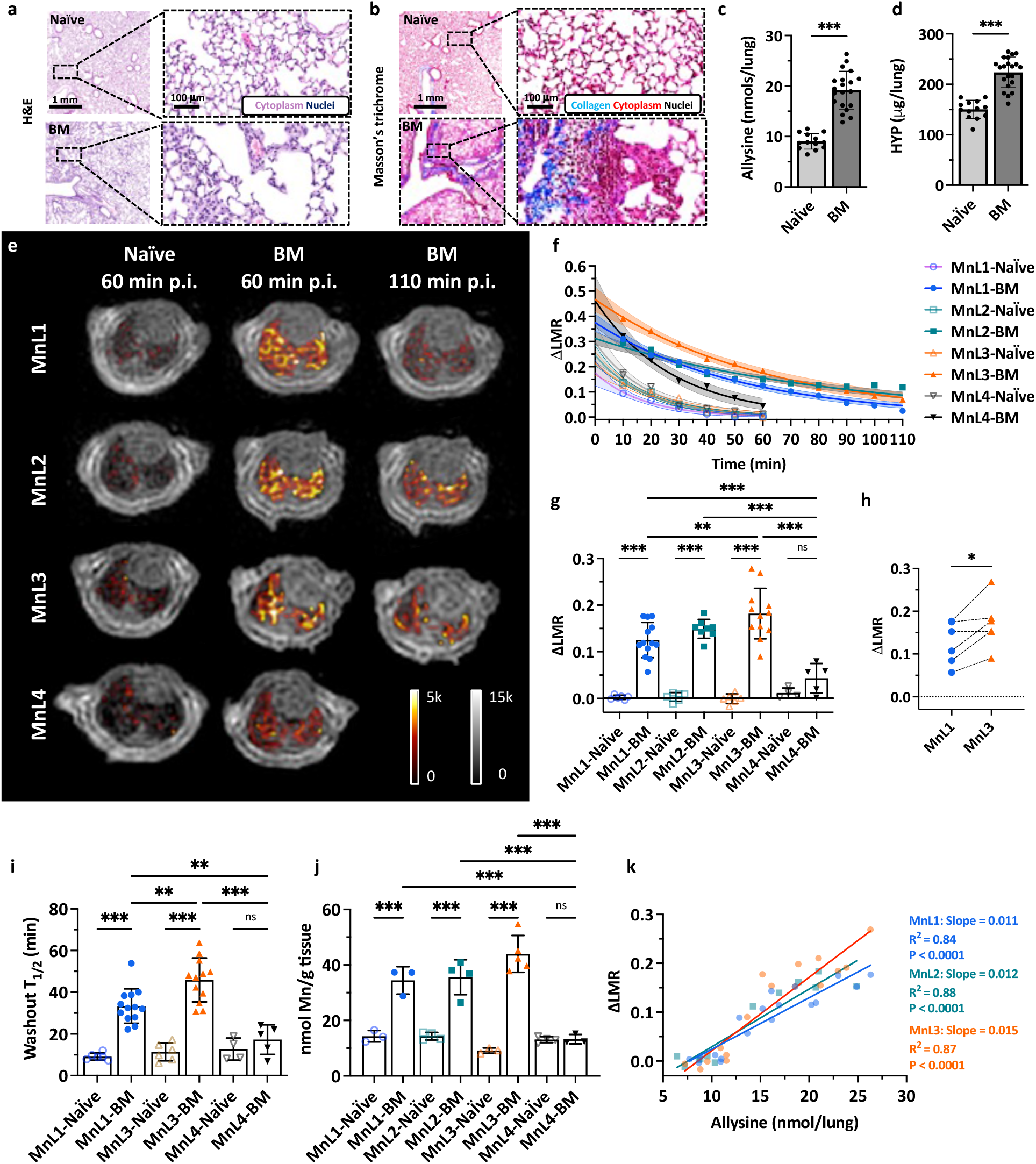
Molecular MRI of bleomycin-induced lung fibrogenesis. Representative H&E (**a**) and Masson’s trichrome (**b**) staining of lung section from naïve and BM mice. Bleomycin-injured lung showed increased tissue density, cellularity, and collagen deposition compared with normal lungs. Lung allysine (**c**) and hydroxyproline (HYP) (**d**) content were significantly increased in BM mice compared with naïve mice. Data presented as mean ± s.d.; n = 13 for naïve mice, n = 20 for BM mice). Statistical analysis was performed using two-tailed unpaired Student’s *t* test, unpaired, ** p < 0.01, *** p < 0.001). **e**, Representative lung enhancement in naïve and BM mice. Coronal UTE images overlaid with false color image of lung enhancement generated by subtraction of the pre-injection UTE image from post-injection UTE image. MnL1, MnL2, and MnL3 produced higher signals in BM mice lungs than in naïve mice and higher signal in MnL4 injected naïve and BM mice at 60 min post-injection. **f**, Change of lung-to-muscle ratio (∆ LMR) as a function of time in naïve and BM mice after injection of MnL1, MnL2, MnL3, and MnL4. Data are presented as mean value with 95% simultaneous confidence bands as shaded regions; Naïve mice: n = 6 for MnL1, MnL2, and MnL3, n = 4 for MnL4; BM mice: n = 13 injected with MnL1, n = 8 injected with MnL2, n = 12 injected with MnL3, and n = 5 injected with MnL4. **g**, Image quantification of ΔLMR in lungs of naïve and BM mice 60 min post-injection of MnL1, MnL2, MnL3, and MnL4. Hydrazine/oxyamine bearing probes exhibited specific lung signal enhancement in the fibrotic lung. Data presented as mean ± s.d. Statistical analysis was performed using one-way ANOVA with Tukey’s post hoc test, ^**^P < 0.01, ^***^P < 0.001, ns not statistically significant). **h**, Pair-wise analysis of ΔLMR in bleomycin-injured mice at 60 min post-injection of MnL1 and MnL3 (n = 6). Statistical analysis was performed using two-tailed paired Student’s *t* test, ^*^P < 0.05. **i**, Washout T_1/2_ of MnL1, MnL3, and MnL4 in naïve and BM mice. MnL3, with higher hydrazone hydrolytic stability, exhibited longer residence time in fibrotic lungs. Data presented as mean ± s.d.; Naïve mice: n = 6 for MnL1, and MnL3, n = 4 for MnL4; BM mice: n = 13 injected with MnL1, n = 12 injected with MnL3, and n = 5 injected with MnL4. Statistical analysis was performed using one-way ANOVA with Tukey’s post hoc test, ^*^P < 0.05, ^***^P < 0.001, ns not statistically significant. **j**, Quantification of Mn content in the left lungs of naïve and BM mice 60 min after injection of MnL1, MnL2, MnL3, and MnL4. Injection of hydrazine/oxyamine-bearing probes produced significantly higher Mn concentrations in the fibrotic lung compared with normal lung. No preferential uptake was observed in naïve or BM mice injected with MnL4. Data are presented as mean ± s.d.; Naïve mice: n = 3 for MnL1, n = 4 for MnL2, n = 3 for MnL3, n = 4 for MnL4; BM mice: n = 3 for MnL1, n = 4 for MnL2, n = 5 for MnL3, n = 3 for MnL4. Statistical analysis was performed using one-way ANOVA with Tukey’s post hoc test, ^*^P < 0.05, ^**^P < 0.01, ^***^P < 0.001, ns not statistically significant. **k**, The MRI lung signal enhancement in naïve and BM mice imaged with MnL1, MnL2, and MnL3 correlates well with allysine content. Compared with MnL1, MnL3 exhibited higher sensitivity in detecting fibrogenesis (slope_MnL1_ = 0.011 vs. slope_MnL3_ = 0.015).

As shown in Fig. 3e, f and Supplementary Fig. 12, from 10 min to 60 min post-injection, lung signal enhancement, expressed as the change in the lung signal to muscle signal ratio (∆ LMR), from all four probes exhibited similar and rapid decreases in naïve mice, demonstrating fast elimination of the probes from normal lungs. In BM mice, the non-binding probe MnL4 produced higher ∆ LMR at 10 min post-injection compared to naïve mice, demonstrating an increased extracellular volume in BM-injured lungs. The MnL4 signal enhancement then exhibited fast decay. In comparison, higher and persistent contrast enhancement was observed in BM mice injected with hydrazine/oxyamine bearing probes, with significantly higher ∆ LMR than in naïve mice and significantly higher ∆ LMR than that of MnL4 in BM mice 60 min post-injection (Fig. 3g). Notably, the signal enhancement of MnL1 and MnL3 in fibrotic lung kept decreasing from 10 min to 110 min post-injection, while the more hydrolytically stable MnL2 showed ∆ LMR reaching a plateau at 60 min post-injection and then remaining stable.

Interestingly, MnL3 exhibited superior signal enhancement compared to MnL1, which was confirmed by pairwise comparison in the same mice to mitigate the inter-animal heterogeneity of this model (Fig. 3h). MnL1 and MnL3 had similar equilibrium constants with aldehyde but different condensation and hydrolysis rates. Therefore, the greater ∆ LMR with MnL3 than MnL1 could be attributed to a slower off-rate with MnL3. We also measured the lung half-life (T_1/2_) of each probe by fitting the change in ∆ LMR with time (Fig. 3i, Supplementary Fig. 13). We found that both MnL1 and MnL3 had significantly longer T_1/2_ in BM mice compared with naïve mice or compared with MnL4 in BM mice. In BM mice, MnL3 exhibited a significantly slower washout than MnL1 (T_1/2_ = 41.5 ± 6.3 min vs. 33.6 ± 5.1 min, P = 0.003), likely due to the greater hydrolytic stability of MnL3. MnL2, which forms a stable oxime bond, showed a very long T_1/2_ value that could not be determined based on the 2-hour imaging study.

Ex vivo determination of lung Mn content at 60 min post-injection of MnL1, MnL2, and MnL3 was significantly elevated in BM mice compared to naïve mice and significantly elevated compared to MnL4 administered to naïve or BM mice, in line with the findings from in vivo MRI (Fig. 3j). The MRI signal enhancement was highly correlated (R^2^ > 0.84) with lung allysine content (Fig. 3k). The slope of ∆ LMR vs allysine concentration was higher for MnL3 (0.015) than for MnL1 (0.011) indicating that MnL3 was 40% more sensitive in detecting fibrogenesis.

The ΔLMR values also significantly correlated with hydroxyproline content in lungs for each of the three aldehyde-targeted probes (Supplementary Fig. 14).

### Noninvasive three-dimensional mapping of fibrogenesis

IPF has a complex phenotype that is manifested by clinical, etiologic, and molecular heterogeneity. In fact, heterogeneity in the radiographic and pathologic features of usual interstitial pneumonia is required for the definitive diagnosis of IPF^40^. However, mapping lung fibrogenesis heterogeneity remains challenging with lung biopsy and other diagnostic modalities. The bleomycin-induced pulmonary fibrogenesis model exhibited strong heterogeneity mimicking the clinical condition. To validate the molecular MRI-based mapping of fibrogenesis with ex vivo histology BM mice were imaged pre- and 40 min post-injection of MnL3, and the %SI enhancement in each pixel was computed to generate a 3D signal enhancement (%SI) map representing the location and extent of fibrotic regions (Fig. 4a, 4b, and Supplementary Video). Following imaging, the injured lung was harvested, fixed, serially sectioned, and stained with Masson’s trichrome for the presence of fibrosis (blue staining, Fig. 4c). We next segmented the blue color in the serial histology images and generated a fibrosis density map for comparison with the corresponding MR %SI map in Fig. 4d; the rest of mice can be seen in Supplementary Fig.15. From the histological representation of fibrosis density, we found the upper-right lobe (dashed line) of this lung exhibited severe fibrotic disease, followed by left lobe (dot-and-dash line), while the lower-right lobe was less fibrotic. Consistently, the corresponding MRI map exhibited a similar pattern of signal enhancement after the injection of the fibrogenesis-sensing probe MnL3. These results demonstrate how MnL3-based molecular MRI can noninvasively track whole lung fibrosis disease activity, and potentially replace, or complement conventional invasive lung histology.

**Fig. 4.**
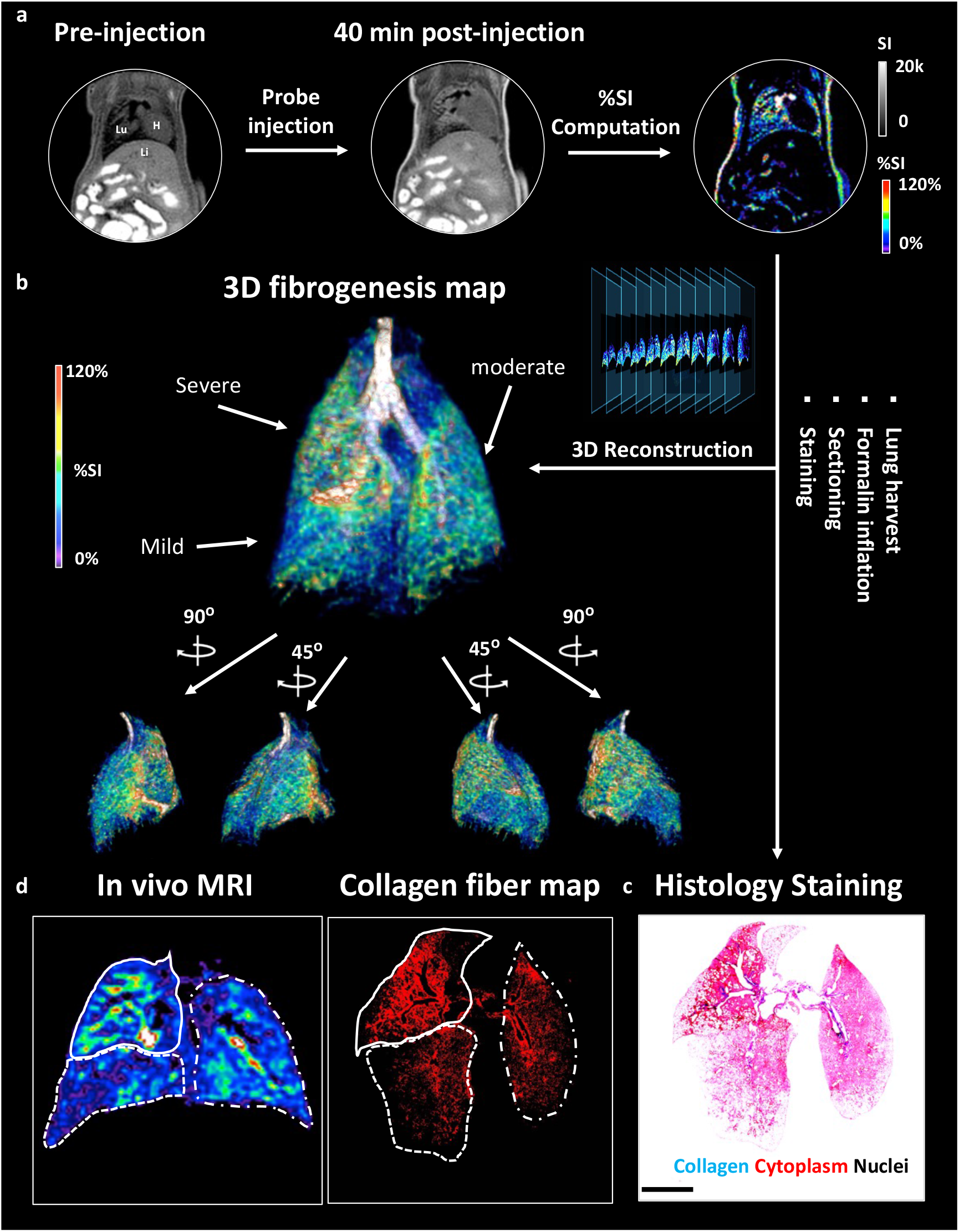
3D mapping of lung fibrogenesis and histology validation. **a**, Representative UTE images of BM mouse pre- and 40 min post-injection of MnL3 were acquired and a pixelwise signal enhancement map was generated (Lu: lung; Li: liver; H: heart). **b**, 3D whole lung fibrogenesis MR map of the fibrotic lung. **c**, Masson’s Trichrome staining. After in vivo imaging, the lung was harvested, fixed, serially sectioned, and stained was performed to validate the MR images. **d**, In vivo MRI signal enhancement exhibited the same fibrotic distribution pattern compared to ex vivo histology.

### MnL1 detection of liver fibrogenesis

After confirming the specificity of MnL1 in targeting fibrogenesis in the lung, we then tested whether MnL1-enhanced MRI could detect liver fibrogenesis in a toxin-induced fibrosis model. Carbon tetrachloride (CCl_4_) is a well-established toxin that when administered repeatedly results in liver fibrosis and ultimately hepatocellular carcinoma^39^. In this study, liver fibrosis was induced in 8-week-old male C57BL/6 mice by oral gavage of 40% CCl_4_ diluted in olive oil for 12 weeks, and vehicle-treated mice were used as controls. Fig. 5a shows that MnL1 administration rapidly produced a significant liver signal enhancement in CCl_4_-treated mice but not in control mice. Change in the liver-to-muscle contrast to noise ratio (∆ CNR) was higher in CCl_4_-treated mice, while the liver only transiently enhanced in vehicle-treated control mice (Fig. 5b). ∆ CNR was 3-fold higher at 20 minutes post-injection, and the area under the ∆ CNR curve was significantly larger in CCl_4_ treated mice than in vehicle-treated mice (Fig. 5c), demonstrating the potential for imaging liver fibrogenesis with MnL1.

**Fig. 5.**
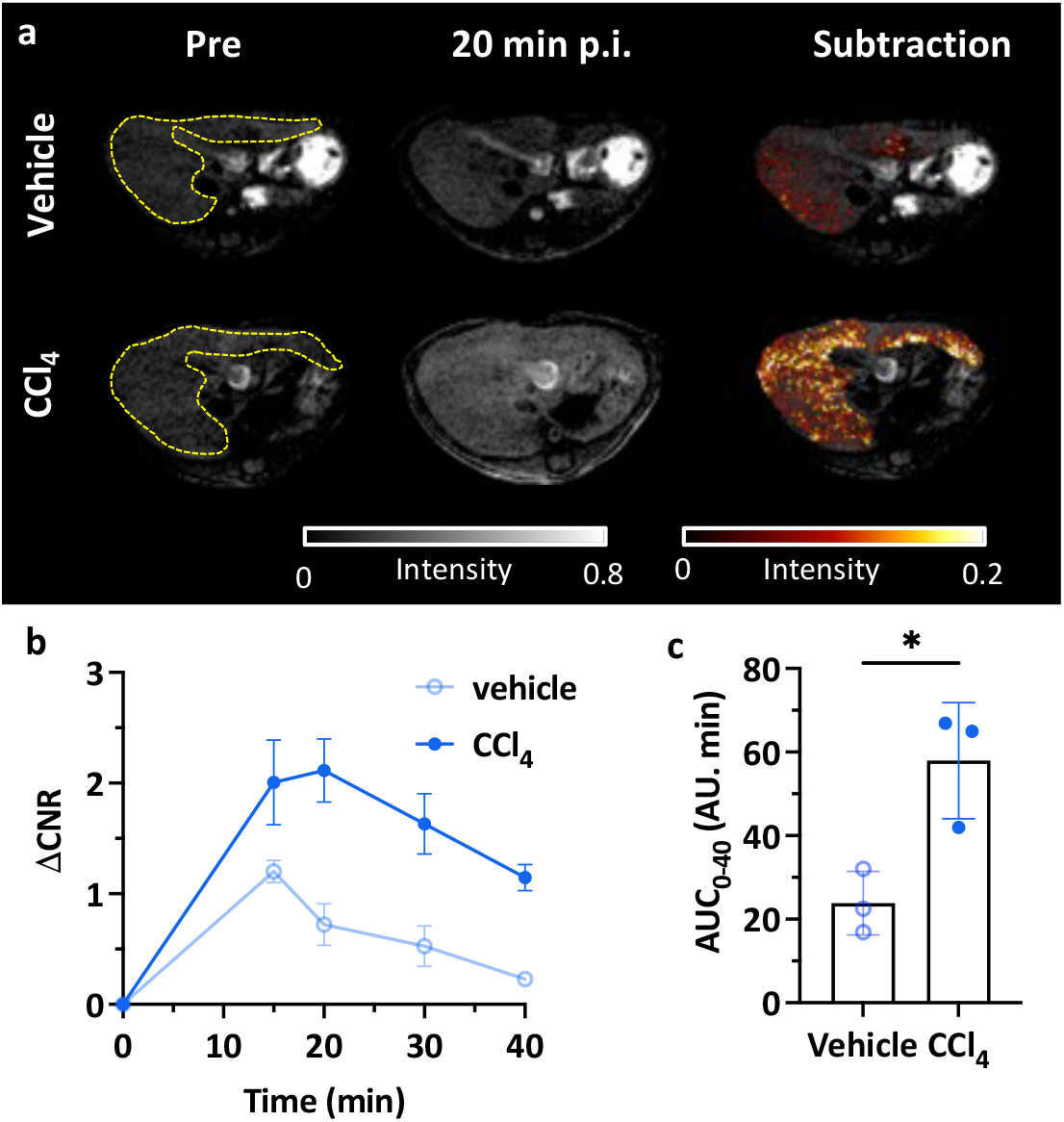
Molecular MRI of hepatic fibrogenesis. **a**, Representative axial MR images showing the livers of mice that received olive oil vehicle (top row) or CCl_4_ (bottom row) for 12 weeks; left: image before MnL1 administration, middle 20 min post-injection of MnL1, and right subtraction of post – pre injection image. **b**, Change in liver-to-muscle contrast to noise ratio (∆ CNR = CNR_Post_ -CNR_Pre_) of vehicle-treated and CCl_4_ mice as a function of time following i.v. injection of MnL1 (Data presented as mean ± s.d.; n = 3 for each group). **c**, Significant difference in the area under the ∆ CNR curve (AUC_0-40_) between vehicle and CCl_4_ mice. Data presented as mean ± s.d.; n = 3 for each group; Statistical analysis was performed using two-tailed unpaired Student’s *t* test, two-tailed unpaired, ^*^P < 0.05.

### Superior hydrolytic stability of the oxime bond enables molecular MR of renal fibrogenesis

Molecular imaging of the kidney is particularly challenging because almost all probes are eliminated to some extent through the kidneys, resulting in a high background signal. To achieve specific molecular imaging of the kidneys, one must wait sufficient time for the nonspecific background renal signal to diminish and the probe must remain bound to its target for this extended period. Here we tested whether the oxyamine probe MnL2 could be used to detect renal fibrosis. We also sought to test whether the 4-fold higher condensation rate constant for MnL2 compared to a previously reported probe GdOA would result in greater signal enhancement in fibrotic kidney.

We used a unilateral renal ischemia-reperfusion injury (IRI) mouse model (Supplementary Fig. 16) where one kidney is clamped and then reperfused. This results in initial inflammation and cell death that then gives way to tissue fibrosis^41^. We evaluated mice at 14 days after IRI and demonstrated fibrosis in both the renal cortex and medulla by histology (Fig. 6a) and by quantifying the biochemical biomarker hydroxyproline (Fig. 6b). Fig. 6a shows obvious collagen deposition in the cortex and medulla of the IRI kidney, but not in the healthy kidney. Consistent with histological staining, hydroxyproline concentrations in the IRI kidney were significantly higher than in the healthy kidney for both cortex and medulla (3.1-times higher than the cortex of healthy kidney; 2.8-times higher than the medulla of the healthy kidney; p < 0.001). Lysyl oxidase like 1 (LOXL 1) and LOXL 2 enzymes are both upregulated in the IRI kidney (Supplementary Fig. 17)

**Fig. 6.**
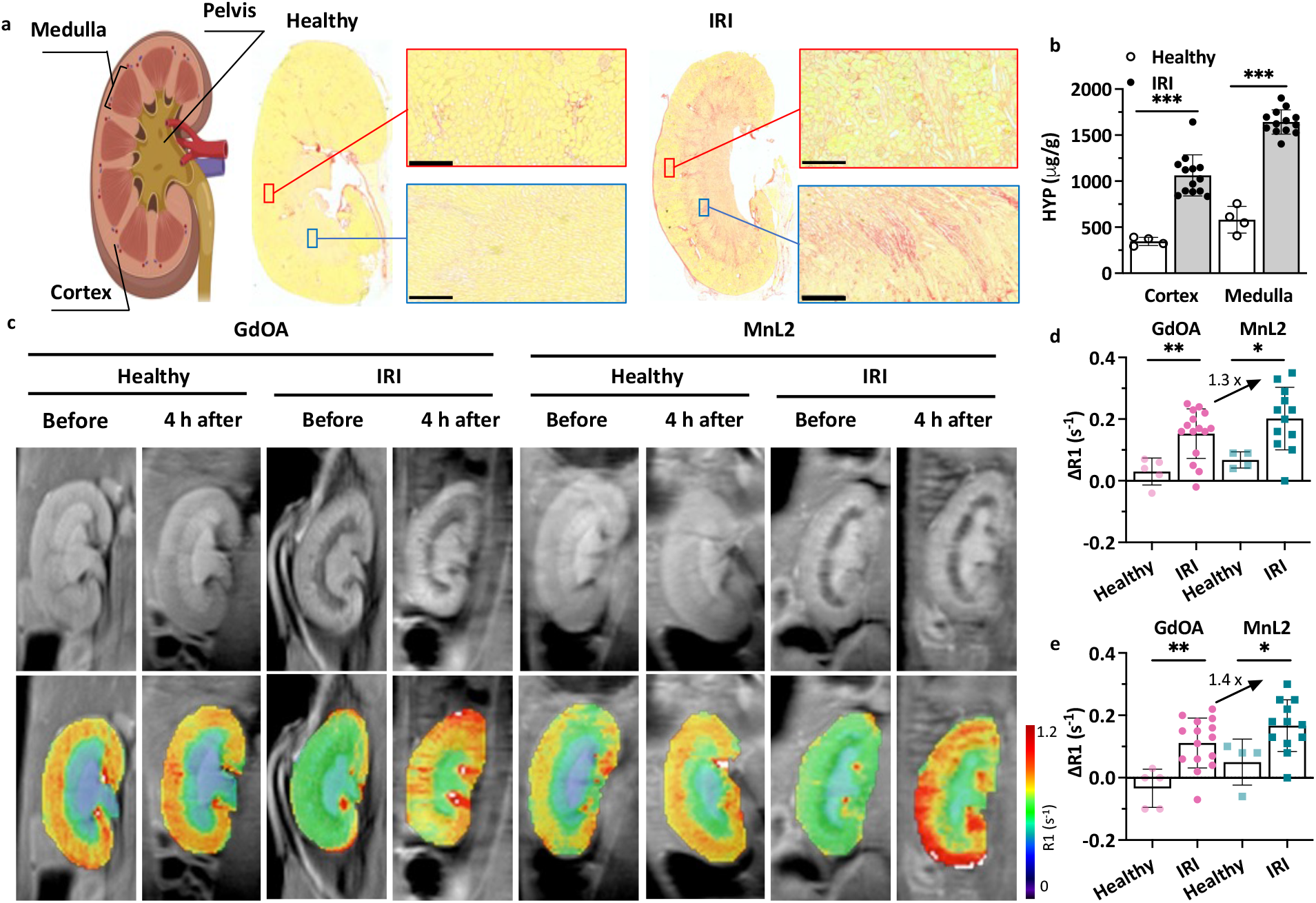
Molecular MRI of renal fibrogenesis. **a**, Left: schematic of anatomy of kidney; middle: Sirius Red stained kidney section from a naïve mouse (healthy); right: Sirius Red stained kidney section from a kidney with ischemia-reperfusion injury (IRI) 14 days prior (collagen: red, normal tissue yellow; scale bar:250 µm). IRI kidney showed increased collagen in the cortex and medulla compared with the healthy kidney. **b**, IRI kidneys showed significantly higher hydroxyproline concentration in the cortex and medulla compared to healthy kidneys. Data are presented as mean ± s.d.; n = 13 for IRI kidney, n = 4 for healthy kidney. Statistical analysis was performed using Student’s *t* test, two-tailed, *** p < 0.001. **c**, Representative mouse MR images showing kidneys of healthy mice and mice 14 days post-IRI. Images are shown before and 4 hours after injection of GdOA or MnL2. Top row shows greyscale kidney anatomy MR image and bottom row shows R1 maps calculated from inversion recovery images. **d**, Quantification of ∆ R1 (R1(4 h post) – R1(pre)) in cortex of healthy and IRI injured mice administered GdOA or MnL2. MnL2 and GdOA exhibited similar low ∆ R1 values in healthy mice, but both probes showed significantly higher ∆ R1 values in the IRI kidney. MnL2 with a higher condensation reaction rate constant compared to GdOA, resulting in higher ∆ R1 than GdOA in the IRI kidney. Data are presented as mean ± s.d.; Naïve healthy mice: n = 5 for GdOA, n = 4 for MnL2; IRI mice: n = 15 for GdOA, n = 11 for MnL2. Statistical analysis was performed using two-tailed unpaired Student’s *t* test, unpaired, ^*^P < 0.05, ^**^P < 0.01. **e**, Quantification of ∆ R1 (R1(4 h post) – R1(pre)) in medulla of healthy and IRI mice administered GdOA or MnL2. MnL2 and GdOA exhibited similar low ∆ R1 values in healthy mice, but both probes showed significantly higher ∆ R1 values in the IRI kidney. MnL2 with a higher condensation reaction rate constant compared to GdOA, resulting in higher ∆ R1 than GdOA in the IRI kidney. Data are presented as mean ± s.d.; Naïve healthy mice: n = 5 for GdOA, n = 4 for MnL2; IRI mice: n = 15 for GdOA, n = 11 for MnL2. Statistical analysis was performed using two-tailed unpaired Student’s *t* test, unpaired, ^*^P < 0.05, ^**^P < 0.01.

We measured kidney T1 values at 9.4 T prior to and 4 hours after injecting the probe. In naïve mice, both probes were effectively eliminated from the kidneys by 4 h post-injection. We computed T1 maps on a pixel basis and then measured the mean R1 (R1 = 1/T1) for cortex and medulla, respectively. We then calculated the change in relaxation rate (∆ R1 = R1_Post_ – R1_Pre_) induced by the probe as a measure of probe concentration in that tissue. In both cortex and medulla of the IRI kidney, ∆ R1 after MnL2 injection was significantly higher than for those regions of healthy kidneys. Using GdOA, we also observed significantly larger ∆ R1 in the injured kidney compared to healthy mice, demonstrating that both oxyamine-containing probes can be used to image renal fibrogenesis. These findings were supported by ex vivo quantification of gadolinium and manganese in these tissues (Supplementary Fig. 18). In comparing the in vivo performance of MnL2 and GdOA, the increase in ∆ R1 was 30% higher for MnL2 in the cortex of the IRI kidney than GdOA and was 40% higher than GdOA in the medulla. This result demonstrated that the 4-fold higher condensation rate constant with MnL2 translated into improved in vivo performance for detection of renal fibrogenesis.

Here we used a unilateral IRI model and thus the contralateral kidney, which did not undergo IRI served as an internal control to the IRI kidney. We performed intra-animal comparisons between ∆ R1 in the contralateral kidney and the IRI kidney for MnL2 and GdOA (Supplementary Fig. 19). For GdOA, ∆ R1 was significantly higher in the IRI medulla compared to the contralateral medulla (p < 0.001), but there was no significant difference in the renal cortex. For MnL2, on the other hand, ∆ R1 was significantly increased in the cortex and medulla of the IRI kidney compared to those regions in the contralateral kidney (p = 0.03 for cortex; p = 0.003 for medulla).

## Discussion

Fibrosis-related diseases present an enormous burden to society. These diseases range from myocardial infarction and hypertrophic cardiomyopathy in the heart^19^, to Crohn’s disease in the intestine^42^, to scleroderma in the skin^43^. IPF is a progressive fibrosing disease of the lung with worse outcomes than many types of cancers^44^. NASH is a chronic liver disease that afflicts approximately 5% of the population in the United States and can lead to cirrhosis, liver failure, primary liver cancer, and/or death^16, 45^. Chronic kidney disease (CKD) is a progressive disease that ultimately leads to renal failure^18^. All these diseases present unmet diagnostic challenges for example in improved early diagnosis, improved prognostication, and monitoring treatment and disease progression. A noninvasive test that could address some or all these needs would be expected to have an enormous impact.

To address this unmet need, we designed molecular probes that target the extracellular aldehyde allysine, which is increased during fibrogenesis. We used MR imaging because of its ability to image any organ in the body with three-dimensional high-resolution images and its operator-independent characteristic. In addition, MR imaging does not require ionizing radiation, which is particularly important for chronic diseases where patients may require multiple imaging exams over their lifetime. We designed small, hydrophilic probes that do not exhibit appreciable nonspecific protein binding to promote rapid renal elimination and minimize hepatobiliary elimination. The lack of hepatobiliary elimination provided a low background in the liver to enable liver imaging applications.

The three aldehyde-targeting Mn(II) probes had similar properties with respect to relaxivity, water exchange kinetics, and kinetic stability of the Mn(II)-PC2A chelate. The relaxivities of MnL1, MnL2, and MnL3 were relatively low (3.1 mM^-1^s^-1^) when measured in PBS but exhibited a 4-fold turn-on in relaxivity when bound to aldehyde modified BSA at 1.4 T. Importantly, the relaxivity increase was negligible in unmodified BSA, indicating that these compounds do not exhibit appreciable nonspecific protein binding. In a Zn^2+^ challenge assay, the Mn probes were about twice as inert to transmetalation as the complex Mn-PyC3A, which has been evaluated in several animal models and is currently undergoing clinical trials. These stable, hydrophilic Mn(II) probes also exhibited similar pharmacokinetic behavior in healthy mice: rapid renal elimination with blood elimination half-lives typical of compounds with no protein binding; exclusive renal elimination; and nearly complete elimination of injected Mn from the body after 24 hours.

The probes differed however in their kinetics of condensation with aldehydes and hydrolysis of the resultant hydrazone or oxime bond. All three probes bind specifically to fibrogenic injured mouse lung in vivo. However, lung MR signal enhancement appears to depend on a balance of on- and off-rates. The α-carboxy hydrazine bearing MnL1 exhibited the highest condensation rate constant, but its fibrotic lung-enhancing properties were significantly inferior to MnL3 which lacked the α-carboxy group. On the other hand, the hydrolysis rate constant for the MnL3 hydrazone was 3-fold lower than the MnL1 hydrazone suggesting that off-rate may be a better predictor of in vivo performance. The α-carboxy oxyamine bearing MnL2 probe had a hydrolysis rate constant that was 3 orders of magnitude smaller than MnL3, but also a 2-fold lower condensation rate constant than MnL3. For the first 30 min after injection, the fibrotic lung signal enhancement with MnL3 was significantly greater than that with MnL2, perhaps indicating that signal enhancement will increase with increasing on-rate provided that the back reaction is relatively slow. Interestingly by 2 h post-injection, the fibrotic lung signal enhancement from MnL3 has diminished as the probe was washed out of the tissue, but the lung enhancement with MnL2 has reached a plateau owing to the slow hydrolysis of the oxime bond.

For applications, these probes offer different options to characterize the fibroproliferative disease. The hydrazine-based probe MnL3 enables specific imaging of fibrogenesis shortly after intravenous administration. The relatively fast hydrolysis of the hydrazone, compared to an oxime, results in the elimination of the probe from the body after the imaging study is performed. We showed here that MnL3-enhanced MRI of the lung allows for three-dimensional characterization of pulmonary fibrogenesis throughout the entire lungs. Pulmonary fibrosis can affect any part of the lungs – upper or lower, right or left, subpleural or mid lung – and methods that assess the whole lungs are needed. Clinical techniques like high-resolution computed tomography (HRCT) can report on the distribution of advanced fibrosis throughout the whole lungs, but HRCT does not report disease activity^46^. Lung biopsy provides rich data but only samples a single, very small region of the lung that may not be reflective of the whole lung and is a procedure that carries the risk of complications and mortality^47^. Endobronchial optical coherence tomography provides high-resolution images of specific lung regions but again does not report on disease activity^48^.

We also showed that MnL1-enhanced MR could detect and quantify liver fibrogenesis within 20 minutes post-injection. In chronic liver diseases like NASH, ultrasound elastography or MR elastography or serum biomarker panels have shown value in detecting the presence of advanced fibrosis^49^. However, these methods are insensitive to detecting fibrosis at earlier stages and do not assess fibrotic disease activity^50^. Prior work in animal models showed that imaging liver fibrogenesis may enable the detection of the earliest stages of liver fibrosis and that imaging disease activity can provide an early readout of response to therapy^12^.

Extending molecular imaging of fibrosis/fibrogenesis to the kidneys is particularly challenging because of the high nonspecific renal signal enhancement as the probe clears through the kidneys. Furthermore, many molecular probes, especially peptide-based probes, are retained in the kidneys. In the MnL2-enhanced MR study of pulmonary fibrosis, we found that the lung signal reached a plateau and that we could not estimate a washout rate of MnL2 from the fibrotic lung over two hours period because the lung signal was not changing. This finding led us to test whether this in vivo hydrolytic stability could be exploited for delayed imaging of the kidney. Our results showed that MnL2 could indeed be used to specifically detect renal fibrosis in IRI model by imaging the mice before and 4 hours post intravenous injection of MnL2. Using T1 mapping, we could accurately quantify fibrogenesis throughout the kidney and segment disease within the cortex and medulla. We also showed that the faster on-rate with MnL2 compared to GdOA resulted in greater MR signal enhancement in diseased tissue, again demonstrating the importance of optimizing the reactivity of these probes toward aldehydes.

In conclusion, this study provides insight into the design of molecular MR probes for noninvasive sensing of extracellular aldehydes and how they might be deployed to characterize human diseases. These probes have a high potential for clinical translation given the straightforward syntheses that are amenable to scale-up. While we did not assess the safety and toxicology of the probes, we note that their lack of nonspecific protein binding, lack of cellular uptake in vivo, rapid renal elimination, and whole-body elimination of injected Mn are all favorable properties that should limit the potential for adverse effects. We showed that the probes exhibit a 4-fold increase in relaxivity upon binding to protein containing aldehyde at 1.4 T, near the frequently used clinical field strength of 1.5 T. Our animal experiments were performed at 4.7 T and 9.4 T where the turn-on effect is much lower, e.g. 2-fold increase at 4.7 T. Thus, we expect the signal enhancement in clinical scanners to be higher than what we measured here. While we demonstrated the utility of these probes in mouse models of lung, liver, and kidney disease, such probes would likely prove valuable in other fibroproliferative diseases, including the heart, intestines, blood vessels, skin, etc.

## Methods

**The details of chemical syntheses, chromatography methods, in vitro characterizations of probes, animals and experimental groups, and post-imaging ex vivo analysis protocols are provided in the Supplementary Information**.

### General synthetic procedures

The general synthetic procedure is shown below using MnL3 as an example. Detailed synthetic procedures of all the other compounds are available in the supplementary information.

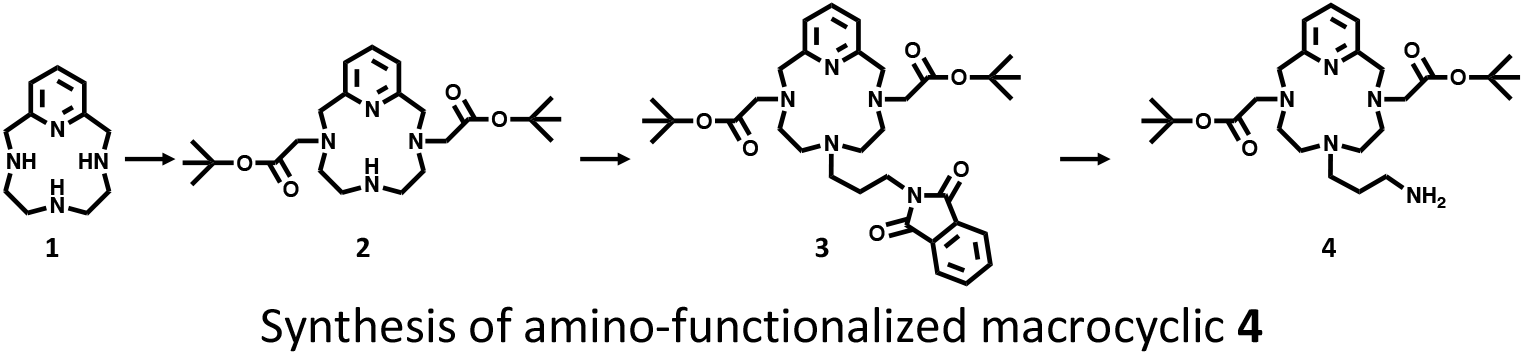

### Regioselective alkylation of the compound 2

The starting material **1** (1.03 g, 5.0 mmol) was dissolved in a mixed solvent containing 50 mL DI water and 25 mL 1,4-dioxane and the pH was adjusted to 8.5 with concentrated HCl. To this solution, *tert*-butyl bromoacetate (1.37 g, 7.0 mmol) dissolved in 1,4-dioxane (25 mL) was added dropwise. The addition of more *tert*-butyl bromoacetate was repeated twice (2×0.23 g, 0.48 mmol) after 12 h and 24 h and the pH was adjusted to 8.5 with 1N NaOH. Reaction completion was monitored by LC-MS (method 1). The reaction mixture was extracted with CHCl_3_ (3×50 mL) and the combined organic phase was evaporated to dry under reduced pressure. The obtained residue was purified *via* flash chromatography (C18 column, method 7 with A and B as solvents) to give **2** as a pale brown oil (1.37 g, 63%). ^1^H NMR (500 MHz, Chloroform-*d*) δ 7.63 – 7.49 (m, 1H), 7.01 (dd, *J* = 7.7, 1.7 Hz, 2H), 3.96 (s, 4H), 3.50 (s, 4H), 3.36 (t, *J* = 5.5 Hz, 4H), 2.98 (t, *J* = 6.1 Hz, 4H), 1.44 (s, 18H). ^13^C NMR (126 MHz, Chloroform-*d*) δ 171.20, 160.12, 137.70, 120.42, 81.64, 57.72, 57.45, 51.83, 46.24, 28.29. LC-MS (method 1): t_R_=3.35 min, m/z = 435.3 [M+H]^+^; calcd: 435.3.

### N7-alkylation of macrocyclic 2

Compound **2** (0.86 g, 2.0 mmol) and K_2_CO_3_ (0.54 g, 4.0 mmol) were suspended in dry ACN (40 mL) and N-(3-Bromopropyl)phthalimide (0.8 g, 3.0 mmol) in dry ACN (20 mL) was added dropwise. The suspension was brought to reflux under argon for 12h. Following removal of the precipitate by filtration, the reaction mixture was concentrated under reduced pressure and then purified via chromatography (method 7 with mobile phase A and B) to give **3** as a yellow oil (1.18g, 95%). ^1^H NMR (500 MHz, Chloroform-*d*) δ 7.82 (dt, *J* = 5.6, 2.9 Hz, 2H), 7.71 (dt, *J* = 5.7, 2.8 Hz, 2H), 7.59 (t, *J* = 7.7 Hz, 1H), 7.02 (d, *J* = 7.6 Hz, 2H), 3.97 (s, 4H), 3.79 (q, *J* = 6.3 Hz, 2H), 3.69 – 3.49 (m, 4H), 3.39 (s, 6H), 3.16 (s, 4H), 2.16 (dt, *J* = 15.3, 7.0 Hz, 2H), 1.40 (s, 18H). ^13^C NMR (126 MHz, Chloroform-*d*) δ 169.95, 168.31, 159.29, 138.18, 134.40, 131.86,123.58, 120.91, 82.10, 58.54, 58.00, 52.39, 50.20, 45.79, 35.59, 28.20, 21.84. LC-MS (method 1): t_R_=3.32 min, m/z = 622.3 [M+H]^+^; calcd: 622.4.

### Liberation of primary amine

Compound **3** (0.62g, 1 mmol) and hydrazine hydrate (2.5 g, 50 mmol) were dissolved in ethanol (20 mL) and the solution was stirred for 1h at 45℃. This mixture was diluted with acetonitrile (20 mL), and the precipitate was removed by filtration and the solvent was removed by reduced pressure. The obtained crude product was purified *via* flash chromatography (C18 column, method 7 with mobile phase A and B) to give **4** as a yellow oil (0.41g, 85%). ^1^H NMR (500 MHz, Chloroform-*d*) δ 7.60 (t, *J* = 7.7 Hz, 1H), 7.03 (d, *J* = 7.7 Hz, 2H), 3.93 (s, 4H), 3.60 – 3.39 (m, 6H), 3.38 – 3.26 (m, 4H), 3.23 – 3.12 (m, 4H), 3.12 – 3.00 (m, 2H), 2.22 (dd, *J* = 10.2, 6.5 Hz, 2H), 1.41 (s, 18H). ^13^C NMR (126 MHz, Chloroform-*d*) δ 170.51, 159.56, 138.25, 120.87, 81.99, 58.24, 57.97, 51.92, 50.18, 43.69, 36.89, 28.08, 19.56. LC-MS (method 1): t_R_=2.742 min, m/z = 492.5 [M+H]^+^; calcd: 492.4. ^1^H NMR (500 MHz, Chloroform-*d*) δ 7.61 (t, *J* = 7.7Hz, 1H), 7.04 (d, *J* = 7.7 Hz, 2H), 4.08 – 3.86 (m, 4H), 3.54 – 3.30 (m, 12H), 3.28 – 2.96 (m, 6H), 2.05 (m, 2H), 1.44 (s, 18H), 1.41 (s, 9H). ^13^C NMR (126 MHz, Chloroform-*d*) δ 171.66, 170.46, 170.24, 159.59, 159.44, 138.46, 121.02, 82.29, 58.39, 57.95, 54.76, 51.87, 50.31, 44.55, 36.20, 28.34, 28.18, 21.56. LC-MS (method 1): t_R_=3.27 min, m/z = 664.4 [M+H]^+^; calcd: 664.6.

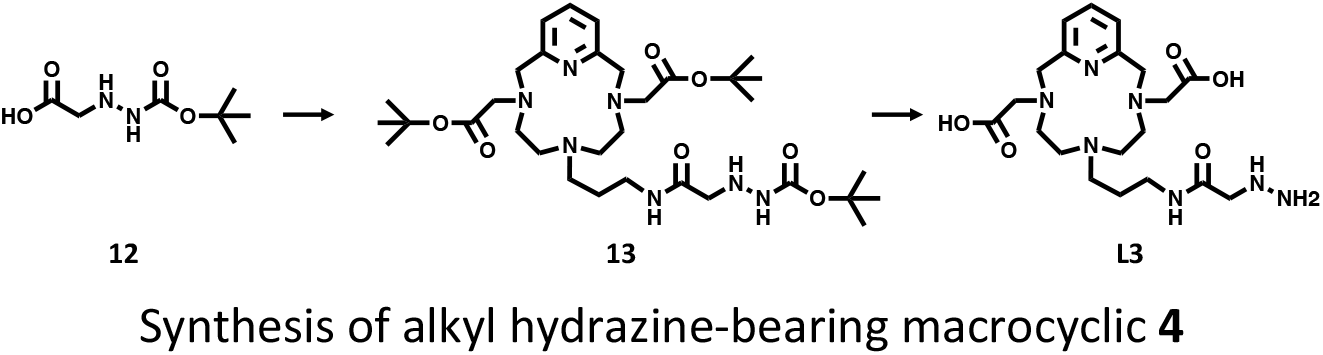

### Coupling between amino functionalized macrocyclic and targeting moiety

Compound **4** (0.25 g, 0.5 mmol), compound **12** (0.14 g, 0.75 mmol), and *N, N*-diisopropylethylamine (0.13 g, 1.0 mmol) were dissolved in dry acetonitrile (20 mL), and HATU (0.29 g, 0.75 mmol) in dry acetonitrile (10 mL) was added. The reaction mixture was stirred at room temperature for 45 min. After evaporating under reduced pressure, the obtained oil was redissolved in dichloromethane and extracted with 10% citric acid aqueous solution and brine. The organic layer was dried (Na_2_SO_4_) and concentrated under reduced pressure. The crude product was purified *via* flash chromatography (C18 column, method 7 with mobile phase A and B to give **13** as a yellow oil (0.26 g, 79%). ^1^H NMR (500 MHz, Chloroform-*d*) δ 7.61 (t, *J* = 7.7Hz, 1H), 7.04 (d, *J* = 7.7 Hz, 2H), 4.08 – 3.86 (m, 4H), 3.54 – 3.30 (m, 12H), 3.28 – 2.96 (m, 6H), 2.05 (m, 2H), 1.44 (s, 18H), 1.41 (s, 9H). ^13^C NMR (126 MHz, Chloroform-*d*) δ 171.66, 170.46, 170.24, 159.59, 159.44, 138.46, 121.02, 82.29, 58.39, 57.95, 54.76, 51.87, 50.31, 44.55, 36.20, 28.34, 28.18, 21.56. LC-MS (method 1): t_R_=3.27 min, m/z = 664.4 [M+H]^+^; calcd: 664.6.

### TFA-promoted deprotection

To compound **13** (132.7 mg, 0.2 mmol) in dichloromethane (5 mL) cooled to 0 ℃, was added anisole (0.4 mL, 2 mL/mmol, cation scavenger) followed by the slow addition of trifluoroacetic acid (5 mL) and the mixture was stirred at 0 ℃ for 1 h followed by room temperature for 4 h. The solvent and trifluoroacetic acid were gently evaporated under reduced pressure and the resulting oily residue was dissolved in water (5 mL) and the organic byproducts were pipetted away with diethyl ether (3 × 5 mL). The aqueous layer was freeze-dried, and the obtained crude product was purified by preparative HPLC (C18 column, method 8 with A and B as solvent) to give **L3** as a white solid (69.5 mg, 77%). ^1^H NMR (500 MHz, D_2_O) δ 8.25 (t, *J* = 7.9 Hz, 1H), 7.63 (d, *J* = 7.9 Hz, 2H), 4.39 (s, 4H), 3.77 (s, 4H), 3.62 (s, 2H), 3.33 – 2.86 (m, 12H), 2.00 – 1.76 (m, 2H). ^13^C NMR (126 MHz, D_2_O) δ 175.13, 169.90, 152.05, 146.46, 124.00, 57.76, 56.67, 52.85, 51.74, 51.69, 50.96, 50.90, 50.59, 36.34, 22.66. LC-MS (method 2): t_R_=2.39 min, m/z = 452.2 [M+H]^+^; calcd: 452.3.

### MnL3 complex preparation

**L3** (0.02 mmol) was dissolved in water (2.0 mL), and the pH was adjusted to 6.5 by 0.1 M aqueous NaOH. Then solid MnCl_2_•4H_2_O (0.02 mmol) was added under stirring. The pH of the solution dropped to about 3.5, and the pH was adjusted back to 5.0 by addition of 1.5 M aqueous NaOAc. Completion of the chelation was confirmed by LC-MS. Any excess unchelated Mn^2+^ was removed by Chelex 100 resin (pH 6.5**). MnL3** was then lyophilized to a powdery solid. **MnL3**: LC-MS (method 5): t_R_=8.02 min, m/z [M + H]^+^ calcd for [C_20_H_32_N_7_O_5_Mn]^+^, 505.2; found 505.2, purity = 94% (250 nm UV detector).

### Kinetics of hydrazone/oxime formation

The condensation reaction rate constants for reactions of MnL1, MnL2, MnL3, and GdOA with butyraldehyde were monitored spectrophotometrically via the increase of the 220/230 nm absorbance. All reactions were carried out at 25 °C in PBS. A series of experiments where the concentrations of probes were varied (0.04 mM, 0.06 mM, 0.08 mM, and 0.10 mM) to estimate the rate law. The probe was reacted with 1 mM butyraldehyde to ensure that the reaction proceeded under pseudo-first order with respect to the probe. For a second-order reaction, the observed rate constant (*k*_*obs*_) is the product of the second-order rate constant (*k*) and [butyraldehyde]. In this regard, the rate law and a corresponding *k* can be estimated by plotting reaction velocity as a function of [MnL1], [MnL2], [MnL3], and [GdOA]. All rate measurements are summarized in Supplementary Figure 1.

### Kinetics of hydrazone/oxime hydrolysis

Hydrazones and oximes were prepared by mixing solution of each probe (1 mL, 1 mM, PBS, pH 7.4) with butyraldehyde (14.4 mg, 200 mmol), and the mixture was rocked gently at room temperature for 10 min, frozen with liquid N_2_, and lyophilized for 12 h. The obtained product was redissolved in 1 mL water. The HPLC trace of the resultant hydrazone or oxime was first measured. Then, a 37% stock solution of formaldehyde was added (final concentration: 200 mM) to trap the liberated nitrogen base, and HPLC traces were obtained at desired time point. The extent of hydrolysis was quantified by monitoring the area under the hydrazone/oxime peak on HPLC trace, and the reactions were allowed to proceed to > 95% completion. Hydrolysis of hydrazone/oxime is a first-order reaction and follows the rate law. Then the extent of hydrolysis was plotted as a function of time to calculate the hydrolysis rate constant.

### Relaxivity measurements

Relaxivity measurements were performed on a Bruker mq60 Minispec, 1.4 T, NMR spectrometer at 37 °C. Longitudinal (T1) relaxation times were measured via an inversion recovery experiment using 10 inversion times of duration ranging between 0.05 x T1 and 5 x T1; Longitudinal relaxivity (r1) was determined from the slope of a plot of 1/T1 vs. [Mn]. Metal ion concentrations were determined by ICP-MS.

MnL1, MnL2, MnL3, MnL4, and GdOA (concentration range: 0.05 mM – 0.2 mM) were incubated with BSA-Ald (21.9 mg mL^-1^, 0.6 mM aldehyde. Please see Supplementary Information for preparation details) or BSA (21.9 mg mL^-1^, 0.06 mM aldehyde) at 37°C for 3h, and then relaxivities were measured. Solutions of MnL1, MnL2, MnL3, MnL4, and GdOA (concentration range: 0.1 mM – 1.0 mM) in PBS were run in parallel as standard controls.

After incubation of each probe with BSA and BSA-Ald, the unbound Mn and Gd probe was separated from the protein-bound probe by ultrafiltration (10 KDa cut-off PLCC cellulosic membrane, 10 min, 10,000 RPM). Following separation, the concentrated protein residue was diluted to a total volume of 300 μL in PBS, and then relaxivities were immediately measured.

### Animals and experimental groups

C57Bl/6 adult male mice at 10-12 weeks of age were purchased from Charles River Laboratories, Wilmington MA. Animals were housed under a 12 h light/12 h dark cycle and provided with water and food ad libitum. The ambient temperature and relative humidity were 20–25 °C and 50–60%, respectively. All experiments and procedures were performed in accordance with the National Institutes of Health’s “Guide for the Care and Use of Laboratory Animals” and were approved by the Massachusetts General Hospital Institutional Animal Care and Use Committee. During MRI and PET scanning, animals were anesthetized with 1–2% isoflurane and air/oxygen mixture to maintain a constant respiration rate (60 ± 10 breaths per minute), kept warm by a thermal pad, and monitored by a small animal physiological monitoring system (SA Instruments Inc., Stony Brook NY).

A total of 117 C57Bl/6 mice were included in this study.

(1)19 mice were included to study probe whole-body elimination.

(2)57 mice were included to study lung fibrosis.

(3)6 mice were included to study liver fibrosis.

(4)35 mice were included to study kidney fibrosis

### MR and PET image acquisition

Briefly, the following MRI systems were utilized: (1) 4.7 T Bruker Biospec scanner with PET insert using custom-built volume coil, operating software: ParaVision 7.0 for MRI and ParaVision 360 for PET (2) 9.4T Bruker scanner (Bruker BioSpin) with a quadruple volume transmit/receiving coil, operating software: ParaVision 5.0. Pre-injection and post-injection MR images were acquired for comparison. Probes were administrated via intravenous bolus injection at 0.1 mmol•kg^-1^ body weight, with the bolus volume of injection never exceeding 100 μL. MR images were acquired with the following sequences and parameters: Probe pharmacokinetics: A series of T1-weighted 3D Fast Low Angle Shot (FLASH, TR/TE/FA=10 ms/ 2.5 ms/ 12°, 0.4 mm isotropic spatial resolution, field of view (FOV) 60 mm × 50 mm, one average, acquisition time = 2 min) were acquired before and dynamically after intravenous injection of the probe.

Whole-body elimination: Static 30 min PET measurement was performed 24 ± 0.5 h post administration of ^52/nat^Mn complexes. The images were reconstructed using MLEM (Maximum Likelihood Expectation Maximization) with 75 iterations and 0.75mm cubic voxels. During PET image acquisition, T1-weighted 3D FLASH (TR/TE/FA=21 ms/ 3 ms/ 12°, 0.25 mm isotropic spatial resolution, field of view (FOV) 60 × 85 × 50 mm^3^, one average, acquisition time = 2.2 min) images were acquired for the anatomical information.

Lung imaging: 3D ultrashort time to echo (3D-UTE, TR/TE/FA=4 ms/ 11.75 μs/ 16°, 0.6 mm isotropic spatial resolution, field of view (FOV) 75 mm × 75 mm, one average, acquisition time = 3 min) images were acquired prior to and dynamically post injection of 0.1 mmol•kg^-1^ probe; 2D Rapid Acquisition with Relaxation Enhancement (2D-RARE, TR/TE/FA=1.5 s/ 8 ms/ 180°, resolution = 0.3 × 0.3 × 1 mm^3^, field of view (FOV) 60 mm × 50 mm, four averages, acquisition time = 3 min) and T1-weighted 3D FLASH (TR/TE/FA=10 ms/ 2.5 ms/ 30°, 0.6 mm isotropic spatial resolution, field of view (FOV) 60 mm × 50 mm, one average, acquisition time = 2 min) were acquired for atomical information.

Liver imaging: A series of T1-weighted 3D FLASH (TR/TE/FA=10 ms/ 2.5 ms/ 30°, 0.6 mm isotropic spatial resolution, field of view (FOV) 60 mm × 50 mm, one average, acquisition time = 2 min) were acquired prior to and dynamically after intravenous injection of MnL1.

Kidney imaging: T1-map was acquired using a RARE sequence with various inversion times (TI): 9 TIs ranging from 7 ms to 5 s, TR 5 s, TE 7.26 ms, respiratory gated, FOV 30mm x 30mm, dimension 128 × 128, single slice with 0.75 mm slice thickness. Animals were first scanned for pre-injection T1-map. Mice were then removed from the magnet for probe injection and allowed to wake up and recover in a rodent cage. Four hours after probe injection, animals were returned to the scanner for post-injection T1-map acquisition.

### MR and PET images analysis

MR and PET images were analyzed and shown using Horos (version 3.3.5) and Amide (version 1.0.1) with the following protocol: Lung image analysis: 2D-RARE images and 3D-FLASH images were used to define regions of interest (ROIs) in the lung that excluded vessels and airways and these ROIs were copied to the UTE images for signal quantification. A total of 6 lung ROIs were defined on axial image slices spanning both lungs to obtain signal intensity (SI); ROIs in dorsal muscle in each slice were also defined as reference. We averaged the lung-to-muscle ratio (LMR, equation 1) from the 6 slices to calculate changes in LMR (ΔLMR, equation 2).

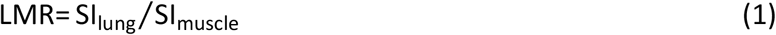

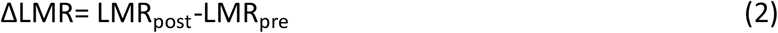

To measure the half-life of each probe in the lung, ∆ LMR values were monoexponentially fit to the following equation:

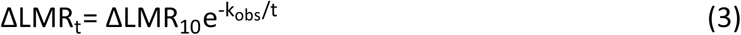

Where ∆ LMR_t_, and ∆ LMR_10_ are the changes in lung-to-muscle ratio at time *t*, and at 10 min post-injection of probe. Equation (3) can be simplified to equation (4):

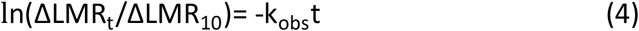

The washout T_1/2_ can be computed by equation (5):

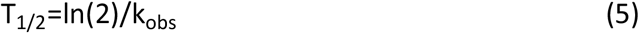

Liver image analysis: A ROI was manually traced encompassing the liver parenchyma while avoiding major blood vessels. A second ROI was placed on the dorsal muscle visible in the same image slice to quantify the signal intensity in the muscle for comparison. Seven ROIs were placed in the field of view without any tissue (air) to measure the variation in background signal. 6 axial slices per mouse across the entire liver were analyzed in this fashion. Contrast to noise ratio (CNR, equation 6) was calculated by measuring the difference in signal intensity (SI) between liver and muscle and normalized to the standard deviation of the signal in the air.

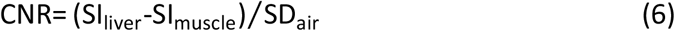

ΔCNR was calculated by subtracting CNR measured prior to probe injection (CNR_Pre_) from CNR measured after injection (CNR_Post_).

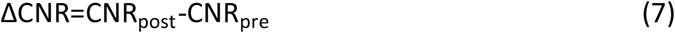

Kidney image analysis: ROIs for renal cortex and medulla were segmented based on signal contrast of the inversion recovery images. Pixel-by-pixel T1 maps over kidneys were obtained by fitting MR signals to TIs (equation 8):

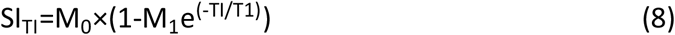

T1 values for a given ROI were obtained by averaging the T1 values for individual pixels within that ROI, and then R1 values were calculated (equation 9).

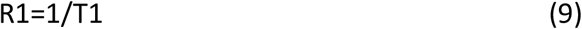

Changes in R1 were calculated by equation 10:

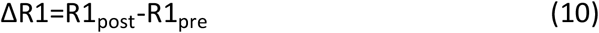

### Statistical analysis

GraphPad Prism (version 9.1.1) was used for statistical analyses. Comparisons were performed with unpaired Student’s t-test to analyze differences between two groups, paired Student’s t-test to analyze intraanimal differences, or one-way ANOVA with Tukey’s post hoc comparison to analyze differences among three or more groups. Values of P < 0.05 were considered statistically significant. Error bars are s.d. or 95% confidence interval, as indicated in the figure legends and main text. Sample sizes can be found in the figure legends and were chosen based on previous experience. Animals were fully randomized.

## Acknowledgments

This work was supported by grants from the National Institutes of Health to P.C. (DK104302, DK121789, HL154125, OD028499, OD032138, OD025234, and OD023503), to E.A. (K01HL155237), to R.K. (K08HL140175) and by a Sponsored Research Agreement with Janssen. We thank Alana Ross for assistance with the biodistribution study.

## Author contributions

All the authors discussed the results and commented on the manuscript. H.M. and P.C. conceived the study, generated the hypotheses, and designed the experiments. H.M. synthesized all the probes. H.M., E.A., and H.W. carried out the in vitro analyses. I.A., R.K., C.Z., M.D., and M.S. generated the animal models and collected the tissue samples. H.M., I.Y.Z., Y.I.C., N.J.R., J.W.W., and B.F.M. performed the in vivo imaging and analyzed the data.

Y.I.C. performed the MRI %SI map generation. M.D., A.T.B., S.Z., and P.P. performed the tissue allysine, hydroxyproline, and ICP analysis. K.K.T., L.P.H., R.G.V., and M.H. advised on animal models and study design. H.M. and P.C. wrote the manuscript. Correspondence and requests for materials should be addressed to P.C.

## Ethics declarations

### Competing interests

H.M., E.A., and P.C. are inventors of a filed patent based on the work here (Molecular probes for in vivo detection of aldehydes. PCT/US2022/072310). P.C. has equity in and is a consultant to Collagen Medical LLC, has equity in Reveal Pharmaceuticals Inc., and has research support from Transcode Therapeutics, Pliant Therapeutics, Takeda, and Janssen. L.P.H. reports grants from Boehringer Ingelheim and has received personal consulting fees from Boehringer Ingelheim, Pliant Therapeutics, Bioclinica, and Biogen Idec. R.G.V., and M.K.H. are employed by Janssen. The other authors declare that they have no competing interests.

