## Supplementary Figures and Methods for "Tailored chemical reactivity probes for systemic imaging of aldehydes in fibroproliferative diseases"

### Experimental Section

#### Materials and instruments

All chemical reagents were purchased from commercial suppliers and used as received unless stated otherwise. MilliQ water was used for every reaction unless otherwise noted. NMR spectra were recorded on a JEOL ECZ 500R 11.7 T NMR system equipped with a 5 mm broadband probe (1H: 499.81 MHz, 13C: 125.68 MHz). The following abbreviations were used in ^1^H NMR: s = singlet; d = doublet; t = triplet; q = quartet; m = multiplet. UV-vis spectra were recorded on a SpectraMax M2 spectrophotometer using quartz cuvettes with a 1 cm path length. Metal ion concentrations were determined using an Agilent 8800-QQQ ICP-MS system. All samples with diluted with 0.1% Triton X-100 in 5% nitric acid. A linear calibration curve for metal ranging from 1.0 ppb to 1000 ppb in diluent was generated daily for the quantification. HPLC-MS analysis was carried out on an Agilent 1260 system (UV detection at 220, 254 and 280 nm) coupled to an Agilent Technologies 6130 Quadrupole MS system using the following method:

Mobile Phases: A: H_2_O (+ 0.1% formic acid, v/v), B: CH_3_CN (+ 0.1% formic acid, v/v), C: H_2_O (+10 mM NH_4_OAc), D: 90% CH_3_CN + 10% solvent C.

Method 1: Column, Phenomenex Luna C18, 5μm, 100 x 2 mm, flow rate: 0.7 mL/min

| min | %A | %B |
| --- | --- | --- |
| 0 | 95 | 5 |
| 3 | 5 | 95 |
| 4.5 | 5 | 95 |
| 5 | 95 | 5 |
| 7 | 95 | 5 |

Method 2: Column, Restek Ultra AQ C18, 5 μm, 100 x 4.6 mm column, flow rate: 1.0 mL/min

| min | %A | %B |
| --- | --- | --- |
| 0 | 95 | 5 |
| 0.5 | 95 | 5 |
| 8 | 5 | 95 |
| 8.5 | 95 | 5 |
| 10 | 95 | 5 |

Method 3: Column, Restek Ultra AQ C18, 5 μm, 100 x 4.6 mm column, flow rate: 1.0 mL/min

| min | %C | %D |
| --- | --- | --- |
| 0 | 95 | 5 |
| 1 | 95 | 5 |
| 11 | 5 | 95 |
| 13 | 5 | 95 |
| 14 | 95 | 5 |
| 16 | 95 | 5 |

Method 4: Column, Restek Ultra AQ C18, 5 μm, 250 x 4.6 mm column, flow rate: 1.0 mL/min

| min | %C | %D |
| --- | --- | --- |
| 0 | 95 | 5 |
| 1 | 95 | 5 |
| 11 | 5 | 95 |
| 13 | 5 | 95 |
| 14 | 95 | 5 |
| 16 | 95 | 5 |

Method 5: Column, Xbridge C18, 5 μm, 150 x 4.6 mm, flow rate: 1.0 mL/min

| min | %C | %D |
| --- | --- | --- |
| 0 | 95 | 5 |
| 1 | 95 | 5 |
| 7 | 5 | 95 |
| 8 | 5 | 95 |
| 9 | 95 | 5 |
| 10 | 95 | 5 |

Method 6: Column, Xbridge C18, 5 μm, 150 x 4.6 mm, flow rate: 1.0 mL/min

| min | %C | %D |
| --- | --- | --- |
| 0 | 95 | 5 |
| 1 | 95 | 5 |
| 8 | 50 | 50 |
| 9 | 5 | 95 |
| 10 | 5 | 95 |
| 11 | 95 | 5 |
| 13 | 95 | 5 |

Large-scale reverse-phase purifications were carried out on a Teledyne ISCO Combiflash system with UV-Vis detection using the following methods:

Mobile Phases: A: H_2_O (+ 0.1% formic acid, v/v) B: CH_3_CN (+ 0.1% formic acid, v/v)

Method 7: Column: 150 g C18 gold, flow rate: 85 mL/min

| Column volume | %A | %B |
| --- | --- | --- |
| 0 | 95 | 5 |
| 3 | 95 | 5 |
| 15 | 10 | 90 |
| 20 | 10 | 90 |
| 23 | 95 | 5 |
| 25 | 95 | 5 |

Preparative reversed-phase HPLC with UV detection at 220, 254 and 280 nm was performed using the following methods:

Mobile Phases: A: H_2_O (+ 0.05% TFA, v/v) B: CH_3_CN (+ 0.05% TFA, v/v)

Method 8: Column: Restek, UltraAqueous C18, 5 μm, 250x21.2 mm, flow rate: 15 mL/min

| min | %A | %B |
| --- | --- | --- |
| 0 | 95 | 5 |
| 1 | 95 | 5 |
| 12 | 90 | 10 |
| 22 | 40 | 60 |
| 25 | 5 | 95 |
| 28 | 95 | 5 |
| 30 | 95 | 5 |

Radio-HPLC analysis was carried out on an Agilent 1100 Series HPLC unit with a Carroll/Ramsey radiation detector with a silicon PIN photodiode using following methods:

Mobile Phases: A: H_2_O (+ 10 mM NaOAc, pH 4.5) B: 90% CH_3_CN + 10% solvent A

Method 9: Column: Kromasil C18, 5 μm, 150 x 4.6 mm, flow rate: 1.0 mL/min

| min | %A | %B |
| --- | --- | --- |
| 0 | 95 | 5 |
| 1 | 95 | 5 |
| 8 | 5 | 95 |
| 9 | 5 | 95 |
| 10 | 95 | 5 |
| 11 | 95 | 5 |

#### In vitro measurements

*Kinetics of Hydrazone formation.* The molar extinction coefficients (ε_probe_) of MnL1 and MnL3 were calculated at 220 nm for MnL1, MnL2, and GdOA, and at 230 nm for MnL3 by measuring the absorbance as a function of concertation (Supplementary Table 1). The molar extinction coefficients (ε_hydrazone_) of the reaction products of each probe with butyraldehyde (MnL-BAld or GdOA-BAld) were calculated by comparing the HPLC trace of the probe alone (1 mM, using Method 4) with the trace of 1 mM probe reacted with 200 equivalents of butyraldehyde added to force the condensation reaction to completion. The ε_hydrazone_ values were calculated by eq 1.

$\varepsilon_{\mathrm{hydrazone}}=\varepsilon_{\mathrm{probe}}\times\frac{\mathrm{AUC}_{\mathrm{hydrazone}}}{\mathrm{AUC}_{\mathrm{probe}}}$	（1）

where AUC_probe_ refers to the peak area of the pure probe and AUC_hydrazone_ is the peak area of the hydrazone product. The *ε* values are summarized in Supplementary Table 1.

The general reaction scheme is shown below,

MnL (or GdOA) $\to$ MnL-Ald (or GdOA-Ald)

At the beginning of reaction (t=0) the concentration of probe is *a_0,_* and that of hydrazone is zero. If after time *t* the concentration of hydrazone is x, that remaining probe is *a_0_-x*. The rate of change of the concentration of Z is *dx/dt*, and thus for a first-order reaction

$\frac{d[x]}{dt}=k_{obs}\left( a_{0}-x \right)$	（2）

Separation of the variables gives

$\frac{dx}{a_{0}-x}=k_{obs}dt$	（3）

and integration gives

$\ln\left( a_{0}-x \right)=-k_{obs}t-I$	（4）

Where *I* is the constant of integration, *k_obs_* is observed rate constant.

The concentration of hydrazone can be estimated by the following equations:

The absorbance of MnL or GdOA is

$A_{MnL}=\varepsilon_{MnL}C_{MnL}l$	（5）

The absorbance of the reaction mixture is

$A_{mix}=\varepsilon_{MnL}{(C}_{MnL}-C_{MnL-Ald})l+\varepsilon_{MnL-BAld}C_{MnL-BAld}l$	（6）

After subtraction between eq. 6 and eq. 5

$${\Delta A=A}_{mix}-A_{MnL}=\varepsilon_{MnL}{(C}_{MnL}-C_{MnL-BAld})l+\varepsilon_{MnL-Ald}C_{MnL-BAld}l-\varepsilon_{MnL}C_{MnL}l$$

$\Delta A=(\varepsilon_{MnL-BAld}{-\varepsilon_{MnL})C}_{MnL-Ald}l$	（7）

For second order reaction, the observed rate constant (*k_obs_*) is the product of the second order rate constant (*k*) and [butyraldehyde]. In this regard, the rate law and a corresponding *k* can be estimated by plotting reaction velocity as a function of [MnL1], [MnL2], [MnL3], and [GdOA]. All rate measurements are summarized in supplementary Figure 1.

*Kinetics of hydrazone hydrolysis.*

The extent of hydrolysis was quantitated by eq 8.

$\%\mathrm{hydrolysis}=100\frac{A_{0}-A}{A_{0}}$	(8)

where A_0_ is the HPLC peak area of the hydrazone before addition of formaldehyde, A is the peak area of the hydrazone/oxime at time *t*. eq 9.

$$\frac{d[y]}{dt}=k_{obs,H}(b_{0}-y)$$

(9)

where the concentration of hydrazone at *t* = 0 is b_0_ and the concentration of hydrolysis product is zero. After time *t* the concentration of hydrolysis product is y, and the concentration of remaining hydrazone is b_0_-y. Separation of the variables followed by integration gives eq 10.

$\ln\left( b_{0}-y \right)=-k_{obs,H}t-I_{H}$	(10)

The constant *I_H_* can be evaluated using the boundary condition that x=0 when t=0; hence$-lnb_{0}=I_{H}$. Insertion of this into eq. S10 leads to eq. 11.

$ln\frac{b_{0}}{b_{0}-y}=kt$	(11)

This eq. can also be written as eq. S12

$b_{0}-y=b_{0}e^{-kt}$	(12)

Or eq. S13

$y=b_{0}(1-e^{-kt})$	(13)

When the reaction reaches a plateau value of y*_max_* at equilibrium, $y_{max}=b_{0}$, and insertion of this into eq. S13 leads to eq. S14

$y=y_{max}(1-e^{-kt})$	(14)

Dividing by the initial concentration of hydrazone gives eq 15:

$Y=Y_{max}(1-e^{-kt})$	(15)

Where Y is the %hydrolysis, *t* is time, *k* is the first-order rate constant, and Y_max_ is the %hydrolysis at t=$\infty$. All kinetics measurements were carried out in triplicate.

*Preparation of oxidized BSA, BSA-Ald:*

To a solution of bovine serum albumin (BSA) (100 mg) dissolved in PBS (2 mL, pH 7.4), sodium ascorbate (20 mg), ferric chloride (120 µL, 10 mM) and 20 µL H_2_O_2_ (30% w) were added. The reaction was stirred at 37 °C for 24 h, and sodium ascorbate (20 mg) was added repeatedly every 8 h. After the reaction, protein was purified using PD-10 Sephadex G25 desalting columns (GE Healthcare), eluted with PBS. Protein concentration was assessed using the ‘BCA Protein Assay Kit’ (Thermo Scientific). BSA-Ald had an aldehyde concentration of 4 aldehyde/protein. The allysine-modified BSA (BSA-Ald) had a concentration of 43.8 mg/mL, while unmodified BSA had a concentration of 50.0 mg/mL. Protein carbonyl concentration was determined by ‘Protein Carbonyl Content Assay Kit’ (Sigma-Aldrich). BSA-Ald had an aldehyde concentration of 1.2 mM aldehyde, while unmodified BSA had an aldehyde concentration of 0.1 mM.

*Transmetallation.* The kinetic inertness of Mn complexes (1 mM) was assessed at 37 °C in MES buffer (0.1 M, pH 6.0), in the presence of 25-fold excess of the exchanging Zn^2+^ (25 mM) to provide pseudo-first order conditions. The transmetallation was monitored by measuring the water proton longitudinal relaxation rate (R_1_=1/T_1_) change of the complex solution at 1.4 T, 37°C. The paramagnetic R_1_^P^ relaxation rate is obtained by subtracting the diamagnetic contribution of the water relaxation (0.34 s^-1^) from the observed relaxation rate. The change in paramagnetic relation rate was calculated by eq. S16:

$\%R_{1}^{P}(t)=\frac{R_{1}^{P}\left( t \right)-R_{1}^{P}(0)}{R_{1}^{P}(0)}\times100$	(16)

where R_1_^P^(0), and R_1_^P^(t) are the R_1_^P^ of 1 mM Mn complex, and R_1_^P^ of mixture of 1 mM Mn complex with 25 mM Zn^2+^ at time *t*. Experimental data were fit to a first-order exponential equation to obtain the half-lifetime of complex dissociation.

Animals and experimental groups**:**

Whole-body elimination assay:

(1) 5 mice were injected with ^52/nat^MnL1.
(2) 5 mice were injected with ^52/nat^MnL2.
(3) 5 mice were injected with ^52/nat^MnL3.
(4) 4 mice were injected with ^52/nat^MnL4.

All the mice were administrated with a cocktail dose of ^52/nat^Mn complexes (^52^Mn complex: 1.0-1.2 MBq, ^nat^Mn complexes: 0.1 mmol•kg^-1^ body weight). After injection, mice were housed in cages with bedding changed every 8 h to prevent reabsorption of excreted dose.

Lung fibrosis imaging:

(1) Intratracheal bleomycin instillation, imaged after 2 weeks (n=26): each mouse was imaged with probe 1 and then imaged again 24 hours later with probe 2. Probe 1 and probe 2 were randomly chosen from MnL1, MnL2, MnL3, and MnL4
(2) For pair-wise comparison, an additional cohort of bleomycin-injured mice (n=6) were imaged randomly with MnL1 or MnL3 on day 14 and imaged again with the other probe on day 15.
(3) Age-matched naïve mice were used as healthy control (n=14): each mouse was imaged with probe 1 and then imaged again 24 hours later with probe 2. Probe 1 and probe 2 were randomly chosen from MnL1, MnL2, MnL3, and MnL4
(4) For the fibrogenesis 3D mapping study, 3 bleomycin-injured mice were imaged with MnL3.

All animals were euthanized after imaging, and various organs were harvested for biochemical and/or histological analysis.

Liver fibrosis imaging:

(1) 3 mice that received CCl_4_ in olive oil for 12 weeks to induce liver fibrosis.
(2) 3 mice that received a vehicle for 12 weeks were used as control.

Renal fibrosis imaging:

(1) 36 mice underwent IRI surgery to induce kidney fibrosis; 15 were injected with GdOA, 11 were injected with MnL2.
(2) 9 age-matched healthy naïve mice were used as healthy control, 4 were injected with MnL2 and 5 were injected with GdOA.

All mice were euthanized after imaging. The right and left kidneys were harvested for further analysis.

*Bleomycin-induced lung fibrosis model:* mice were first anesthetized with ketamine/xylazine (100/10 mg•kg^-1^; intraperitoneal injection). Then a single dose of bleomycin (Fresenius Kabi, Lake Zurich, Il), prepared in sterile PBS, was intratracheally injected at 1.1 units per kg body weight (50 μL total volume). Mice were imaged 14- and 15-days after bleomycin administration. Buprenorphine (0.1 mg•kg^-1^) for analgesia was subcutaneously injected 30 min before anesthesia and every 8-12 hours after bleomycin administration for 3 days. Age-matched naïve mice were used as healthy control.

*CCl_4_-induced liver fibrosis model:* mice were treated with CCl_4_ in olive oil for 12 weeks (0.1 ml of 20% CCl_4_ in olive oil the first week, 30% the second week, and 40% from weeks 3-12) by oral gavage, 2-3 times per week. Control mice were fed vehicle (olive oil) only.

*Mice renal IRI model:* Mice were anesthetized with ketamine/xylazine mixture (ketamine: 100 mg•kg^-1^, xylazine: 10 mg•kg^-1^). Animals were then placed prone while maintaining rectal temperature strictly at 37°C with a feedback-regulated heating pad. The skin incision was performed on the left lower flank while following strict aseptic procedures. After accessing the retroperitoneal space, the left kidney and the left renal artery - vein was identified. Ischemia was induced by applying a micro serrefine clip onto the renal artery and vein. Successful ischemia was visually confirmed by a gradual uniform darkening of the kidney. The clamp was removed 26 minutes later, and a rapid change in kidney color from a dark maroon to a healthy dark pink verified successful reperfusion. The skin was closed using surgical staples, and animals were returned to their home cages. Analgesia was provided by buprenorphine (0.1 mg•kg^-1^, SC, bid for three days starting 1 hour before IRI surgery). Animals were imaged 14 days after IRI. Age-matched naïve mice were used as control.

#### Post-imaging ex vivo tissue analysis

*Ex vivo whole-body elimination assay:* After PET-MR imaging, all animals were sacrificed and the following tissues were collected: brain, blood, heart, lung, liver, kidney, stomach, spleen, intestine, muscle, tail, bone, and skin. All the tissues were weighed, and radioactivity in each tissue was measured on a gamma counter (Wizard2Auto Gamma, PerkinElmer). Probe distribution is expressed as percent of injected dose per gram of tissue (%ID/g) for all the tissues collected.

*Histological analysis*: For the lung fibrosis study, mice were euthanized at appropriate time points followed by exsanguination, and the diaphragm was carefully cut open without damaging the lungs. A small incision was made in the proximal region of the trachea and the lungs were inflated with 4% formaldehyde using a 30G needle. The inflated lungs were dissected and fixed with 4% formaldehyde for 24h at room temperature. For the kidney fibrosis study, the left and right kidneys were harvested and fixed with 4% formaldehyde for 24h at room temperature. Fixed tissues were embedded in paraffin, sectioned at 5 μm, then hematoxylin and eosin (H&E), Sirius red, and Masson’s trichrome staining were carried out using standard techniques (Harvard Medical School Pathology Core).

*Left lung allysine assay.* Left lungs were digested in 1 mL of 6 M HCl, 650 μL of H_2_O, and 100 μL of 4M fluorescein, and the allysine was reacted with 40 mg sodium 2-naphthol-6-sulfonate. Dysprosium (40 µL of 10,000 ppb) was added as an internal standard for tissue metal ion concentration measurement (see below). These samples were digested overnight in an oil bath at 105 °C. The digest was then weighted, and a 200 μL aliquot was taken and weighed for pH balancing. The aliquot was balanced to a pH of 2 with 6 M NaOH and 1 M HCl, then weighed again to determine the dilution factor. The samples were then split into 2 separate HPLC vials and diluted in equal amounts of DMSO. One of these samples was then spiked with a known amount of labeled allysine for later analysis. The samples were then subjected to HPLC following published method^1^.

*Right lung and kidney hydroxyproline assay:* The hydroxyproline assay was performed as previously reported^2^. For the lung fibrosis study, right lungs were used for hydroxyproline assay. For the kidney fibrosis study, the renal cortices and medullae were separated and prepared by weight.

*Mn and Gd content in tissues:* Metal ion concentrations in the tissue were determined by ICP-MS, as previously described^3^. Calibration standards for Mn and Gd were used, and dysprosium was used as an internal standard.

### Synthetic Section

Compounds **1**, **5**, **9** and **12** were synthesized according to Ref^4-6^.

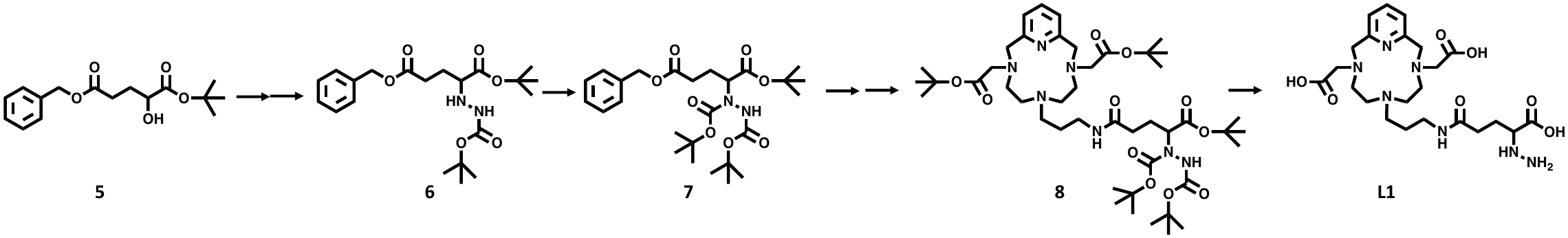

**Synthesis of 6.** A solution of alcohol **5** (2.94 g, 10 mmol) in dry dichloromethane (50 mL) was treated with 2,6-lutidine (3.21g, 30 mmol) and Tf_2_O (3.38 g, 12 mmol) successively at 0 ℃. After 1 h at this temperature, the obtained mixture of compound **5** was added dropwise to tert-Butyl carbazate (6.6 g, 50 mmol) in 20 mL dry dichloromethane at 0 ℃. After stirring for 1 h, the solution was washed with aqueous 10% citric acid solution and brine. The organic layer was dried (Na_2_SO_4_) and concentrated under reduced pressure. The crude product was purified via flash chromatography to give **6** as a yellow oil (method 7, 1.76 g, 43%). ^1^H NMR (500 MHz, Chloroform-*d*) δ 7.40 – 7.29 (m, 5H), 6.15 (s, 1H), 5.11 (s, 2H), 4.21 (s, 1H), 3.53 (s, 1H), 2.52 (t, J = 7.4 Hz, 2H), 2.15 – 2.04 (m, 1H), 1.92 (dd, J = 14.3, 7.2 Hz, 1H), 1.45 (s, 9H), 1.43 (s, 9H). ^13^C NMR (126 MHz, Chloroform-*d*) δ 173.32, 171.89, 156.33, 136.04, 128.58, 128.23, 81.98, 80.41, 66.35, 62.45, 30.31, 28.38, 28.12, 25.16. LC-MS (method 1): t_R_=4.50 min, m/z = 409.3 [M+H]^+^; calcd: 409.2.

**Synthesis of 7.** Compound **6** (0.82 g, 2 mmol) was combined with di-*tert*-butyl decarbonate (0.88 g, 4 mmol) and dissolved in 30 mL dichloromethane and a solution of 4-dimethylaminopyridine (24 mg, 0.2 mmol) in dichloromethane was added dropwise. The obtained mixture was stirred overnight at room temperature. After extraction with 10% citric acid aqueous solution and brine, the organic layer was dried (Na_2_SO_4_) and concentrated under reduced pressure. The crude product was purified via flash chromatography to give **7** as a yellow oil (method 7, 0.88 g, 87%). ^1^H NMR (500 MHz, Chloroform-*d*) δ 7.39 – 7.28 (m, 5H), 5.10 (d, *J* = 3.1 Hz, 2H), 3.53 (t, *J* = 5.5 Hz, 1H), 2.74 (m, 1H), 2.36 (m, 1H), 2.01 (m, 2H), 1.47 (s, 18H), 1.44 (s, 9H). ^13^C NMR (126 MHz, Chloroform-*d*) δ 173.11, 170.67, 152.16, 136.05, 128.61, 128.29, 128.26, 83.79, 82.33, 66.31, 62.54, 29.81, 28.06, 25.83. LC-MS (method 1): t_R_=4.90 min, m/z = 509.3 [M+H]^+^; calcd: 509.3.

**Synthesis of 8.** Compound **7** (0.51 g, 1 mmol) was added to a slurry of palladium on carbon (50% water, 50 mg) in methanol (20 mL). The mixture was subject to two cycles of vacuum and hydrogen purge and then stirred under an atmosphere of hydrogen for 12 h at room temperature. After evacuating the system of hydrogen, celite was added and the slurry filtered through a methanol-wet bed of celite. The filtrate was concentrated under reduced pressure to give a colorless oil. The obtained colorless oil, compound **4** (0.25 g, 0.5 mmol) and *N, N*-diisopropylethylamine (0.13 g, 1.0 mmol) were dissolved in dry acetonitrile (20 mL), and O-(7-Aza-1H-benzotriazol-1-yl)-N, N, N', N'-tetramethyluronium hexafluorophosphate (HATU, 0.29 g, 0.75 mmol) in dry acetonitrile (10 mL) was added. The reaction mixture was stirred at room temperature for 45 min. After evaporating under reduced pressure, the obtained oil was redissolved in dichloromethane and extracted with 10% citric acid aqueous solution and brine. The organic layer was dried (Na_2_SO_4_) and concentrated under reduced pressure. The crude product was purified via flash chromatography to give **9** as a yellow oil (method 7, 0.88 g, 87%). ^1^H NMR (500 MHz, Chloroform-*d*) δ 7.56 (t, *J* = 7.7 Hz, 1H), 7.00 (d, *J* = 7.7 Hz, 2H), 5.09 (s, 1H), 3.91 (s, 4H), 3.45 – 3.03 (m, 18H), 2.51 – 2.38 (m, 1H), 2.34 – 2.15 (m, 1H), 2.07 – 1.78 (m, 4H), 1.43 (s, 18H), 1.39 (s, 27H). ^13^C NMR (126 MHz, Chloroform-*d*) δ 152.47, 139.12, 121.82, 83.54, 82.51, 62.97, 58.05, 52.18, 50.43, 36.72, 31.84, 29.71, 28.18, 28.08, 27.02, 22.46. LC-MS (method 1): t_R_=3.67 min, m/z = 892.5 [M+H]^+^; calcd: 892.6.

**Synthesis of L1.** To compound **9** (178.3 mg, 0.2 mmol) in dichloromethane (5 mL) cooled to 0 ℃, was added anisole (0.4 mL, 2 mL/mmol)(cation scavenger) followed by the slow addition of trifluoroacetic acid (5 mL) and the mixture was stirred at 0 ℃ for 1 h followed by room temperature for 4 h. The solvent and trifluoroacetic acid were gently evaporated under reduced pressure. The resulting oily residue was dissolved in water (5 mL) and the organic byproducts were pipetted away with diethyl ether (3 x 5 mL). The aqueous layer was freeze-dried, and the obtained crude product was purified by preparative HPLC to give **L1** as a white solid (method 8, 88.9 mg, 85%). ^1^H NMR (500 MHz, D_2_O) δ 8.16 (m, 1H), 7.56 (d, *J* = 7.6 Hz, 2H), 4.39 (s, 4H), 3.75 (s, 4H), 3.62 (t, *J* = 6.5 Hz, 1H), 3.26 – 2.92 (m, 12H), 2.26 (m, 2H), 1.95 (m, 1H), 1.88 (m, 1H), 1.84 – 1.74 (m, 2H). ^13^C NMR (126 MHz, D_2_O) δ 175.02, 174.71, 171.25, 150.65, 141.00, 122.98, 62.50, 59.13, 57.44, 52.62, 51.90, 50.15, 37.09, 31.74, 25.17, 24.08. LC-MS (method 2): t_R_=5.58 min, m/z = 524.2 [M+H]^+^; calcd: 524.3.

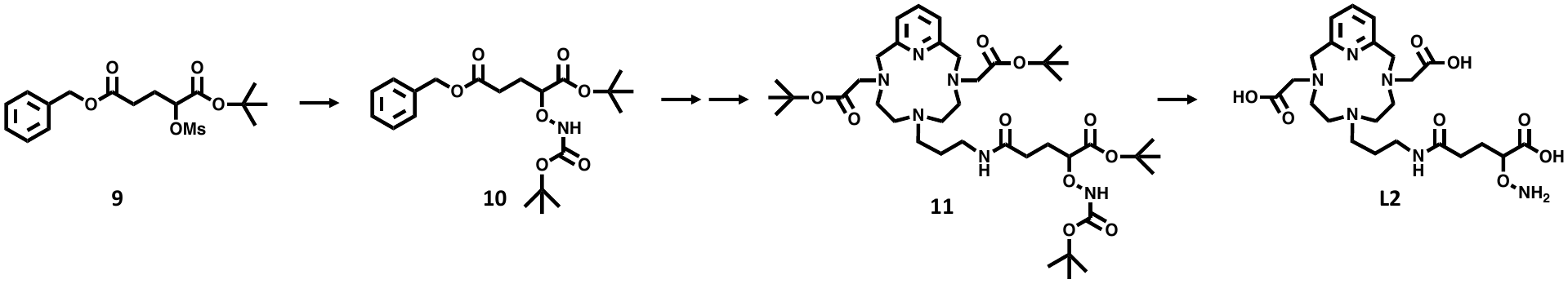

**Synthesis of 10:** To a stirring solution of N-Boc hydroxylamine (1.99 g, 15 mmol) in dimethylformamide (20 mL) was added DBU (3.04 g, 20 mmol) in dimethylformamide (1 mL). Then compound 9 (3.72 g, 10 mmol) was added dropwise. The reaction was allowed to stir for 20 h at 50 ℃. The reaction solution was then concentrated, and the residue was purified via flash chromatography to give Compound **10** as a yellow oil (method 7, 2.17 g, 53%). ^1^H NMR (500 MHz, CDCl3) δ 7.40 – 7.29 (m, 5H), 5.11 (s, 2H), 4.24 (dd, J = 9.2, 3.8 Hz, 1H), 2.63 (m, 2H), 2.22 – 2.12 (m, 1H), 2.07 – 1.89 (m, 1H), 1.45 (s, 9H), 1.46 (s, 9H). 13C NMR (126 MHz, CDCl3) δ 172.83, 170.48, 156.37, 136.00, 128.63, 128.27, 82.76 -82.00, 66.42, 29.99, 28.27 -28.17, 26.00. LC-MS (method 1): t_R_=4.45 min, m/z = 432.1 [M+Na]^+^; calcd: 432.2.

**Synthesis of 11:** compound **10** (0.41 g, 1.0 mmol) was added to a slurry of palladium on carbon (50% water, 50 mg) in methanol (10 mL). The mixture was purged twice with hydrogen and then stirred at room temperature under argon for 12 h. Celite was added into the reaction mixture and the slurry was filtered through a celite bed preliminary wetted with methanol. The filtrate was concentrated under reduced pressure to give a colorless oil of product which was used for the next step without further purification. The obtained colorless oil, compound **4** (0.25 g, 0.5 mmol), and DIPEA (0.13 g, 1.0 mmol) were dissolved in dry acetonitrile (20 mL), and HATU (0.29 g, 0.75 mmol) in acetonitrile (10 mL) was added. The reaction mixture was stirred at room temperature for 45 min. After the solvent was removed under reduced pressure, the obtained oil was redissolved in dichloromethane and extract with 10% citric acid aqueous solution and brine. The organic layer was dried over Na_2_SO_4_ and concentrated under reduced pressure. The crude product was purified via flash chromatography to give compound **11** as a yellow oil (method 7, 0.29 g, 73%). ^1^H NMR (500 MHz, CDCl_3_) δ 7.60 (t, J = 7.6 Hz, 1H), 7.03 (d, J = 7.7 Hz, 2H), 4.19 (dd, J = 9.5, 3.7 Hz, 1H), 3.97 (s, 4H), 3.58 – 3.09 (m, 16H), 2.57 – 2.46 (m, 1H), 2.44 – 2.34 (m, 1H), 2.20 – 2.08 (m, 1H), 2.06 – 1.91 (m, 3H), 1.44 (s, 9H), 1.43 (s, 27H). 13C NMR (126 MHz, CDCl3) δ 173.55, 170.28, 167.01, 159.54, 138.16, 120.86, 82.95, 82.13, 81.78, 58.39, 58.01, 52.35, 50.41, 45.41, 36.54, 32.00, 28.32, 28.24, 28.14, 26.96, 22.06. LC-MS (method 1): t_R_=3.49 min, m/z = 793.7 [M+H]^+^; calcd: 793.5.

**Synthesis of L2:** To compound **11** (155.5 mg, 0.2 mmol) in dichloromethane (5 mL), cooled to 0 ℃, was added anisole (0.4 mL, 2 mL/mmol) followed by the slow addition of trifluoroacetic acid (5 mL). The mixture was stirred for 1 h at 0 ℃ and then at room temperature for 4 h. The solvent and trifluoroacetic acid were carefully evaporated under reduced pressure, the resulting oily residue was redissolved in water (5 mL), and the organic byproducts were decanted with diethyl ether (3 x 5 mL). The aqueous layer was lyophilized, and the obtained crude product was purified by preparative HPLC to give **L2** as a white solid (method 8, 84.9 mg, 81%). ^1^H NMR (500 MHz, D2O) δ 7.61 (t, J = 7.7 Hz, 1H), 7.07 (d, J = 7.8 Hz, 2H), 3.88 (s, 4H), 3.82 (dd, J = 8.1, 4.3 Hz, 1H), 3.27 (s, 4H), 3.21 – 2.84 (m, 13H), 2.25 – 2.10 (m, 2H), 1.90 – 1.77 (m, 3H), 1.74-1.70 (m, 1H). ^13^C NMR (126 MHz, D2O) δ 179.23, 179.07, 176.12, 159.03, 138.73, 121.20, 84.38, 60.51, 58.37, 51.45, 49.95, 45.04, 36.56, 32.05, 27.58, 20.98. LC-MS (method 2): t_R_=5.58 min, m/z = 525.3 [M+H]^+^; calcd: 525.3.

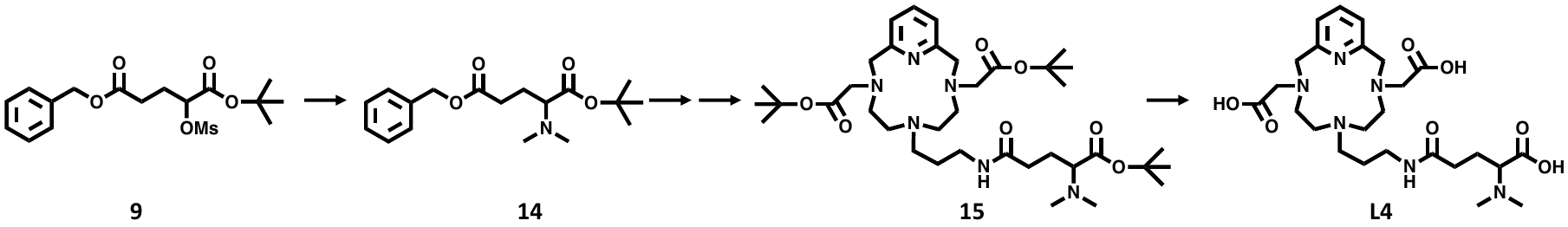

**Synthesis of 14.** Compound **9** (3.72 g, 10.0 mmol) and diethylamine (2 M in THF, 7.5 mL, 15.0 mmol) were mixed together in 40 mL dry acetonitrile, which were added K_2_CO_3_ (2.76 g, 20.0 mmol) and NaI (0.15 g, 1.0 mmol). The mixture was stirred under reflux for 12 h. The solid was then removed by filtration and washed with dichloromethane. The crude product was purified via flash chromatography to give **14** as a yellow oil (method 7, 1.70 g, 53%). ^1^H NMR (500 MHz, Chloroform-*d*) δ 7.39 – 7.25 (m, 5H), 5.09 (s, 2H), 3.03 (t, *J* = 7.6 Hz, 1H), 2.40 (t, *J* = 7.5 Hz, 2H), 2.31 (s, 6H), 1.95 (q, *J* = 7.5 Hz, 2H), 1.44 (s, 9H). ^13^C NMR (126 MHz, Chloroform-*d*) δ 173.14, 171.09, 136.12, 128.62, 128.24, 81.26, 67.05, 66.26, 41.48, 30.90, 28.37, 24.74. LC-MS (method 1): t_R_=4.15 min, m/z = 322.4 [M+H]^+^; calcd: 322.2.

**Synthesis of 15.** Compound **14** (0.32 g, 1 mmol) was added to a slurry of palladium on carbon (50% water, 50 mg) in methanol (20 mL). The mixture was subject to two cycles of vacuum and hydrogen purge and then stirred under an atmosphere of hydrogen for 12 h under room temperature. After evacuating the system of hydrogen, celite was added and the slurry filtered through a methanol-wet bed of celite. The filtrate was concentrated under reduced pressure to give a colorless oil of product which was used in the next step without further purification. The obtained colorless oil, compound 4 (0.25 g, 0.5 mmol), and *N,N*-diisopropylethylamine (0.13 g, 1.0 mmol) were dissolved in dry acetonitrile (20 mL), HATU (0.29 g, 0.75 mmol) in dry acetonitrile (10 mL) was added. The reaction mixture was stirred at room temperature for 45 min. After evaporated under reduced pressure, the obtained oil was redissolved in dichloromethane and extract with 10% citric acid aqueous solution and brine. The organic layer was dried (Na_2_SO_4_) and concentrated under reduced pressure. The crude product was purified via flash chromatography to give **15** as a yellow oil (method 7, 0.27 g, 81%). ^1^H NMR (500 MHz, Chloroform-*d*) δ 7.58 (t, *J* = 7.7 Hz, 1H), 7.01 (d, *J* = 7.7 Hz, 2H), 5.25 (s, 1H), 3.91 (s, 4H), 3.51 – 2.98 (m, 16H), 2.34 – 2.15 (m, 8H), 1.92 (ddt, *J* = 22.6, 8.4, 6.4 Hz, 4H), 1.40 (d, *J* = 1.9 Hz, 27H). ^13^C NMR (126 MHz, Chloroform-*d*) δ 173.88, 170.25, 167.01, 138.23, 120.88, 82.12, 81.12, 67.22, 58.36, 57.95, 52.05, 50.38, 44.89, 41.32, 36.43, 32.62, 28.35, 28.21, 25.46, 21.73. LC-MS (method 1): t_R_=2.84 min, m/z = 705.5 [M+H]^+^; calcd: 705.5.

**Synthesis of L4.** To compound **15** (132.7 mg, 0.2 mmol) in dichloromethane (5 mL) cooled to 0 ℃, was added anisole (0.4 mL, 2 mL/mmol) followed by the slow addition of trifluoroacetic acid (5 mL) and the mixture was stirred at 0 ℃ for 1 h followed by room temperature for 4 h. The solvent and trifluoroacetic acid were gently evaporated under reduced pressure, and the resulting oily residue was dissolved in water (5 mL). Then organic byproducts were pipetted away with diethyl ether (3 x 5 mL), the aqueous layer was freeze-dried, and the obtained crude product was purified by preparative HPLC to give **L4** as a white solid (method 8, 89.0 mg, 83%). The concentration of the metal solution was determined by ICP-MS. ^1^H NMR (500 MHz, D_2_O) δ 8.24 (t, *J* = 7.9 Hz, 1H), 7.63 (d, *J* = 7.9 Hz, 2H), 4.37 (s, 4H), 3.84 (dd, *J* = 9.6, 3.8 Hz, 1H), 3.75 (s, 4H), 3.28 – 2.88 (m, 12H), 2.78 (d, *J* = 15.1 Hz, 6H), 2.43 – 2.24 (m, 2H), 2.15 (ddt, *J* = 15.9, 8.0, 3.9 Hz, 1H), 2.01 (qt, *J* = 8.0, 5.8 Hz, 1H), 1.90 – 1.77 (m, 2H). ^13^C NMR (126 MHz, D_2_O) δ 175.10, 173.93, 169.94, 151.97, 146.63, 124.08, 66.60, 57.70, 56.60, 52.86, 51.82, 50.83, 42.61, 40.07, 36.40, 31.06, 22.62, 22.00. LC-MS (method 2): t_R_=6.64 min, m/z = 537.2 [M+H]^+^; calcd: 537.3.

**MnL1**: LC-MS (method 3): t_R_=7.51 min, m/z [M + H]^+^ calcd for [C_23_H_36_N_7_O_7_Mn]^+^, 577.2; found 577.2, purity = 96% (250 nm UV detector).

**MnL2**: LC-MS (method 3): t_R_=8.02 min, m/z [M + H]^+^ calcd for [C_23_H_35_N_6_O_8_Mn]^+^, 578.2; found 578.1, purity = 98% (250 nm UV detector).

**MnL4**: LC-MS (method 3): t_R_=8.06 min, m/z [M + H]^+^ calcd for [C_25_H_39_N_6_O_7_Mn]^+^, 590.2; found 590.1, purity = 98% (250 nm UV detector).

**Radiosynthesis.** ^52^MnCl_2_ was purchased from University of Alabama at Birmingham Radiology Cyclotron Facility. All the buffers and solutions were prepared with ultrapure water, purified over Chelex 100 resin (Bio rad) and filtered using a 0.2 μm sterile filter. To solutions of ligands L1, L2, L3, or L4 in 0.15 M NaOAc buffer (1 mg/mL, 180 μL, pH 4.5), was added 20 μL ^52^MnCl_2_ in 0.01M HCl (5 MBq), and the solution was heated at 65 °C for 5 min. After cooling to room temperature, 200 μL of PBS was added, and the pH adjusted to 6 by 1 M NaOH. The radiochemical purity of the dose to be injected was determined to be >95% for all four labeled compounds using method 9 (Supplementary Fig. 9). The reactivities of the ^52^Mn-labeled complexes with aldehyde were confirmed by radio-HPLC measurement of aldehyde spiked samples (2 μL, 10 mM pyridine-2-carbaldehyde spiked to 20 μL radiolabeling solution).

### Supplementary Figures

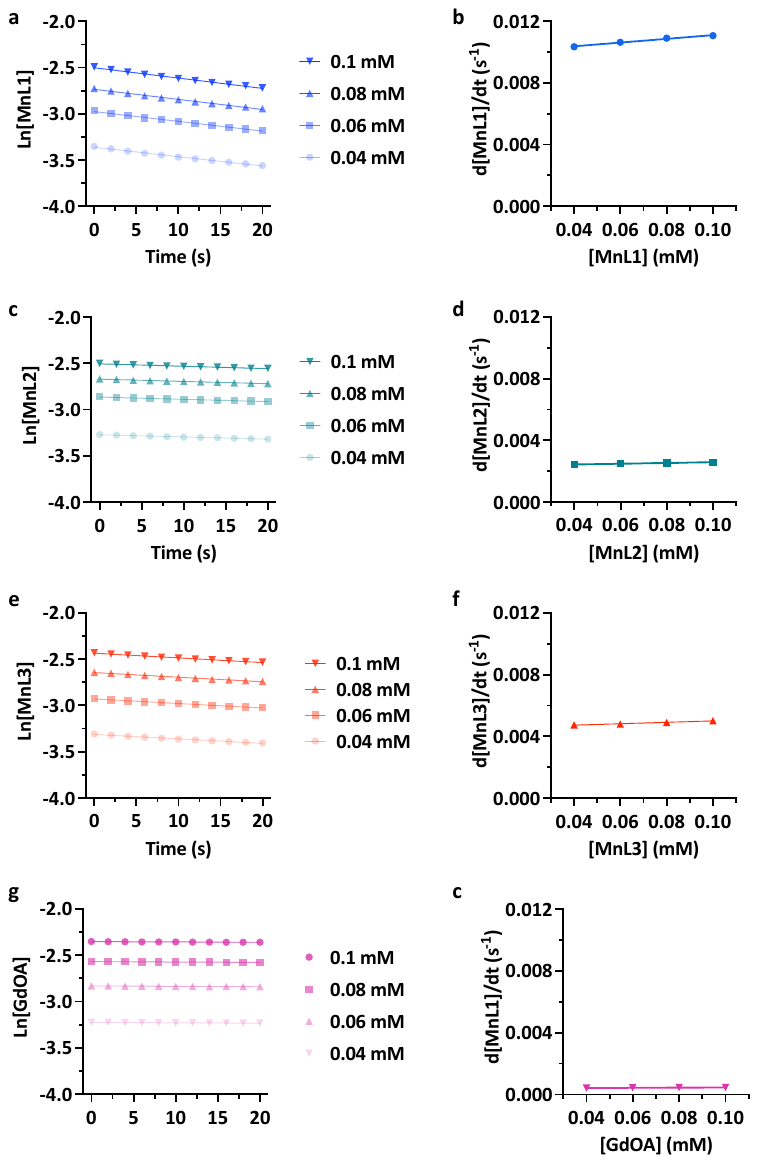

**Supplementary Figure 1 | Condensation reaction rate constant measurements.** The kinetics of condensation reaction of MnL1, MnL2, MnL3, and GdOA with butyraldehyde were measured spectrophotometrically under pseudo-first order conditions with respect to butyraldehyde (pH 7.4, PBS, 25°C). **a**, **c**, **e**, **g**, Time-dependent hydrazone formation of different concentration of MnL1, MnL2, MnL3, and GdOA in the presence of 1 mM butyraldehyde. Observed pseudo first-order reaction rates of different concentrations of probe with butyraldehyde were calculated from the slopes. **b**, **d**, **f**, **h**, Plots of observed condensation reaction rate of 0.04, 0.06, 0.08, 0.1 mM of MnL1, MnL2, MnL3, and GdOA with 1 mM butyraldehyde as a function of Mn complex concentration. Second-order reaction rate constants were calculated from the slopes.

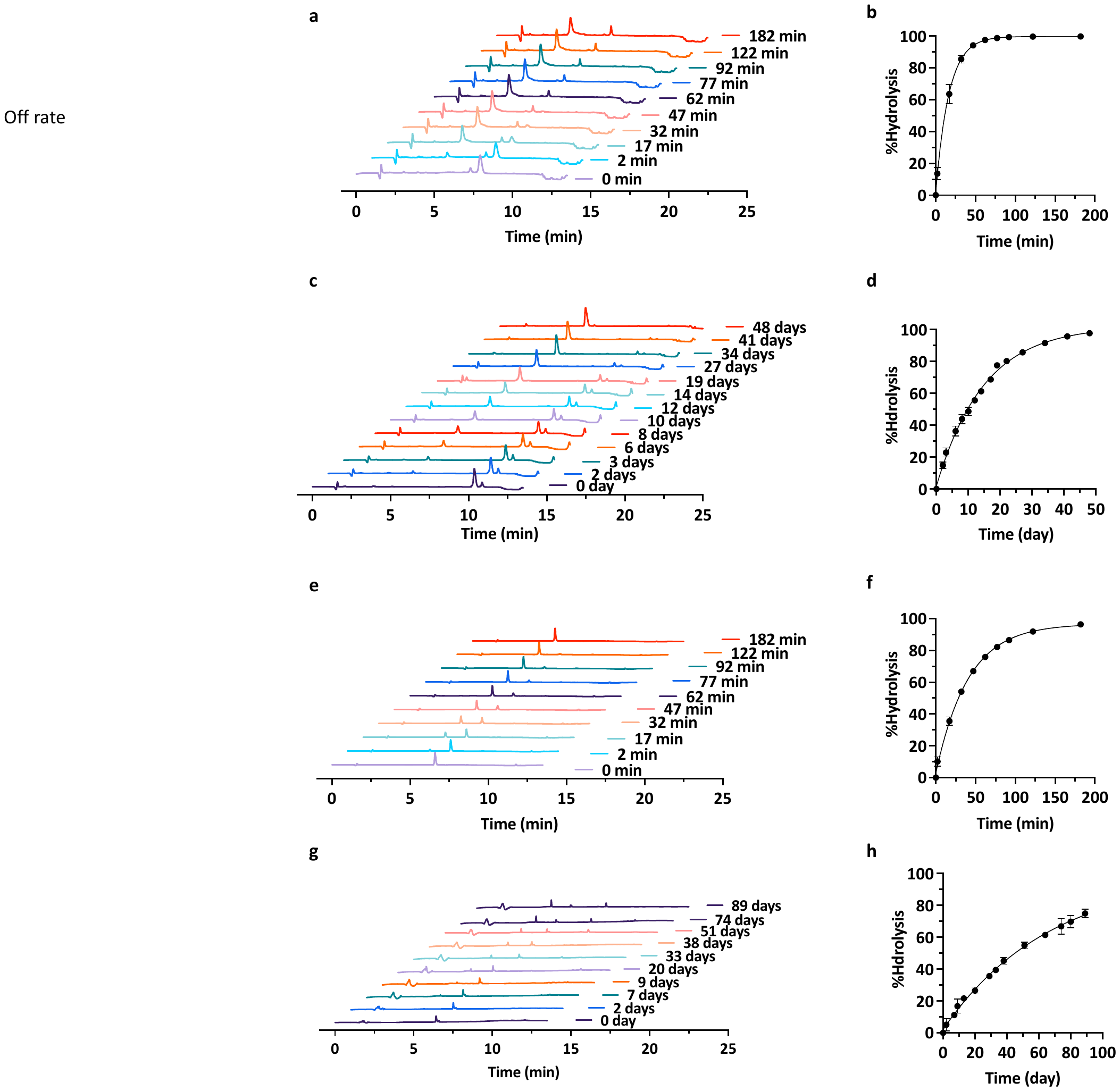

**Supplementary Figure 2 |** **Hydrolysis rate constant measurement.** The kinetics of the hydrolysis reactions of MnL1-hydrazone, MnL2-oxime, MnL3-hydrazone, and GdOA-oxime (formed by condensation between MnL1, MnL2, MnL3, and GdOA with butyraldehyde) were measured by HPLC (1 mM, pH 7.4, PBS, 25°C, HPLC method 6). Waterfall plots of HPLC traces of MnL1-hydrazone (**a**), MnL2-oxime (**c**), MnL3-Hydrazone (**e**), and GdOA-oxime (**g**) pre- and post-addition of formaldehyde (100 mM) as a hydrazine or oxyamine scavenger. %Hydrolysis vs time for the hydrolysis of MnL1-hydrazone (**b**), MnL2-oxime (**d**), MnL3-Hydrazone (**f**), and GdOA-oxime (**h**) obtained by integration of HPLC trace peaks corresponding to the hydrazone or oxime (n = 3 for each group).

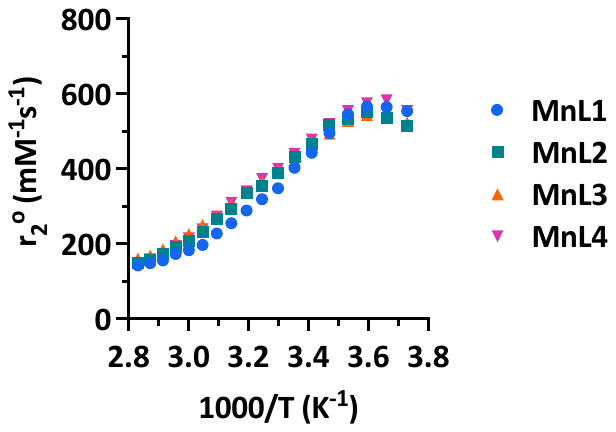

**Supplementary Figure 3 |** **Measurement of hydration number each Mn complex.** The number of coordinated water ligands can be calculated from the maximum ^17^O relaxivity (r_2_^o^) recorded at 11.7 T using the method reported by Gale et al^7^. Complexes (0.88 ± 0.15 mM) were prepared in water at pH 7.0 and the H_2_^17^O linewidth was measured as a function of temperature. r_2_^O^ was calculated as the difference in O-17 linewidth at half max (ν_1/2_) at a given temperature between a solution containing Mn and a solution without Mn (ref sample) and then normalized to Mn concentration, r_2_^O^ = (πν_1/2_(Mn)- πν_1/2_(ref))/[Mn].

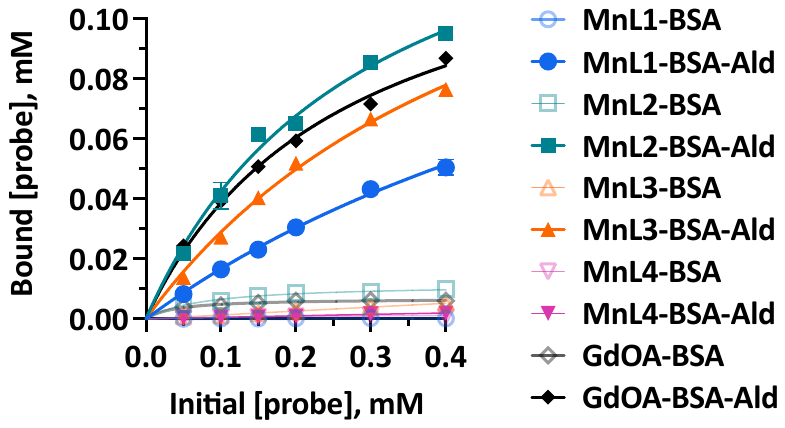

**Supplementary Figure 4 |** **Measurement of protein bound complex concentration.** The concertation of protein-bound MnL1, MnL2, MnL3, MnL4, and GdOA incubated with BSA-Ald or BSA as determined by ICP-MS (Data are presented as mean± SEM, n = 3 for each group). The samples passed through an ultrafiltration filter (10 KDa cut-off) to separate the protein-bound probe from the unbound probe. The Mn and Gd concentration of the residue was determined using ICP-MS. There is a concentration-dependent increase in binding of both MnL1, MnL2, MnL3, and GdOA to BSA-Ald, while nonspecific binding to unmodified BSA was low for all probes. MnL4 exhibited little/no binding to both BSA or BSA-Ald.

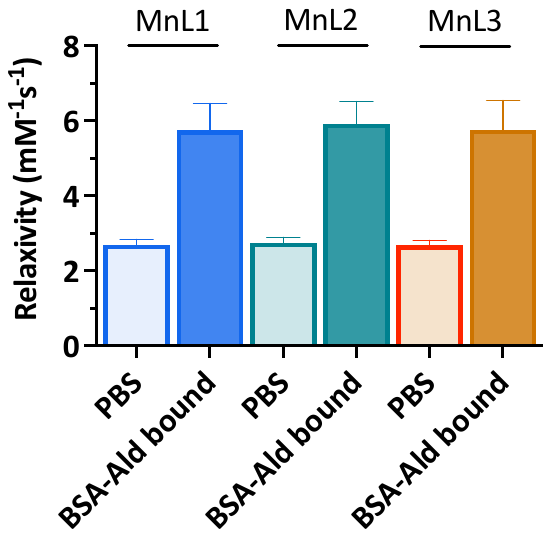

**Supplementary Figure 5 |** **Relaxivities of MnL1, MnL2, and MnL3 at 4.7 T.** At 4.7 T, relaxivities of MnL1, MnL2, and MnL3 were about 3 mM^-1^ s^-1^ in PBS and exhibited a 70% enhancement after binding to BSA-Ald (4.7 T, 37°C).

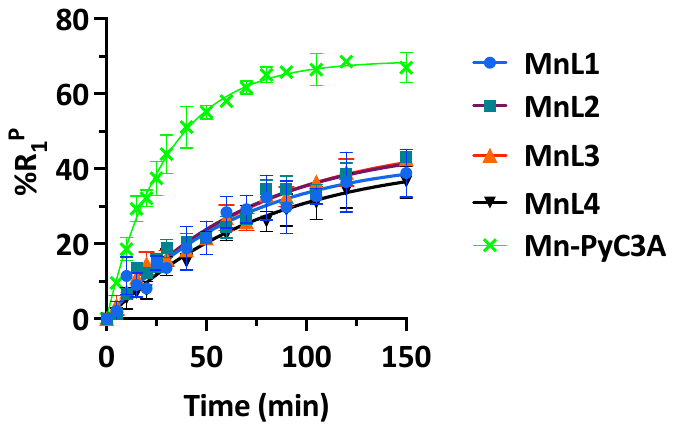

**Supplementary Figure 6 |** **Kinetic inertness of Mn-based probes.** Transmetalation of 1 mM MnL1, MnL2, MnL3, MnL4, and Mn-PyC3A by 25 mM Zn^2+^ monitored by %R_1_^P^ change as a function of time in 50 mM pH 6.0 MES buffer, 37 °C, 1.4 T (Data presented as mean ± SEM, n = 3 for each group). The macrocyclic PC2A-Mn-based complexes exhibited 2-fold higher kinetic inertness compared to Mn-PyC3A.

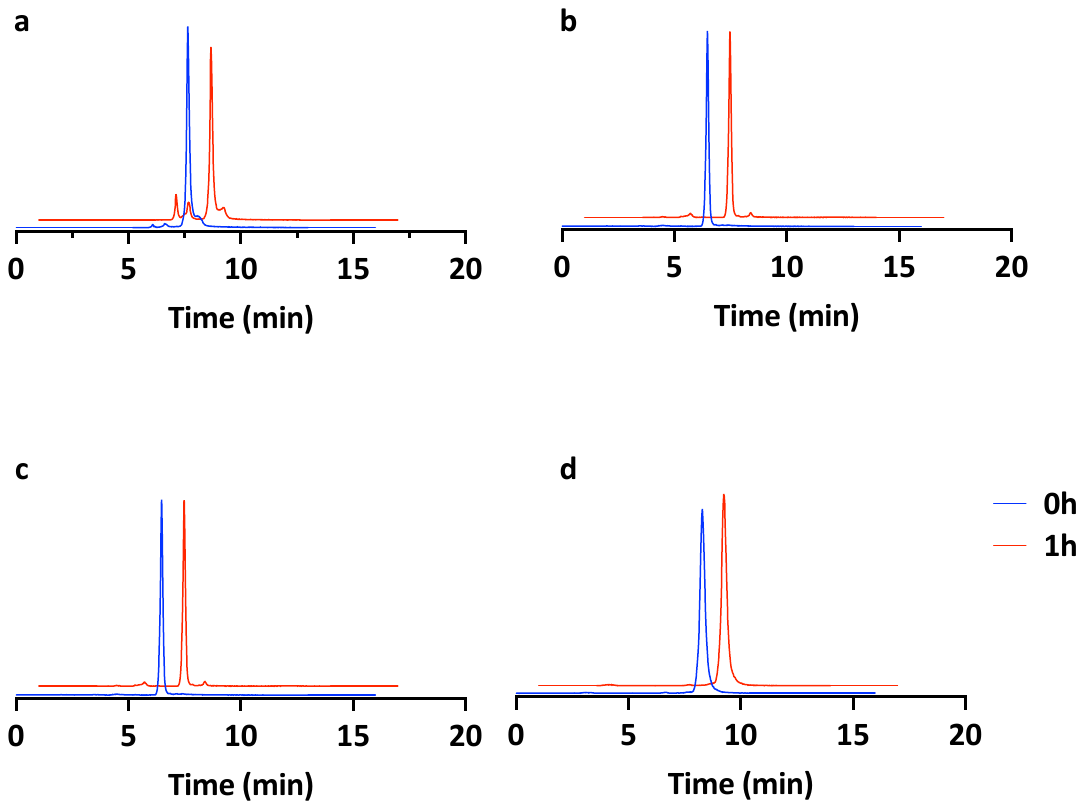

**Supplementary Figure 7 |** **Stability of Mn-based probes in human plasma.** LC-ICP-MS traces analyzed for Mn from solutions of MnL1 (**a**), MnL2 (**b**), MnL3 (**c**), and MnL4 (**d**) incubated in human plasma at 37 °C for 0, and 1 h.

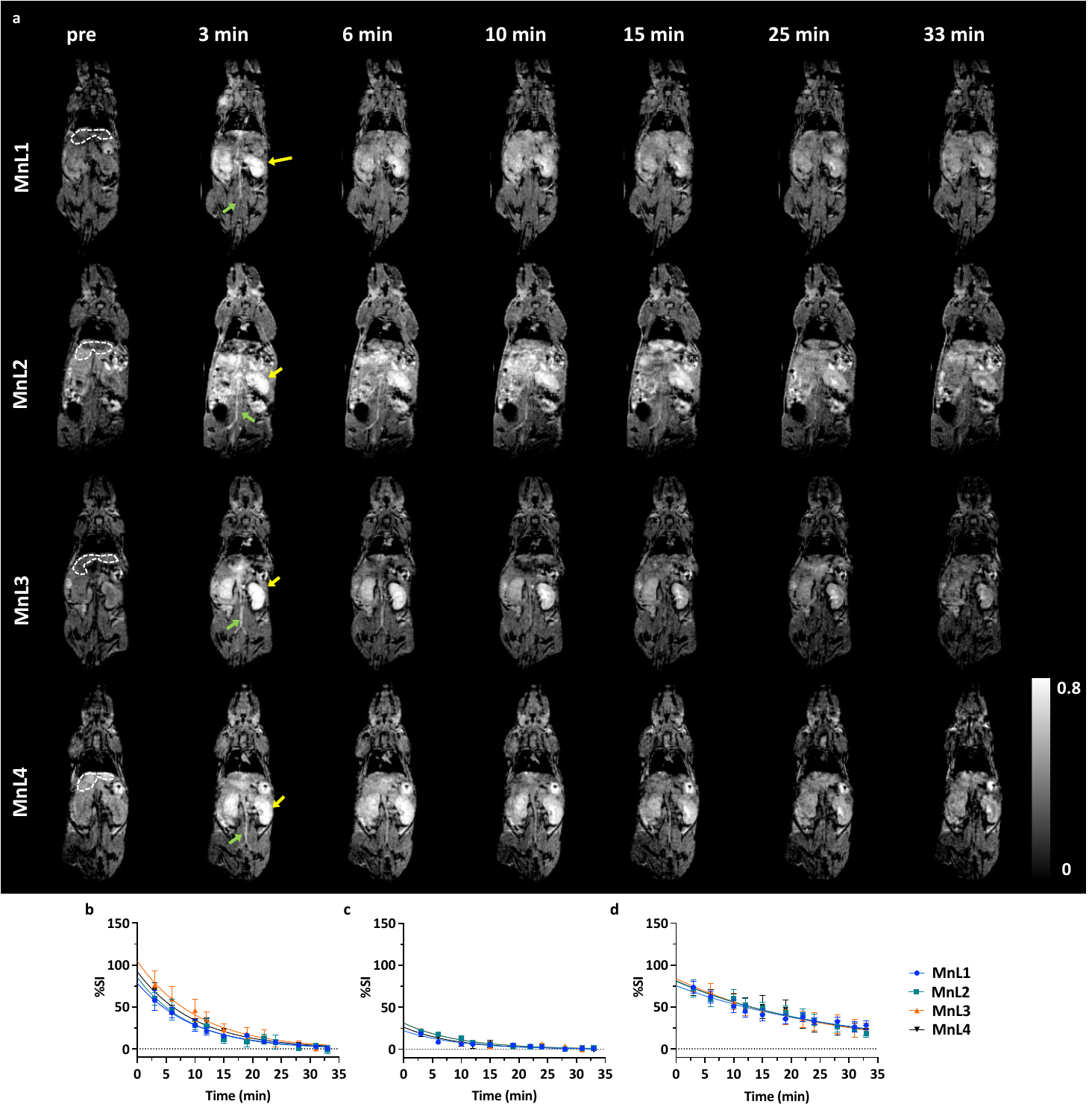

**Supplementary Figure 8 |** **Pharmacokinetics of Mn-based probes.** **a**, Representative time-dependent coronal T1-weighted 3D-FLASH MR images of naïve mice before and at several time points after injection with MnL1, MnL2, MnL3, or MnL4. In the field of view at the slice plane, the kidney (yellow), part of liver (white), aorta (green arrow) are visible. Immediately following injection, the signal is enhanced in all tissue and clears rapidly with time. The clearance of the probe from the blood was measured by determining the signal in the blood (aorta) and fitting the data to a monoexponential decay. Quantification of the time dependent change in signal intensity change in the aorta (**b**), liver (**c**), kidney (**d**), and normalized aorta-to-liver ratio (**e**), normalized to pre-injection aorta-to-liver ratio are shown graphically (Data presented as mean ± SEM, n = 4 for each group).

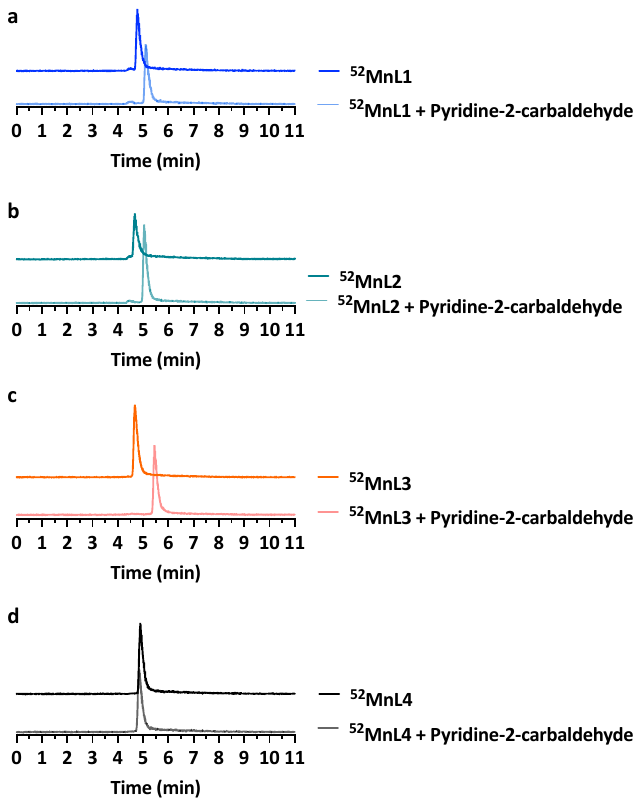

**Supplementary Figure 9 |** **Radio-HPLC trace.** After radiolabeling, the purity of ^52^MnL1 (**a**), ^52^MnL2 (**b**), ^52^MnL3 (**c**), and ^52^MnL4 (**d**) were measured by radio-HPLC (HPLC method 9, radiochemical purity >96%). The reactivity of radiolabeled compound to aldehyde was confirmed by adding pyridine-2-carbaldehyde to the HPLC vial and then reanalyzing by radio-HPLC. A new species is formed when pyridine-2-carbaldehyde is spiked into ^52^MnL1, ^52^MnL2, and ^52^MnL3 solution, while the retention time remained the same for ^52^MnL4 spiked with pyridine-2-carbaldehyde.

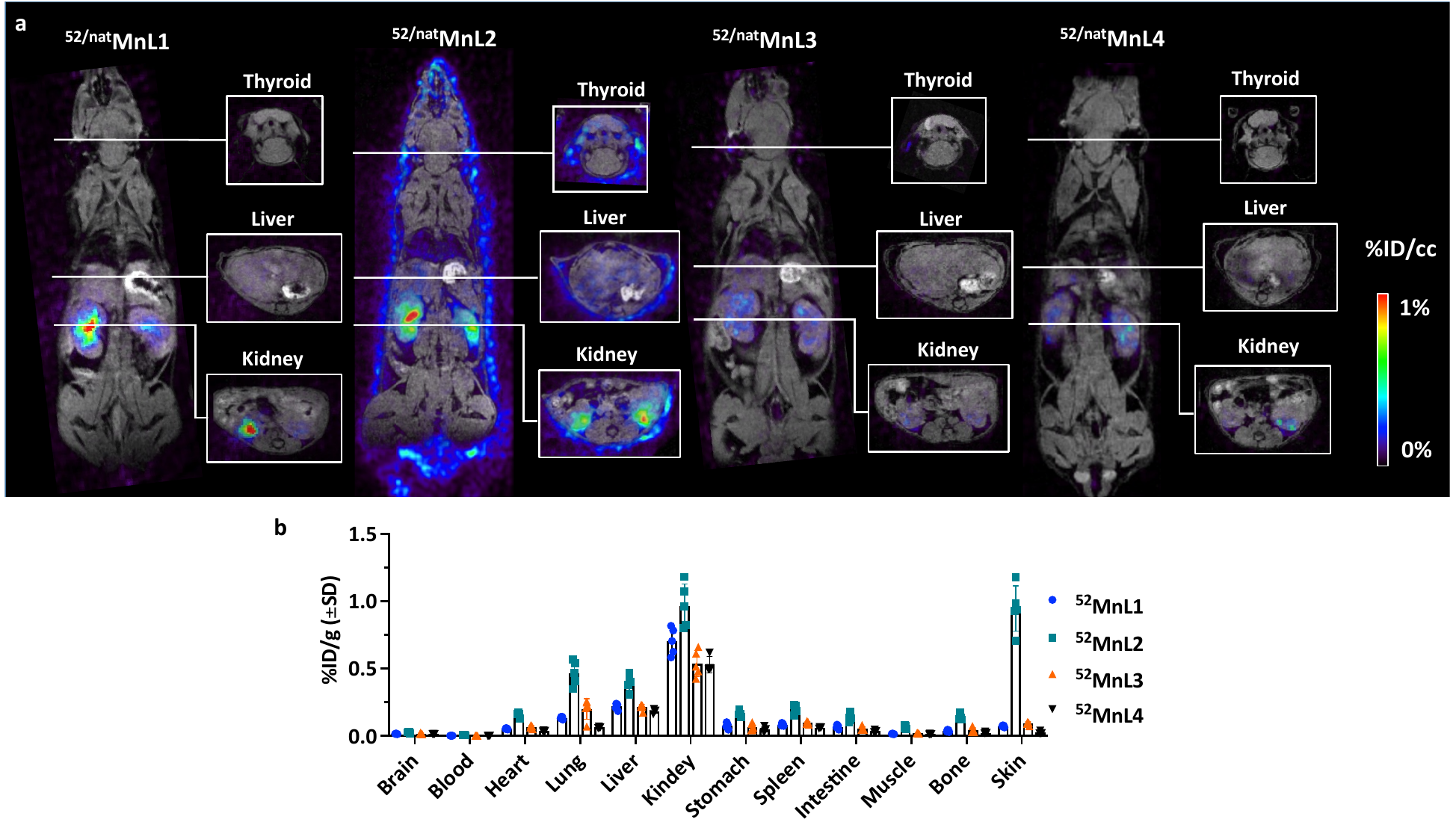

**Supplementary Figure 10** **|** **Biodistribution of Mn(II)-based probes.** **a**, Representative fused PET (color scale) - MRI (grayscale) coronal and axial images of naïve mice 24 h post-injection of ^52/nat^MnL1, ^52/nat^MnL2, ^52/nat^MnL3, and ^52/nat^MnL4. Negligible radioactivity was observed in the salivary glands or liver (sites of free Mn^2+^ deposition) and demonstrated that each probe likely remained intact 24 h post-injection. Owing to the much higher stability of oxime bond, higher retention of ^52/nat^MnL2 was seen compared to hydrazine-based or non-targeted probes, especially in skin. **b**, Ex vivo bio-distribution of ^52^MnL1, ^52^MnL2, ^52^MnL3, and ^52^MnL4. Data expressed as percent injected dose per gram of tissue (%ID/g) (mean ± SD) in various organs.

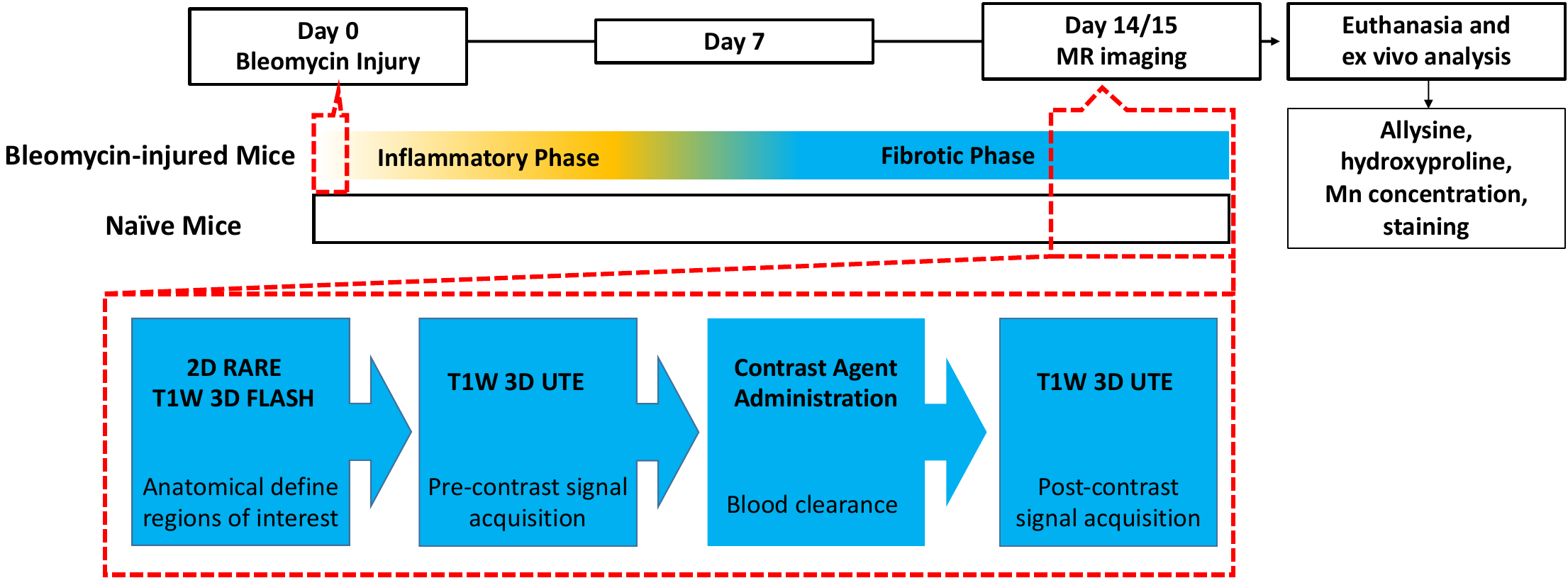

**Supplementary Figure 11 |** **Study timeline and imaging sequences.** Bleomycin administration to initiate early (oxidative and inflammatory) and late (fibrotic) lung injury. On day 14 and day 15, the animals were imaged dynamically with MnL1, MnL2, MnL3, or MnL4, and then euthanized, and the organs were harvested for analysis. For MRI, anatomical 2D RARE and T1-weighted 3D FLASH images were acquired before probe injection and are used along with the 3D-FLASH images post-injection to define regions of interest (ROIs). The T1-weighted 3D UTE images obtained before and after Mn probe injection were used to quantify the signal enhancement in the lungs before and after probe injection.

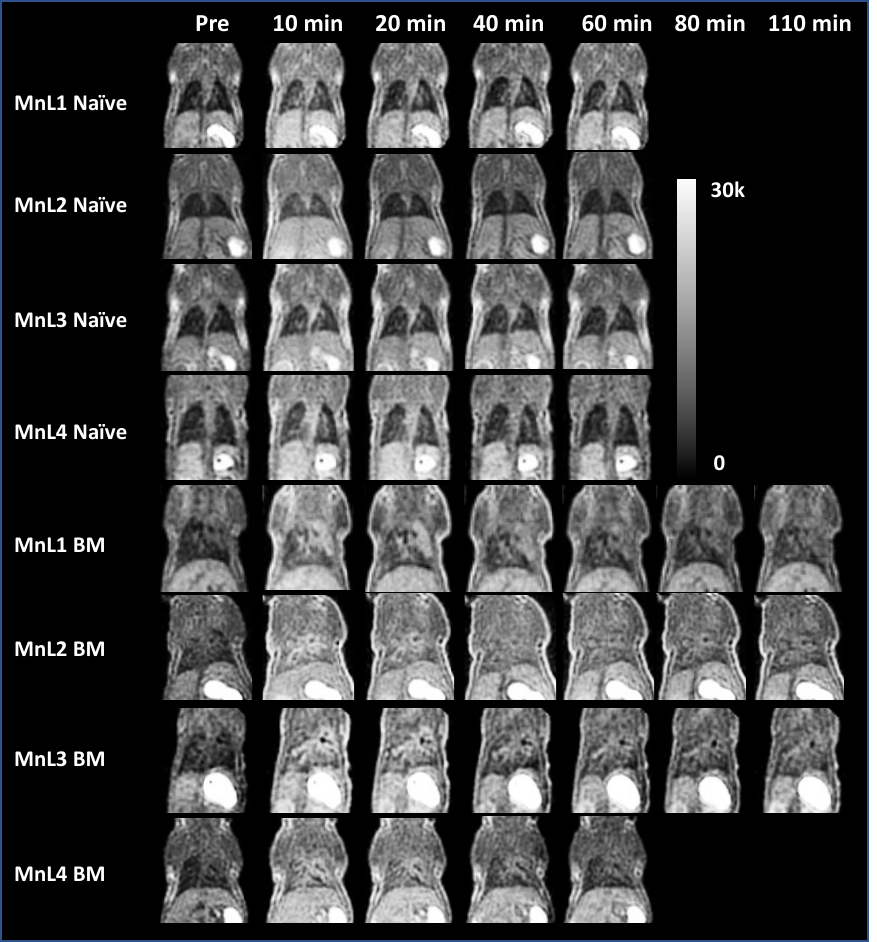

**Supplementary Figure 12 |** **3D UTE MR images of naïve and bleomycin-injured (BM) mice before and dynamically post intravenous administration of 0.1 mmol•kg^-1^ MnL1, MnL2, MnL3, and MnL4.** All the probes exhibited minimal signal enhancement in normal lungs. The nonspecific probe, MnL4, showed higher signal enhancement in fibrotic lung likely as a result of higher extracellular volume caused by the injury. In comparison to naïve mice, at 60 min post-injection of the allysine-targeted probes, higher signal enhancement was observed in fibrotic lungs, and the oxyamine-based MnL2 probe exhibited superior long-lasting MRI signal enhancement. Due to the higher hydrolytic stability of the hydrazone formed by MnL3 compared to that formed with MnL1, the signal enhancement in the fibrotic lung was greater with MnL3.

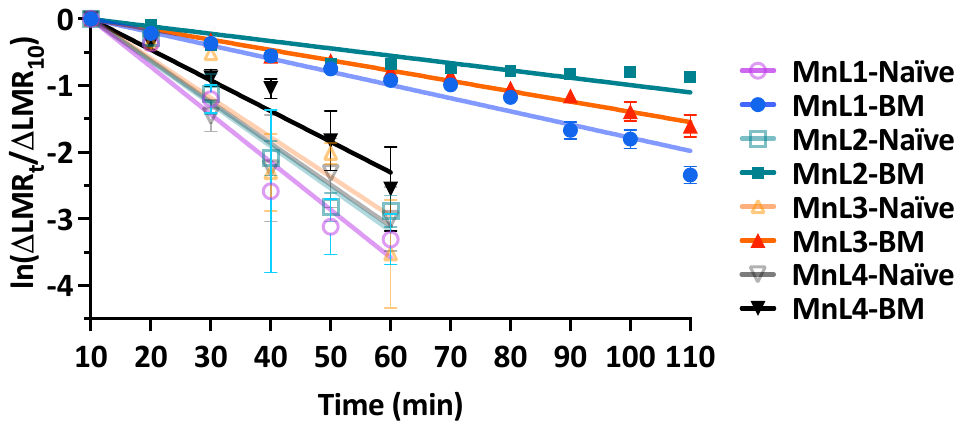

**Supplementary Figure 13 |** **Probe washout rate in naïve and bleomycin-injured (BM) lungs measured by MRI.** The natural logarithm of the ratio of lung signal to muscle ratio, ln (∆LMR_t_ / ∆LMR_10_) is plotted as a function of time, where ∆LMR_10_ is the change of lung to muscle ratio (∆LMR) at 10 min post-injection and ∆LMR_t_ is ∆LMR at time *t.* The half-life for the probe in the lung is estimated from the slope of these curves. Data presented as mean ± SEM. Naïve mice: n = 6 for MnL1, n = 6 for MnL3, n = 4 for MnL4; BM mice: n = 13 for MnL1, n = 12 for MnL3, n = 5 for MnL4.

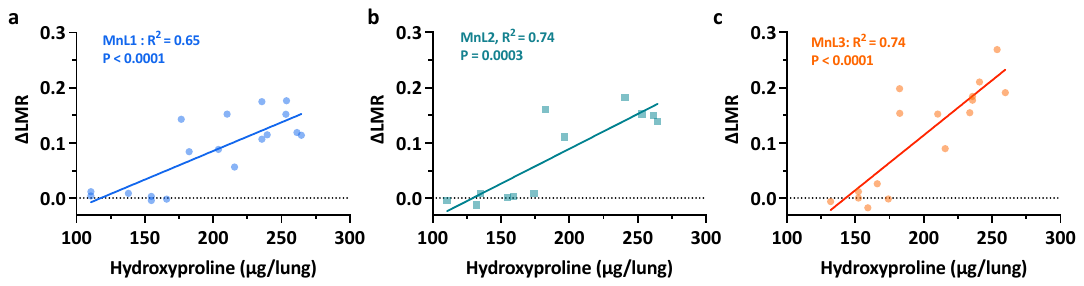

**Supplementary Figure 14 |** **Correlation between hydroxyproline content and MRI signal enhancement.** The MRI signal enhancement in naïve and BM mice imaged with MnL1 (**a**), MnL2 (**b**), and MnL3 (**c**) correlates well with hydroxyproline content.

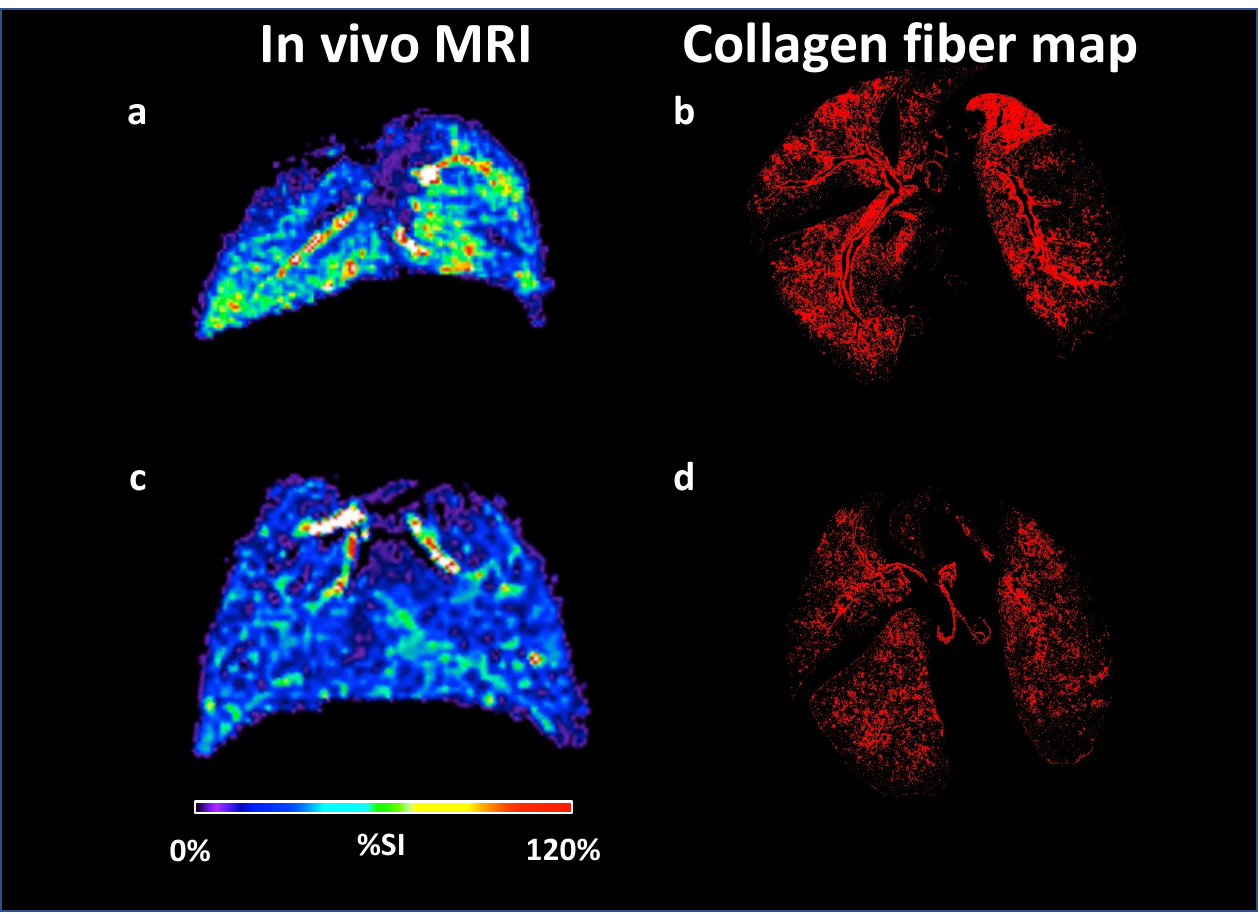

**Supplementary Figure 15 | Histological validation of lung fibrotic region MR imaging.** The percentage MRI signal change produced by MnL3 (**a, c**) exhibited similar distribution patterns compared with histological staining (**b, d**, red signal presenting extracted collagen signal in Masson’s Trichrome staining)

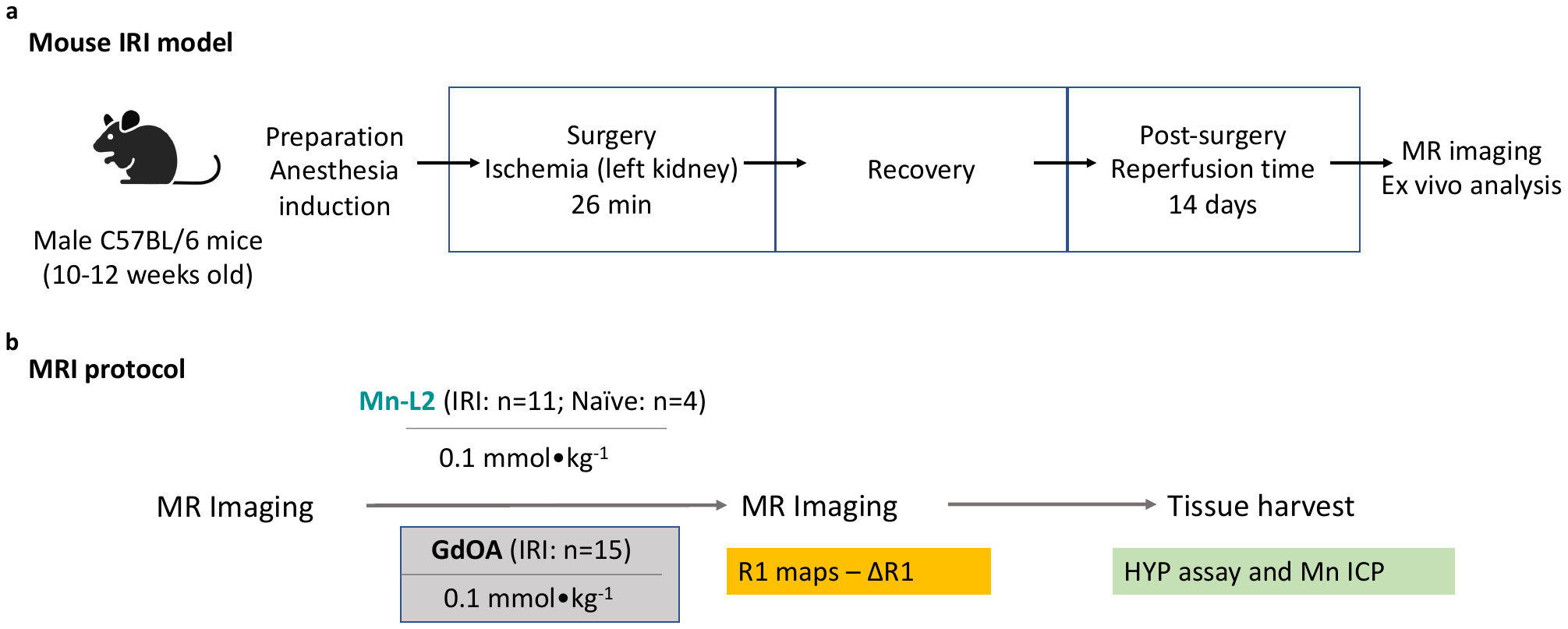

**Supplementary Figure 16 |** **Experimental renal IRI model in mice and in vivo study protocol.** **a**, Phases of experiments and interventions (anesthesia induction, ischemia, and reperfusion) are shown. **b**, 14 days post-surgery, naïve and IRI mice were imaged on a 9.4 T scanner prior to and post-injection of MnL2, and GdOA. After imaging, the IRI kidney, contralateral kidneys, and normal kidneys were harvested, separated using microdissection and kept under -80 °C for ex vivo analysis.

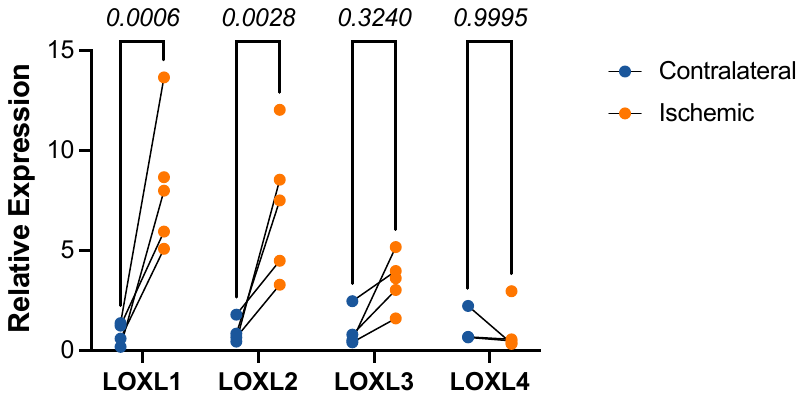

**Supplementary Figure 17 |.** Expression levels of LOXL 1-4 genes relative to 18S rRNA in the ischemic and contralateral kidneys. Mixed effect analysis with Sidak’s multiple comparison test (n=3-5).

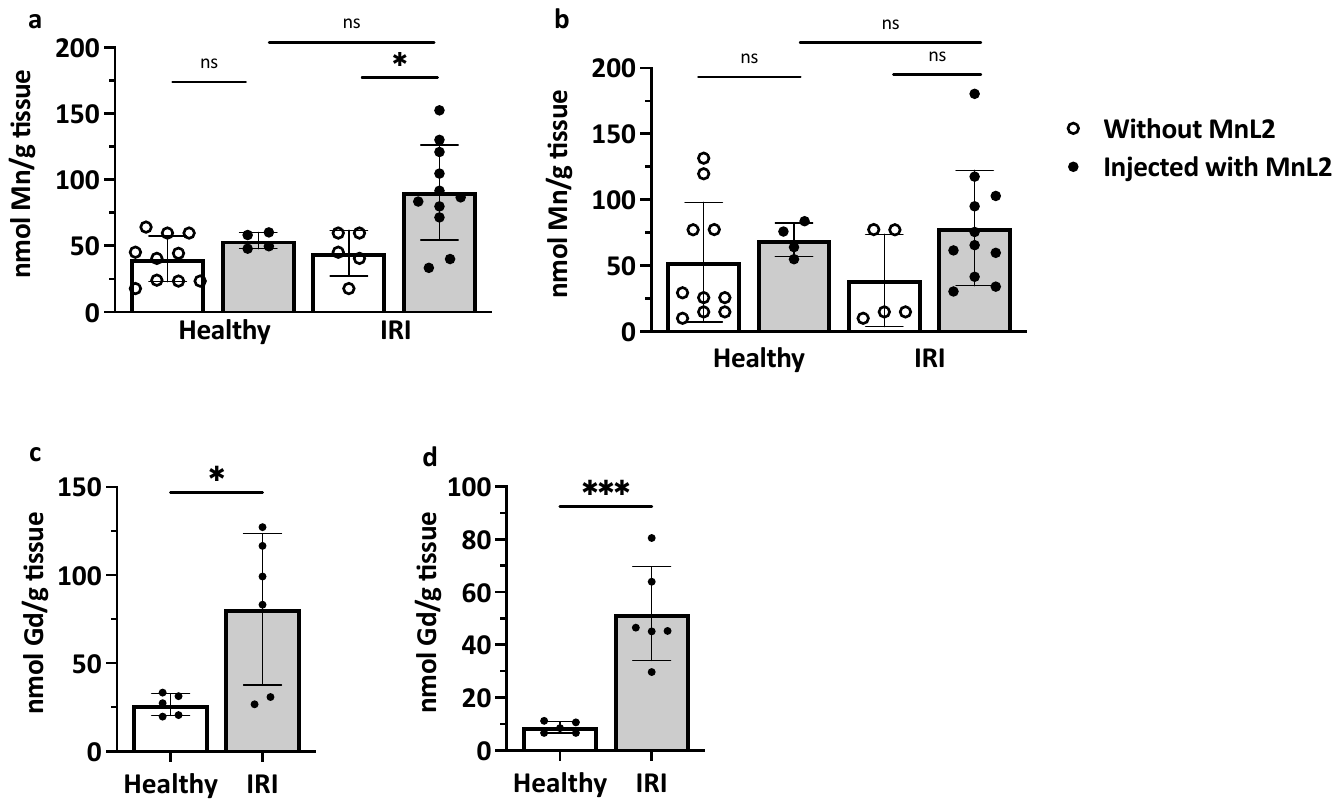

**Supplementary Figure 18 | Ex vivo kidney Mn and Gd quantification at 4 h post-injection of MnL2 or GdOA.** Because Mn is an endogenous element we measured Mn levels in mice that did not receive MnL2 to assess basal Mn levels in renal cortex and medulla. **a**, In kidney cortex from healthy mice there was no significant increase in Mn levels injected with MnL2 compared to mice naïve to MnL2 demonstrating efficient elimination of injected MnL2. In IRI mice, the Mn concentration in kidney cortex was significantly higher in MnL2 injected mice than in animals naïve to MnL2. **b**, In the medulla there was no significant difference between healthy mice treated with MnL2 and mice naïve to MnL2, not was there a significant differemce in IRI mice. (Naïve mice: n = 10 for non-injection group, n = 4 injected with MnL2; IRI mice: n = 5 for non-injection group, n = 11 injected with MnL2). Statistical analysis was performed using two-tailed t test, ns, not significant. c, Gd concentration was significantly higher in cortex (**c**) and medulla (**d**) of IRI kidneys 4 h post-injection of GdOA (n = 5 for naïve mice, n = 6 for IRI mice). Statistical analysis was performed using two-tailed t test, * p < 0.05, *** p < 0.001.

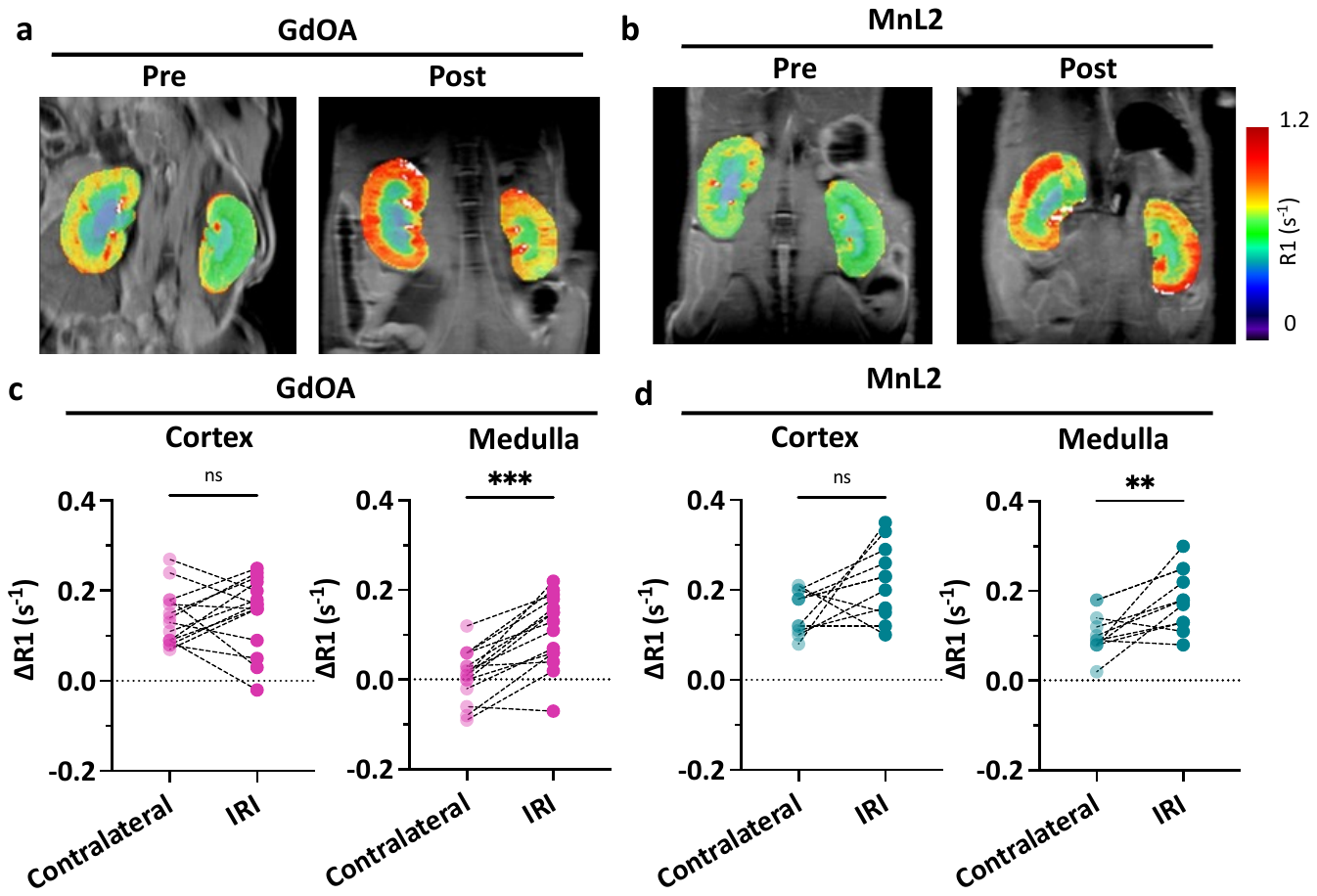

**Supplementary Figure 19 |** **Molecular MRI of renal fibrogenesis.** Representative MR images obtained from mice with renal IRI, 14 days after the injury. Images were obtained prior to and 4 h post-injection of GdOA (**a**), and MnL2 (**b**). Quantification of ∆R1 in the cortex and medulla of IRI mice injected with GdOA (**c**) or MnL2 (**d**). MnL2 with a higher condensation reaction rate constant compared to GdOA produced higher dynamic range for renal fibrogenesis imaging. Data are presented as mean ± s.d.; IRI mice: n = 15 for GdOA, n = 11 for MnL2. Statistical analysis was performed using two-tailed paired Student’s *t* test, ** p < 0.01, *** p < 0.001, ns, not significant.

### Supplementary Table

**Table 1 | Molar extinction coefficient of molecular MR probes and the hydrazone product formed with butyraldehyde**

| Probe | *ε* x10^4^ (M^-1^cm^-1^) |
| --- | --- |
| MnL1 | 0.48 ± 0.02 |
| MnL1-BAld | 1.07 ± 0.08 |
| MnL2 | 0.52 ± 0.02 |
| MnL2-BAld | 0.97 ± 0.02 |
| MnL3 | 0.18 ± 0.02 |
| MnL3-BAld | 1.92 ± 0.01 |
| GdOA | 0.068 ± 0.004 |
| GdOA-BAld | 0.36 ±0.06 |

n = 3 independent samples

### NMR spectra

**Compound 2: ^1^H NMR (500 MHz, CDCl_3_)**

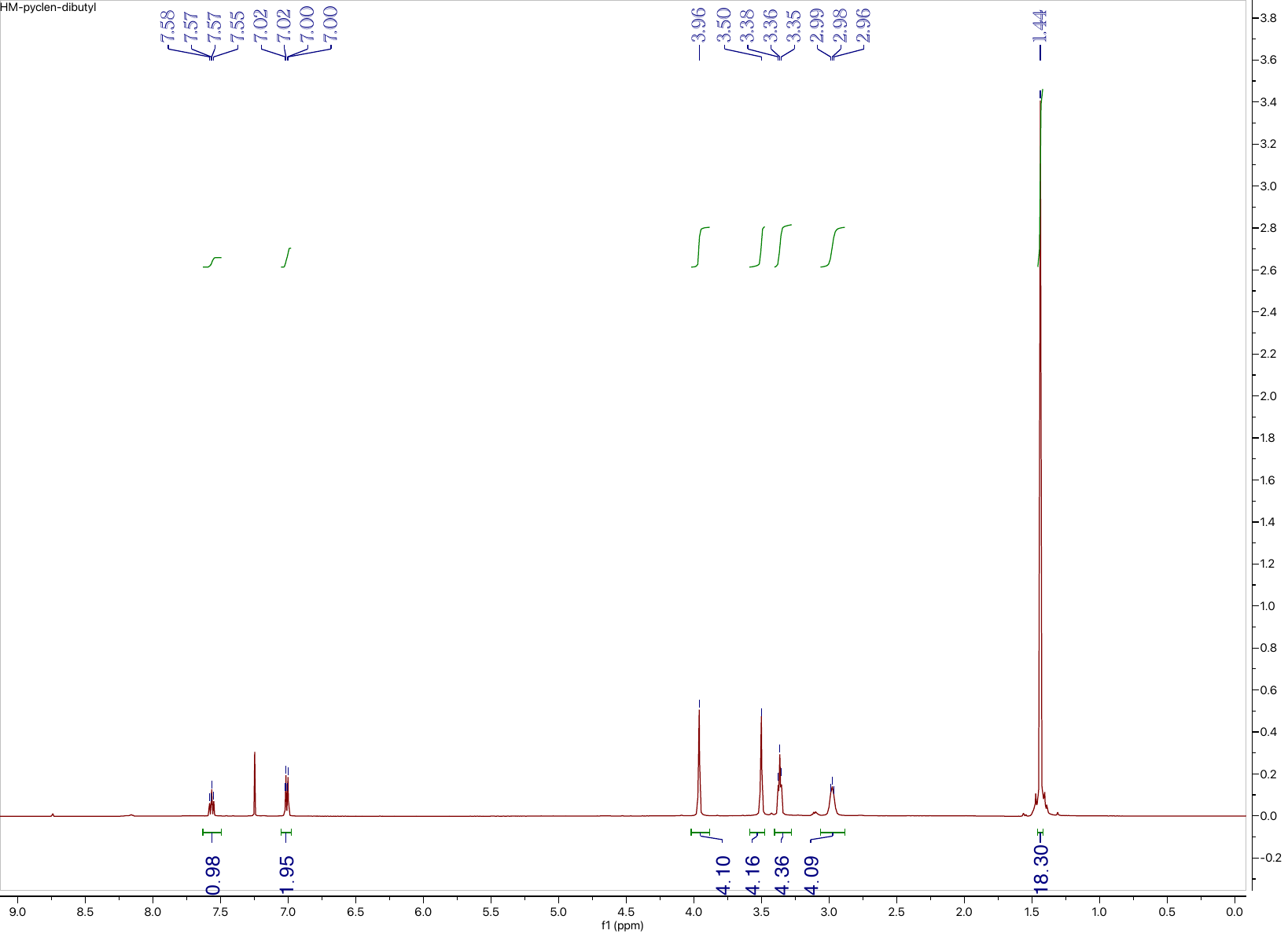

**Compound 2: ^13^C NMR (126 MHz, CDCl_3_)**

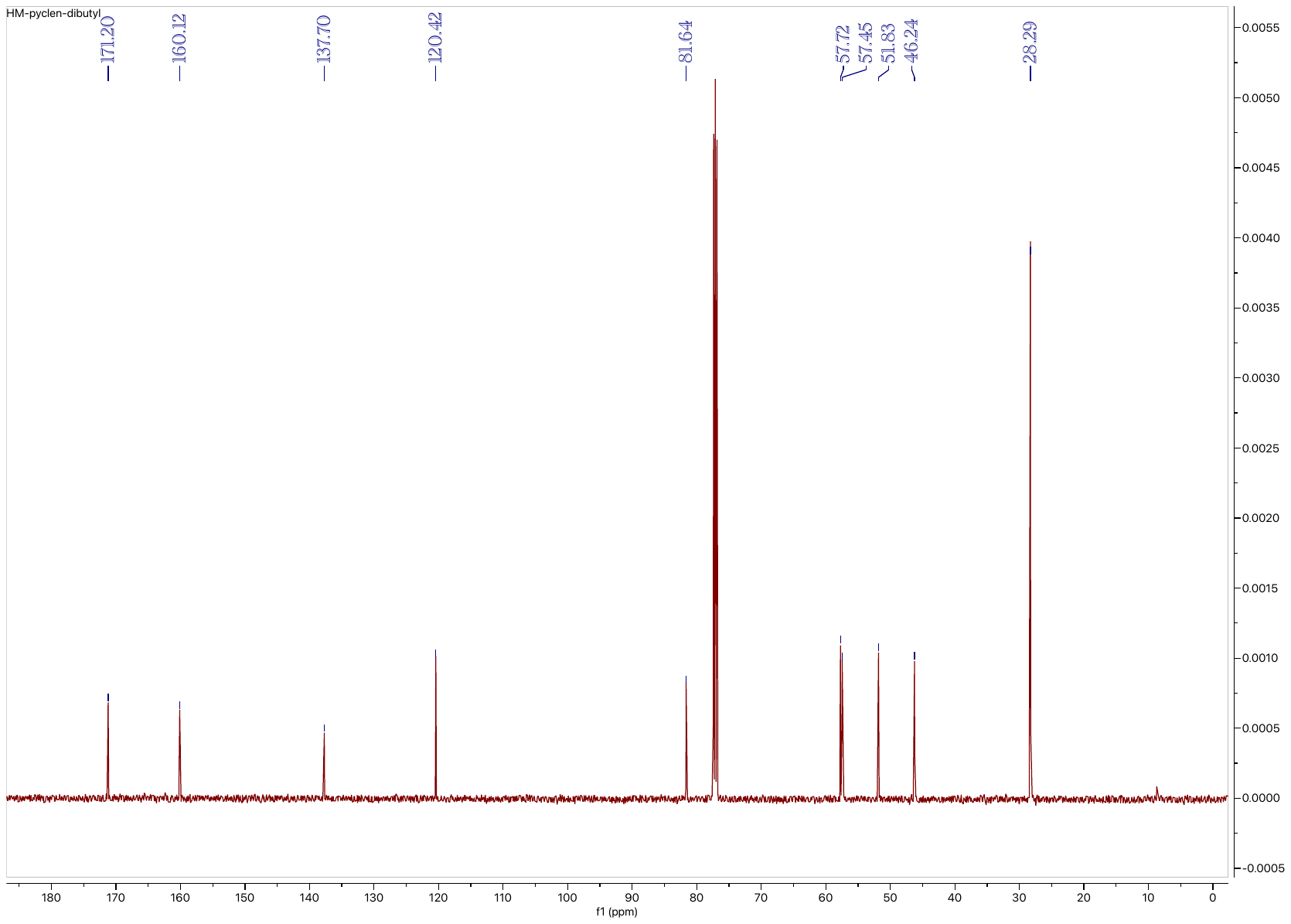

**Compound 3: ^1^H NMR (500 MHz, CDCl_3_)**

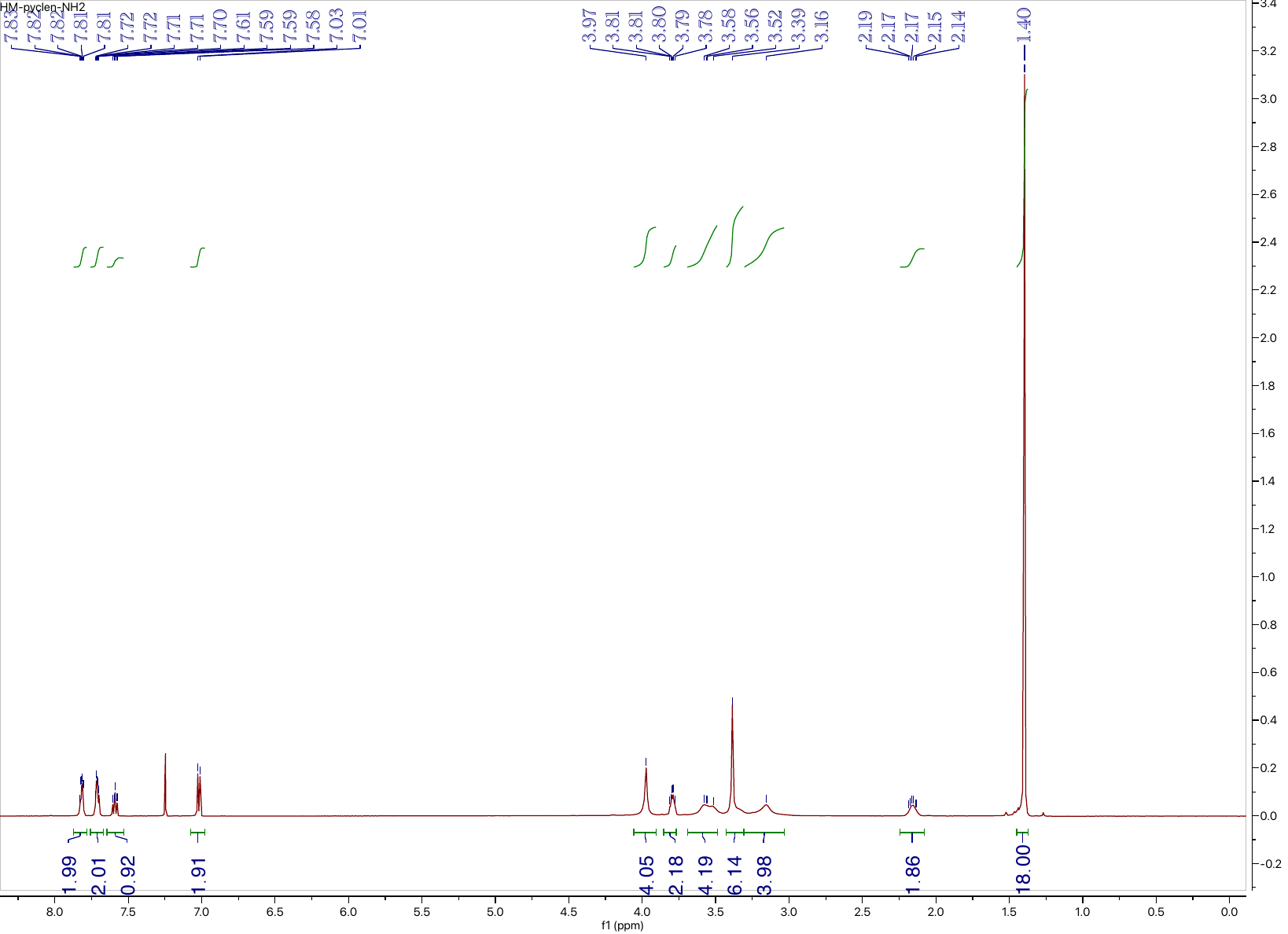

**Compound 3: ^13^C NMR (126 MHz, CDCl_3_)**

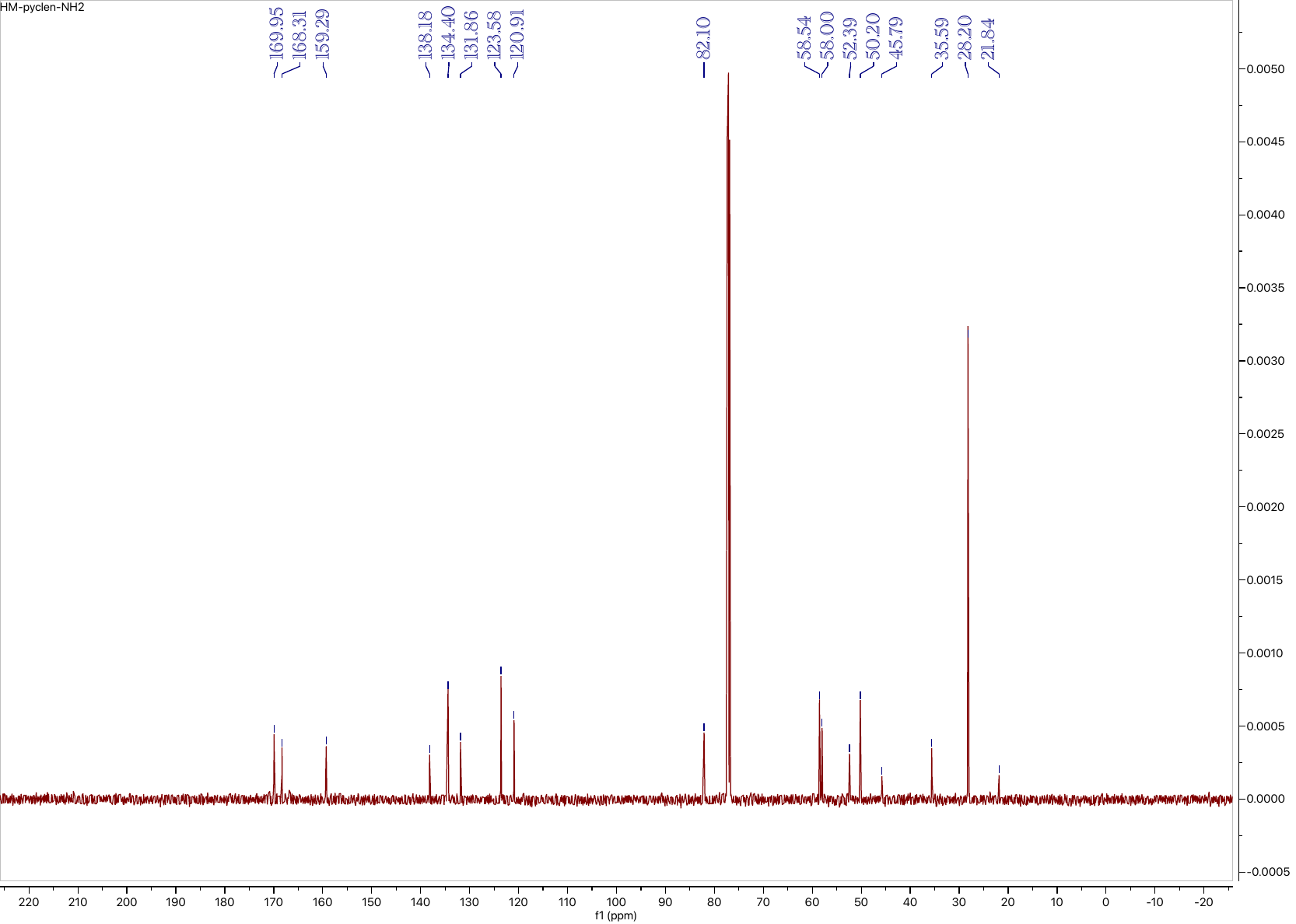

**Compound 4: ^1^H NMR (500 MHz, CDCl_3_)**

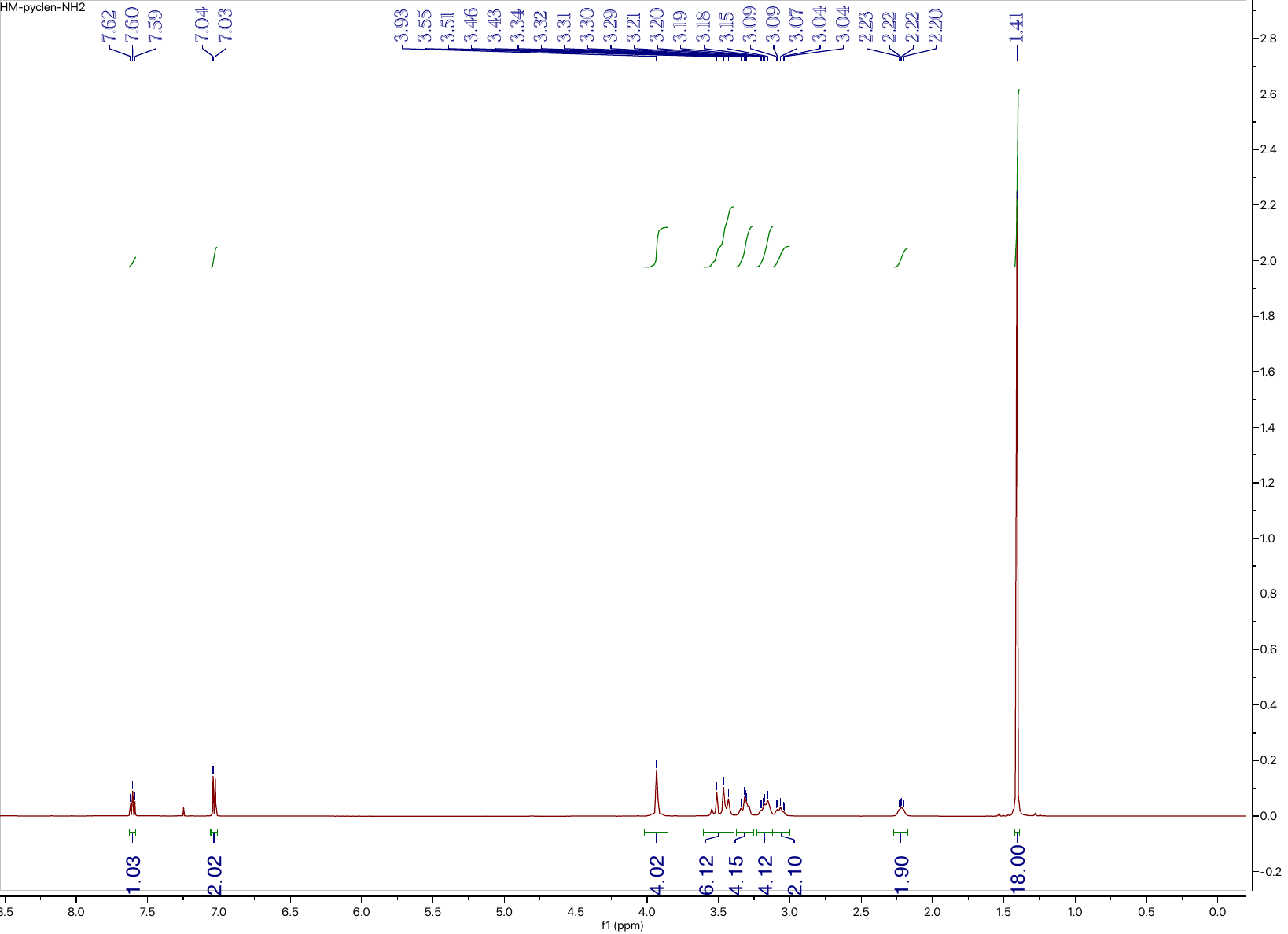

**Compound 4: ^13^C NMR (126 MHz, CDCl_3_)**

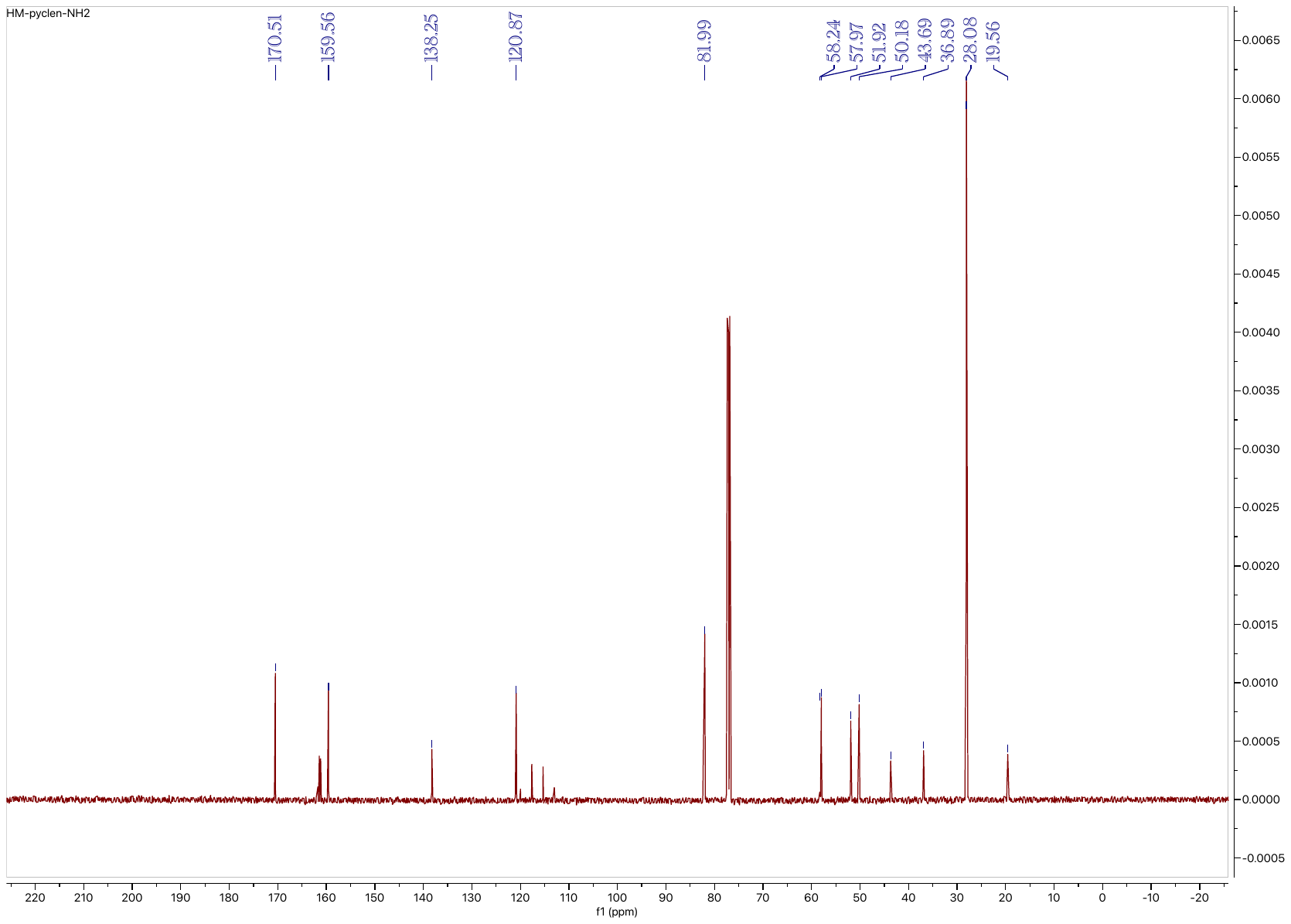

**Compound 6: ^1^H NMR (500 MHz, CDCl_3_)**

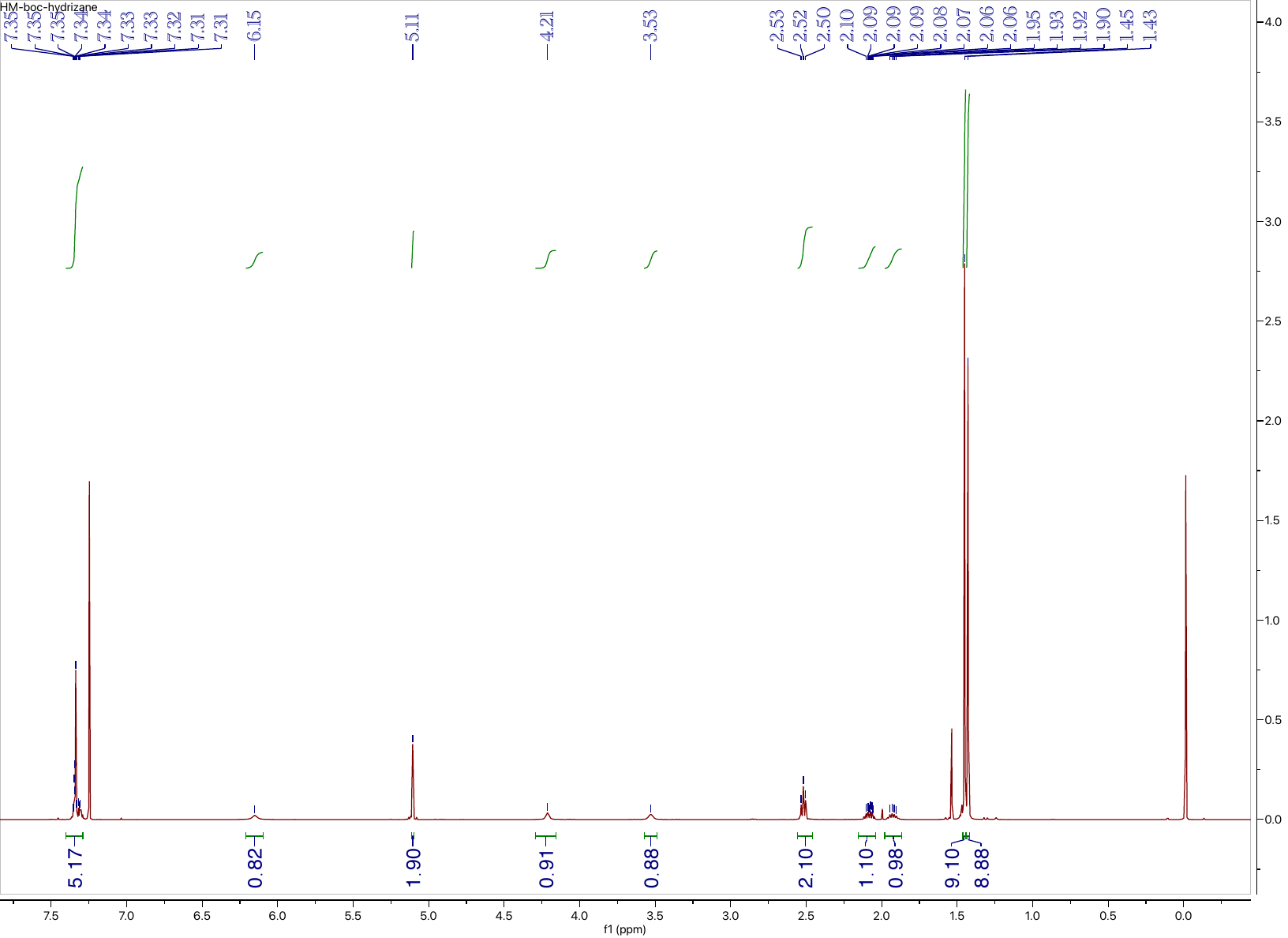

**Compound 6: ^13^C NMR (126 MHz, CDCl_3_)**

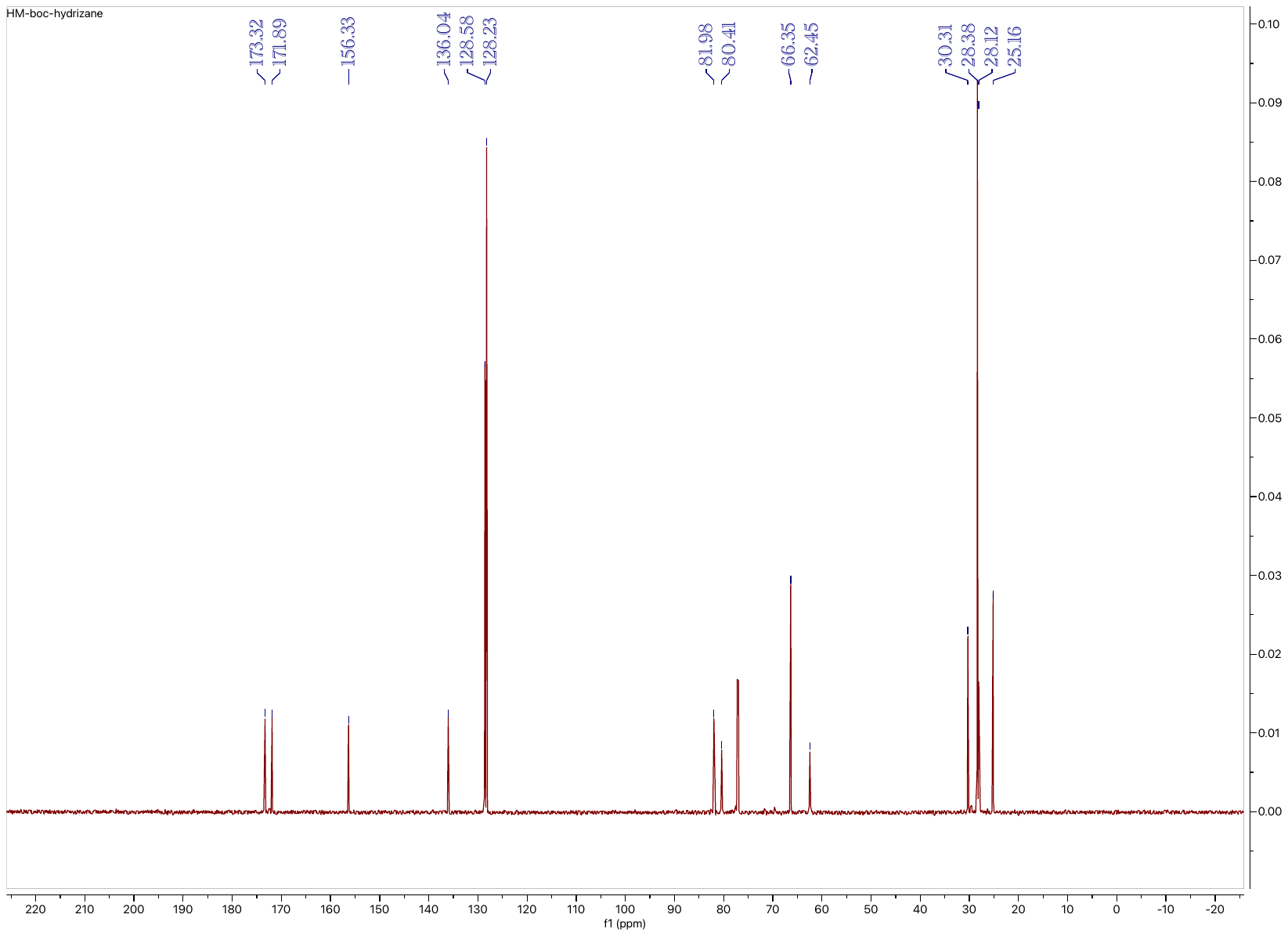

**Compound 7: ^1^H NMR (500 MHz, CDCl_3_)**

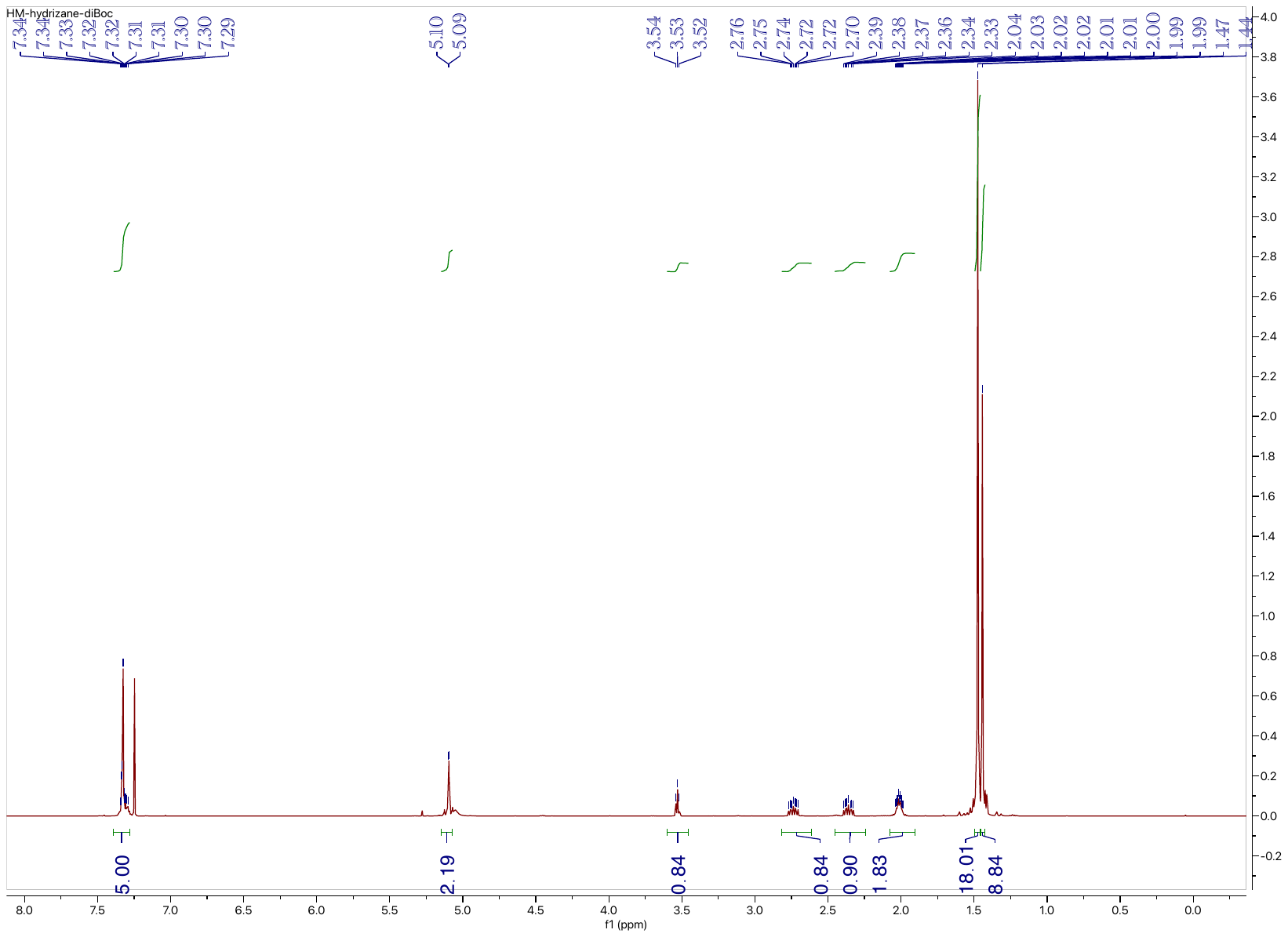

**Compound 7: ^13^C NMR (126 MHz, CDCl_3_)**

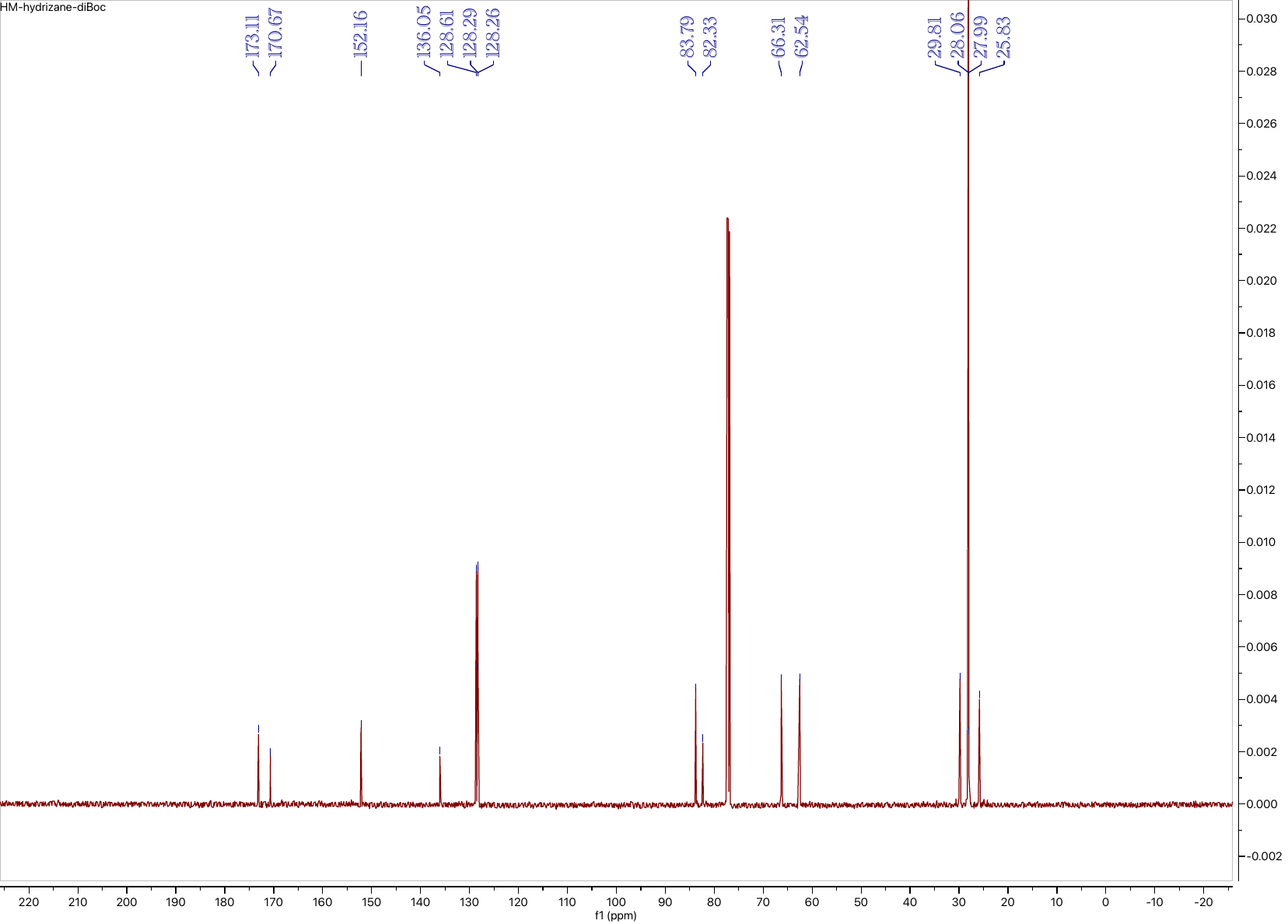

**Compound 8: ^1^H NMR (126 MHz, CDCl_3_)**

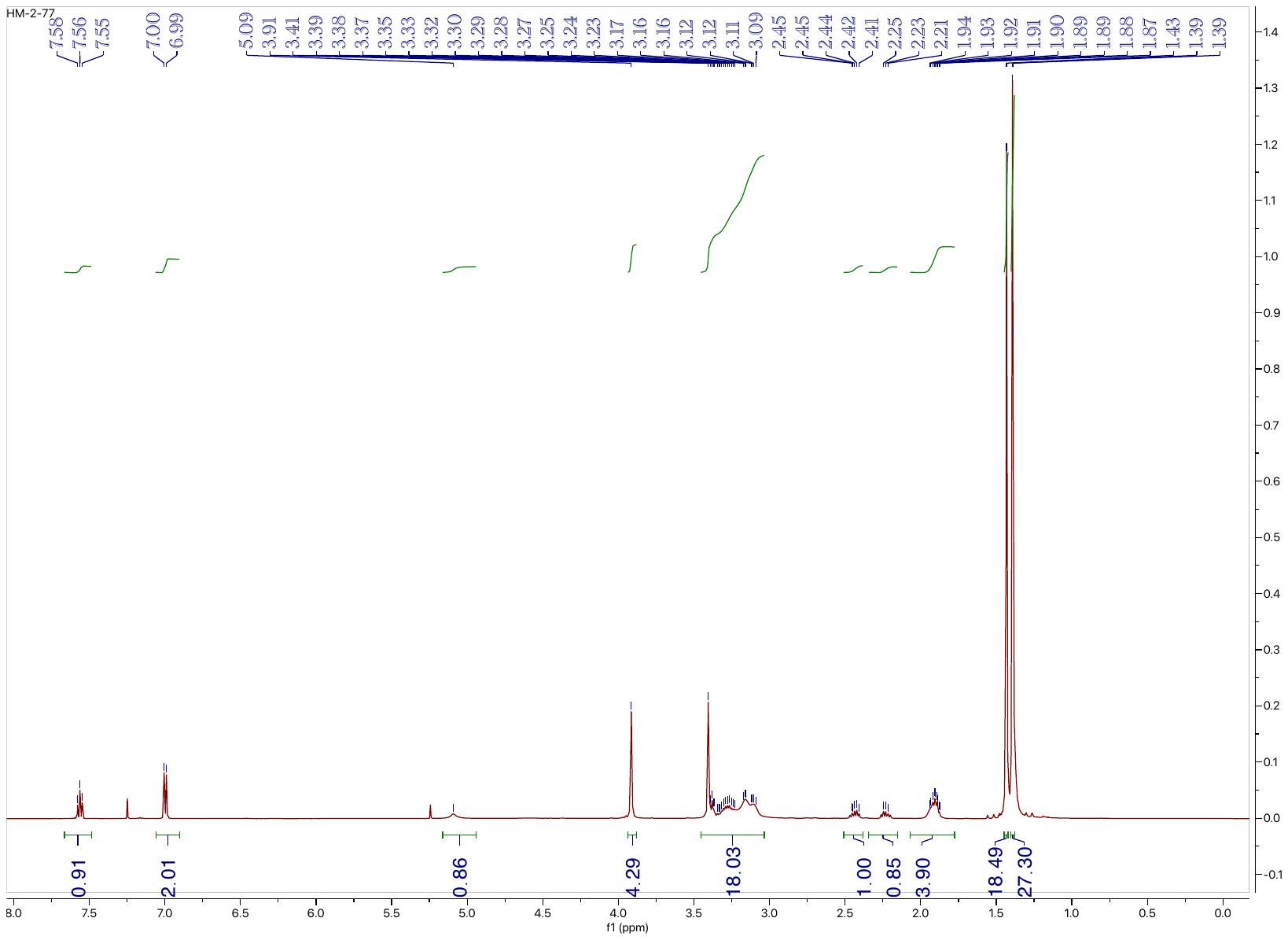

**Compound 8: ^13^C NMR (126 MHz, CDCl_3_)**

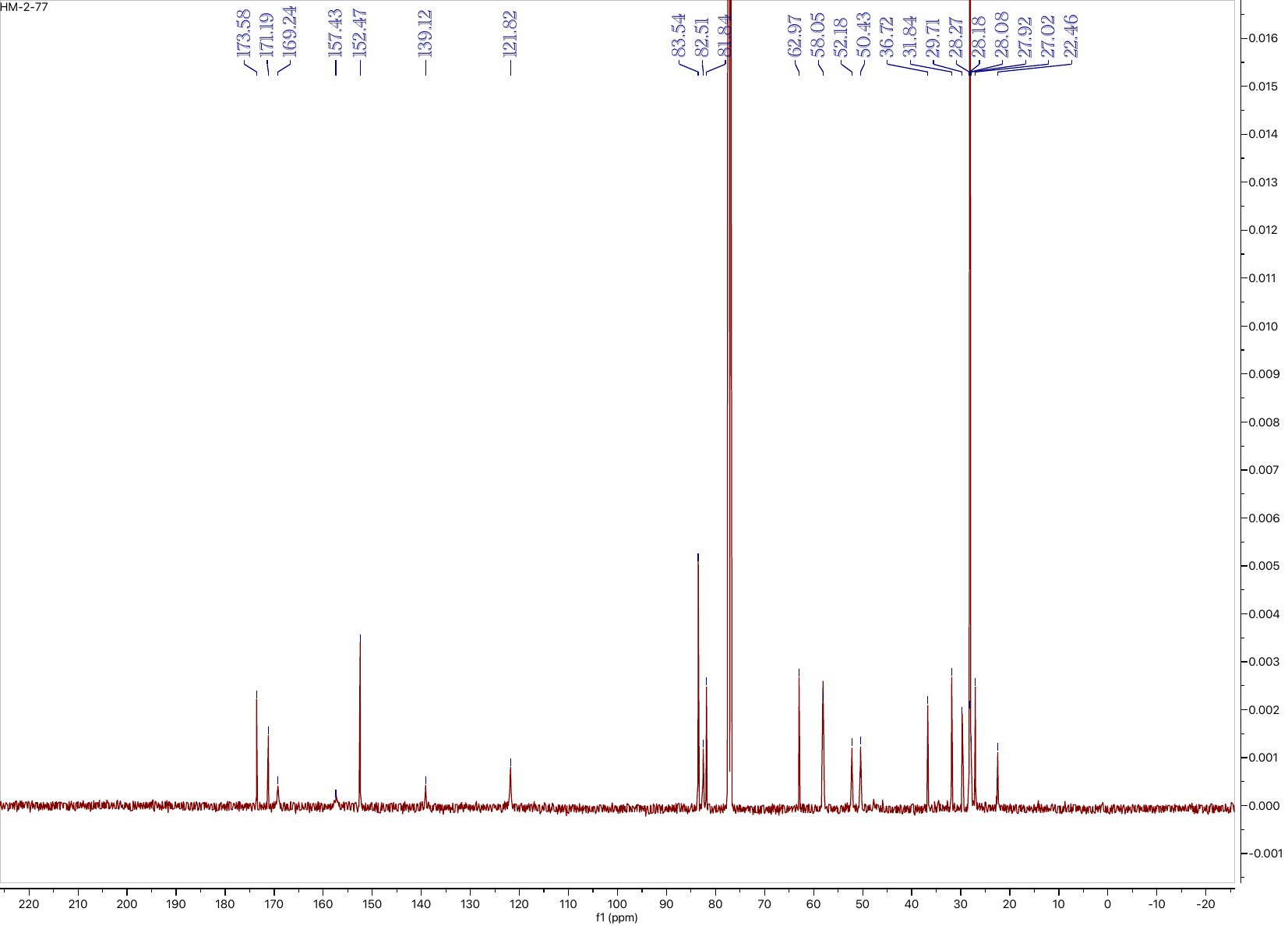

**Compound L1: ^1^H NMR (500 MHz, D_2_O)**

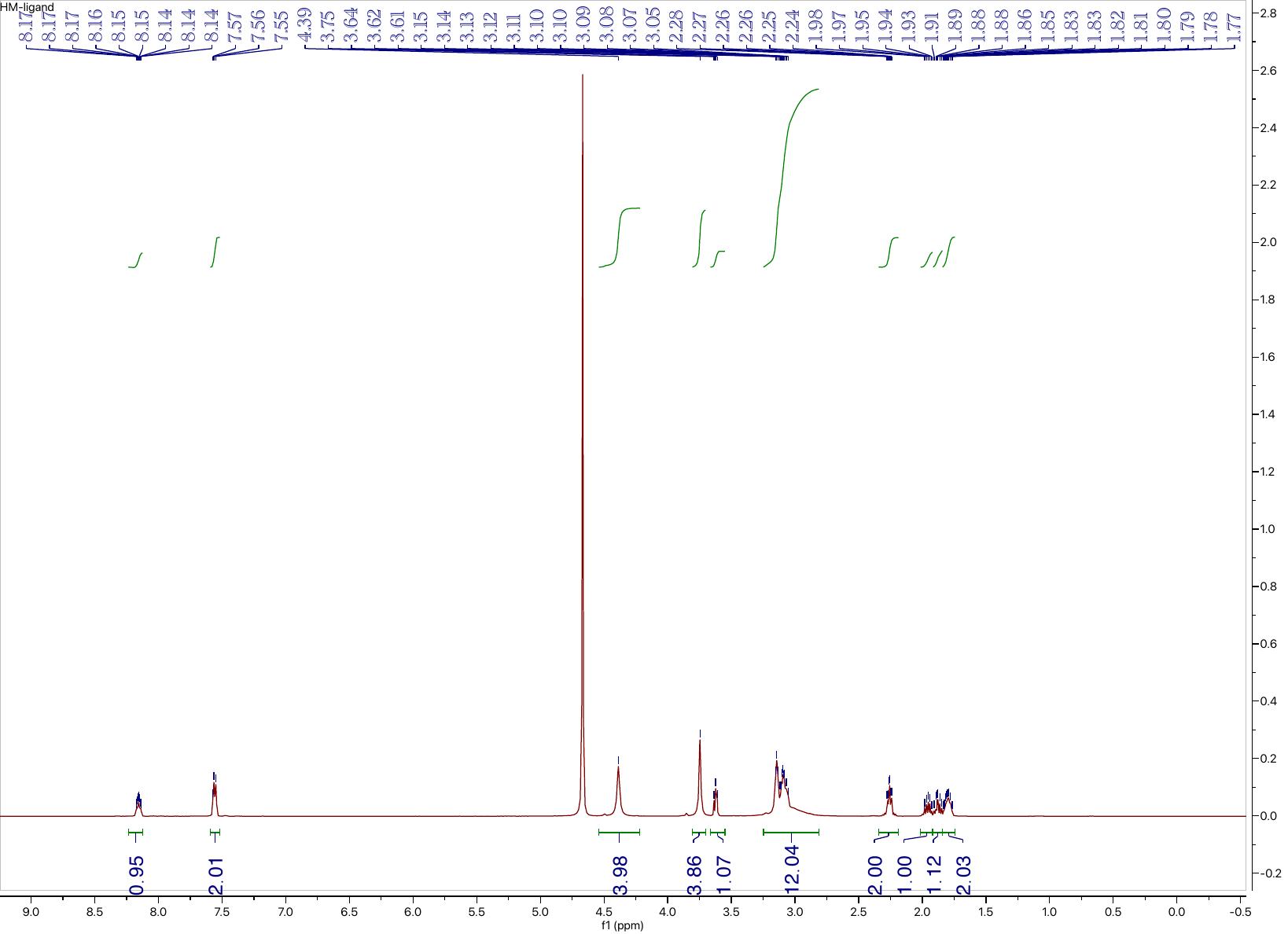

**Compound L1: ^13^C NMR (126 MHz, D_2_O)**

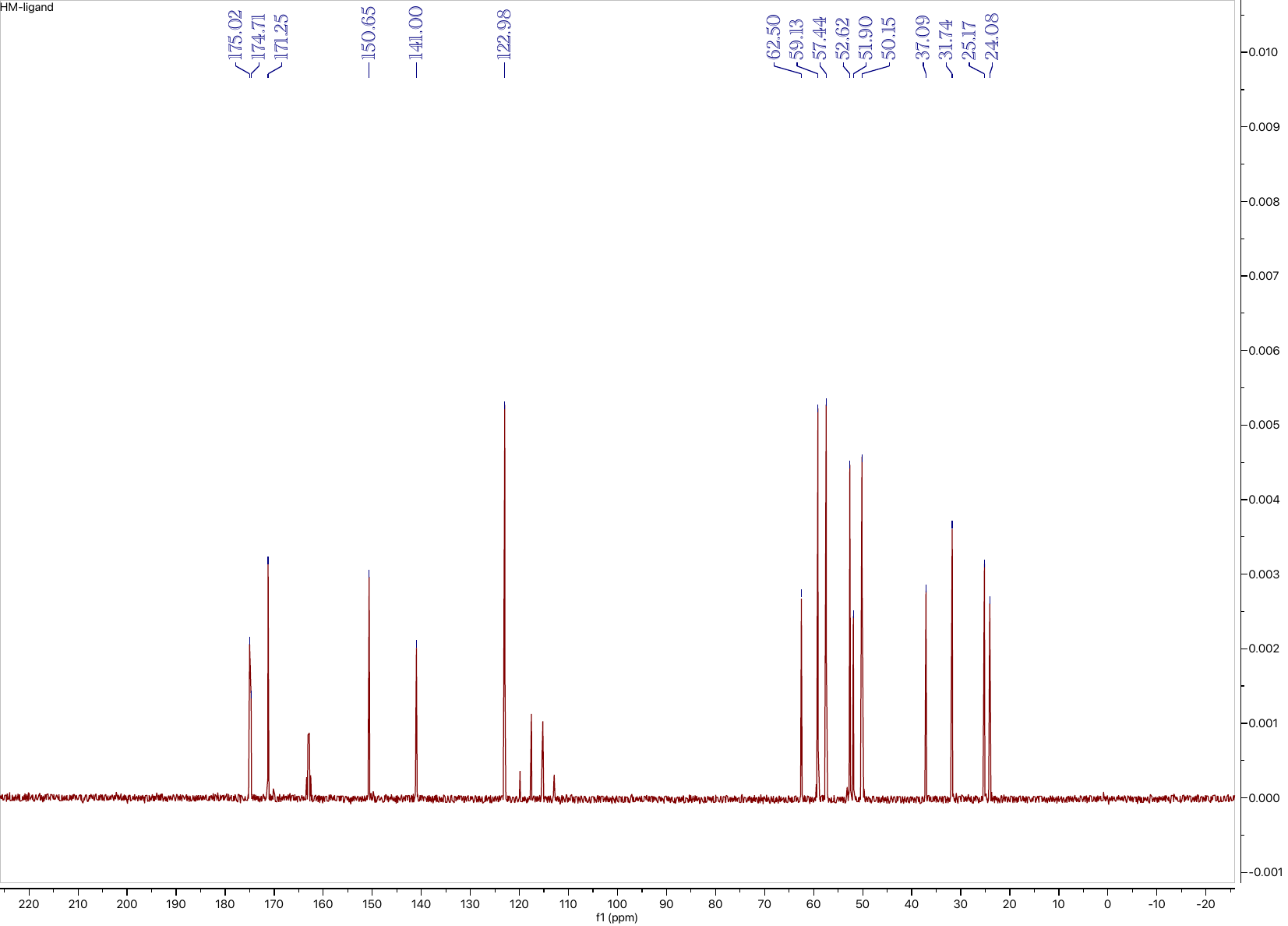

**Compound 10: ^1^H NMR (500 MHz, CDCl_3_)**

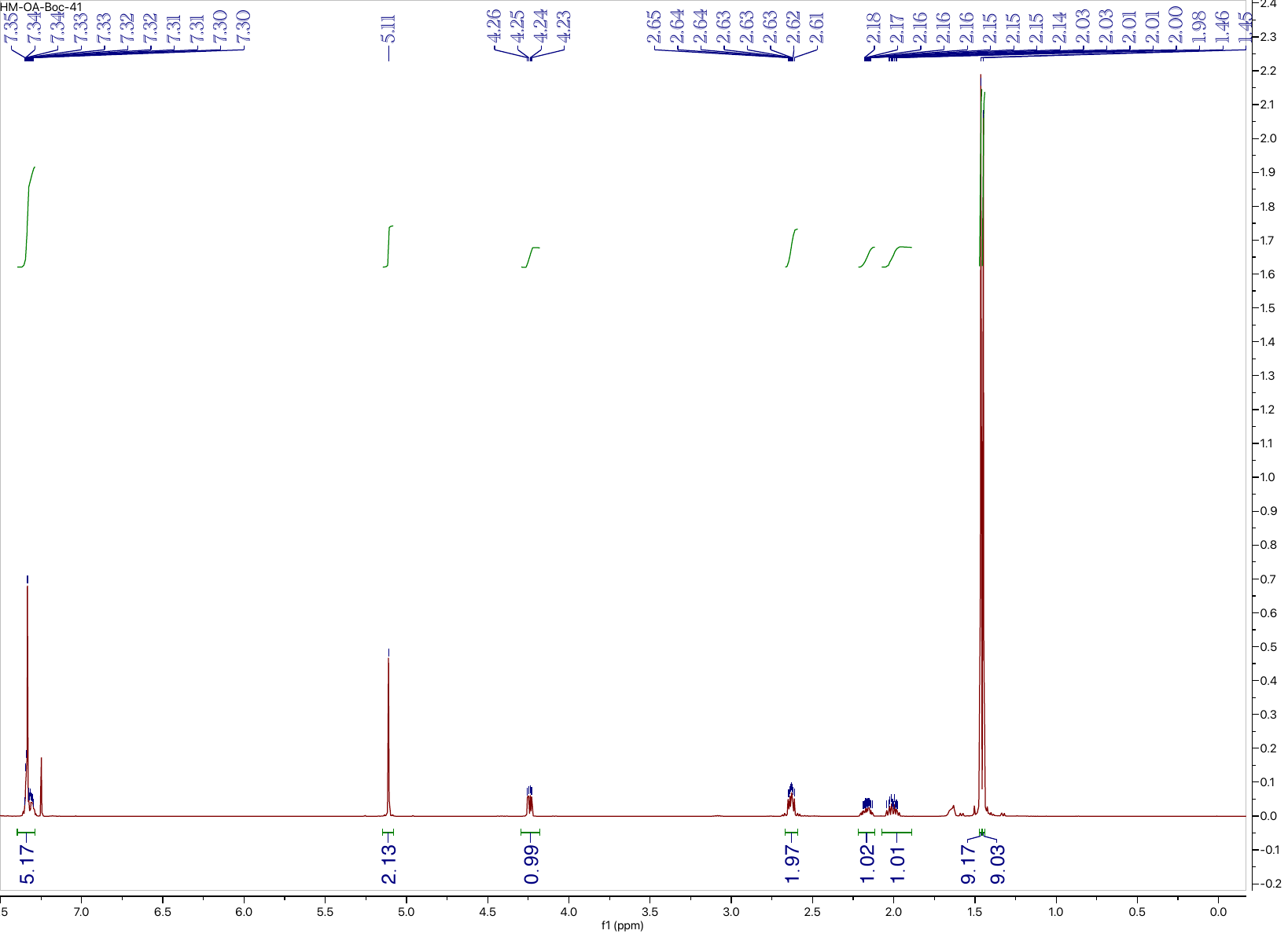

**Compound 10: ^13^C NMR (126 MHz, CDCl_3_)**

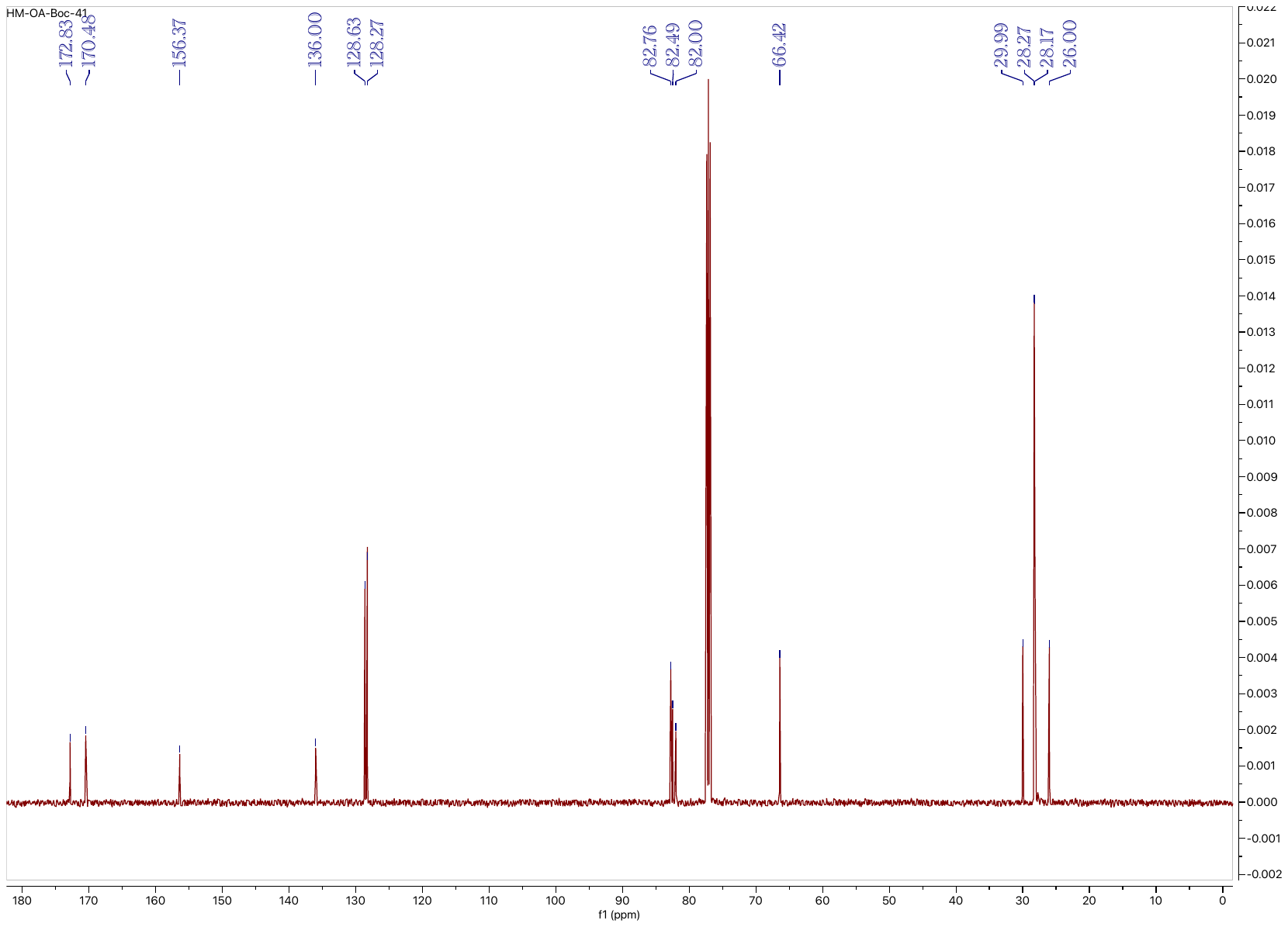

**Compound 11: ^1^H NMR (500 MHz, CDCl_3_)**

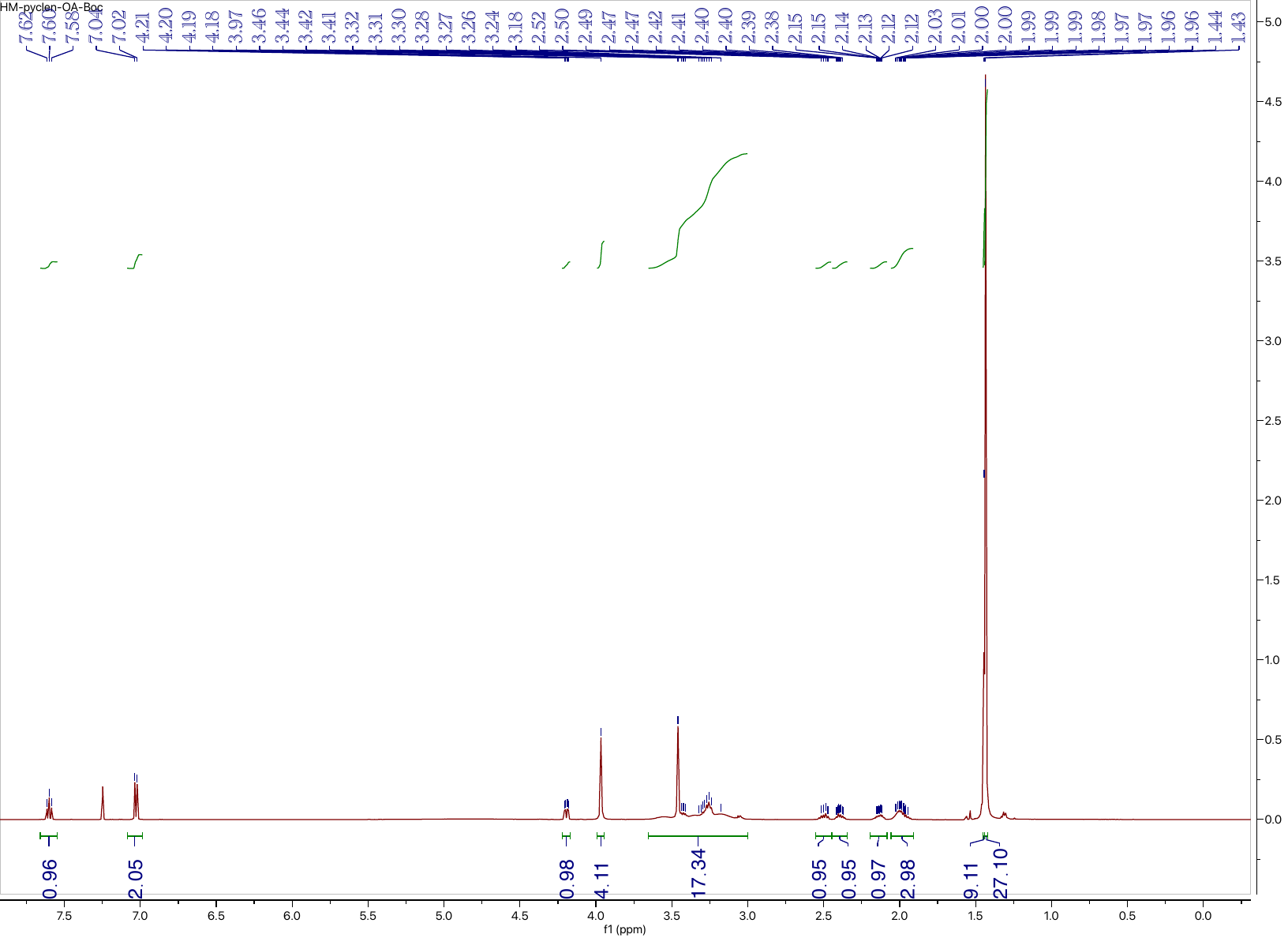

**Compound 11: ^13^C NMR (126 MHz, CDCl_3_)**

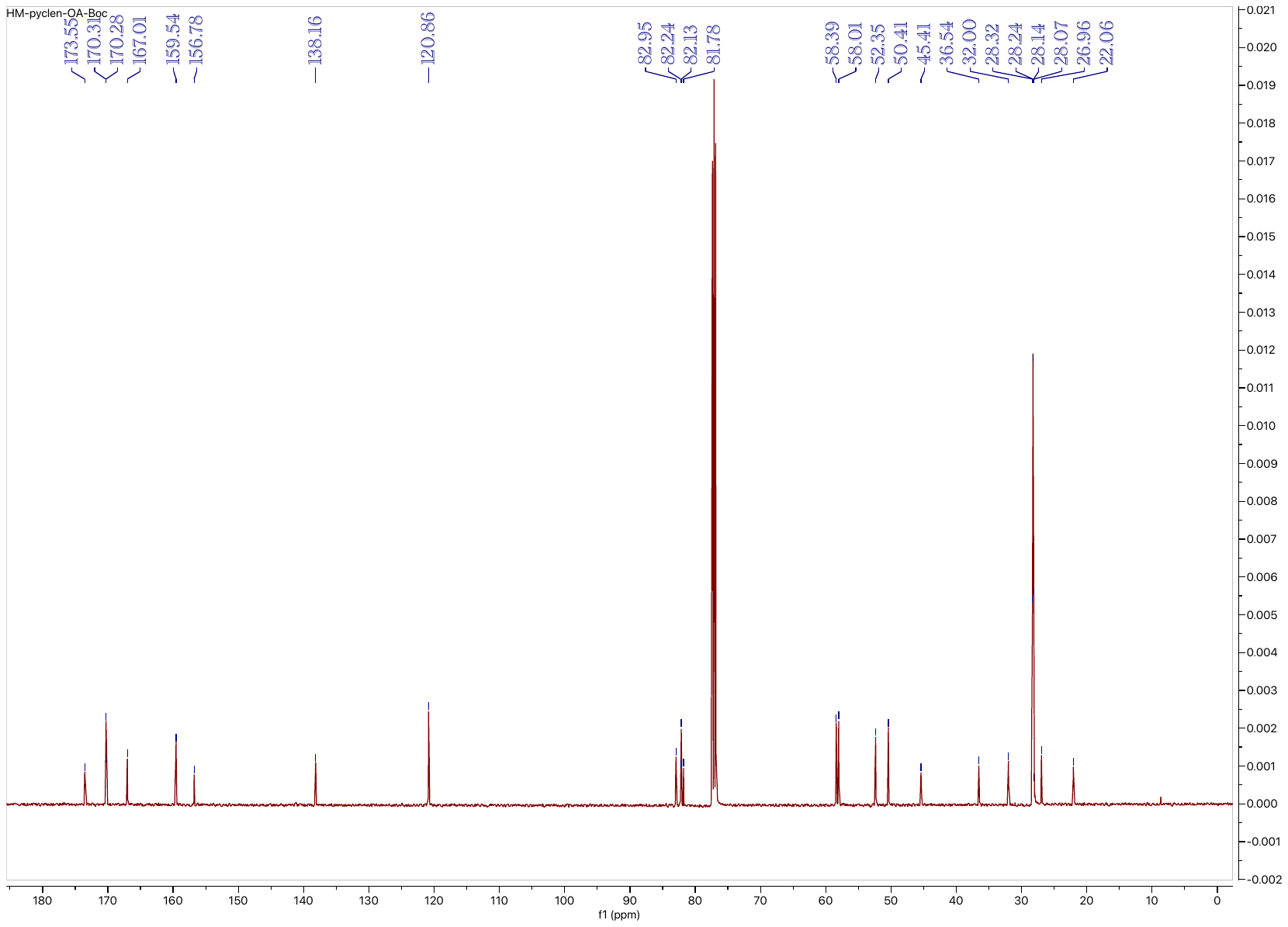

**Compound L2: ^1^H NMR (500 MHz, D_2_O)**

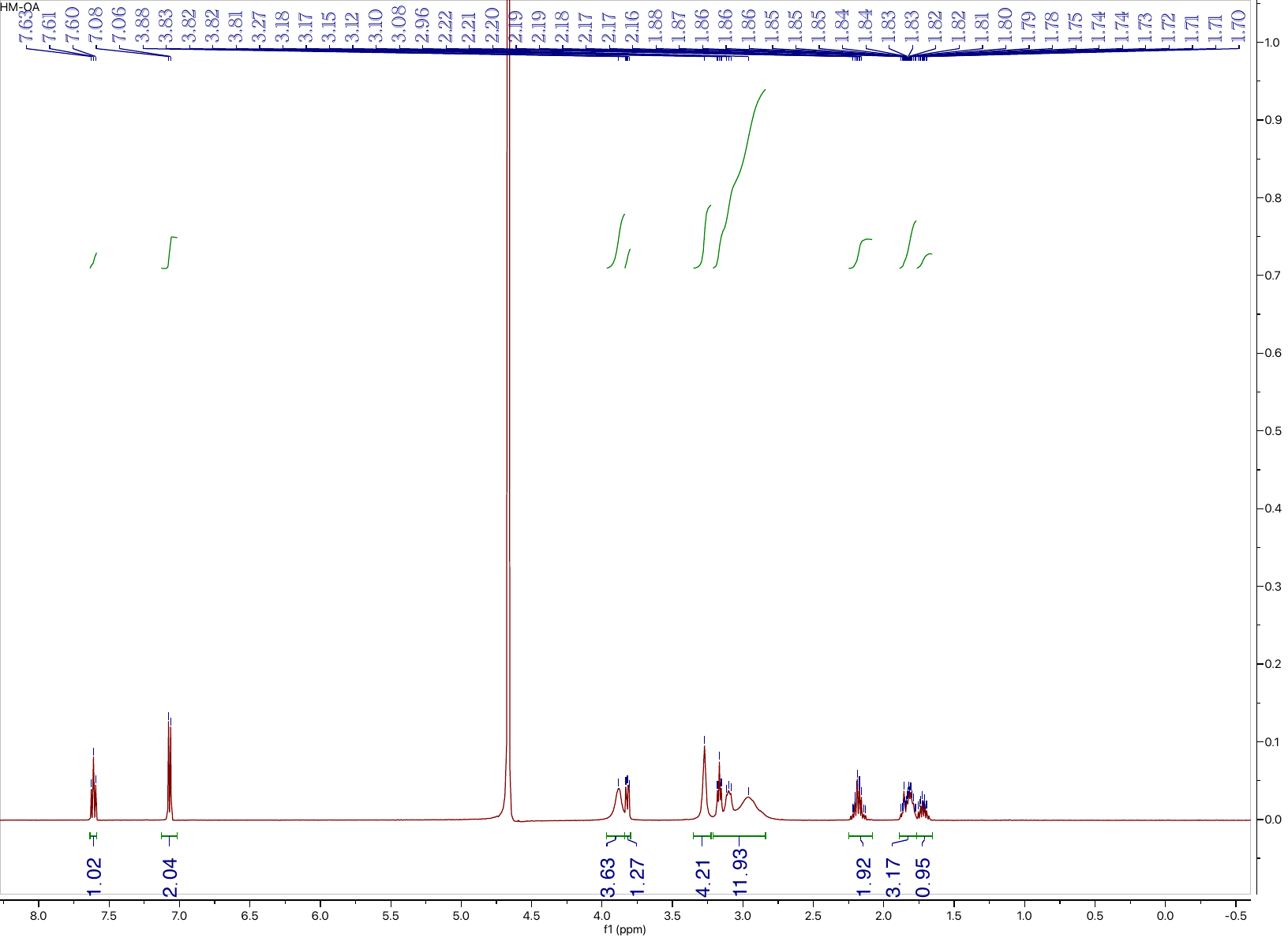

**Compound L2: ^13^C NMR (126 MHz, D_2_O)**

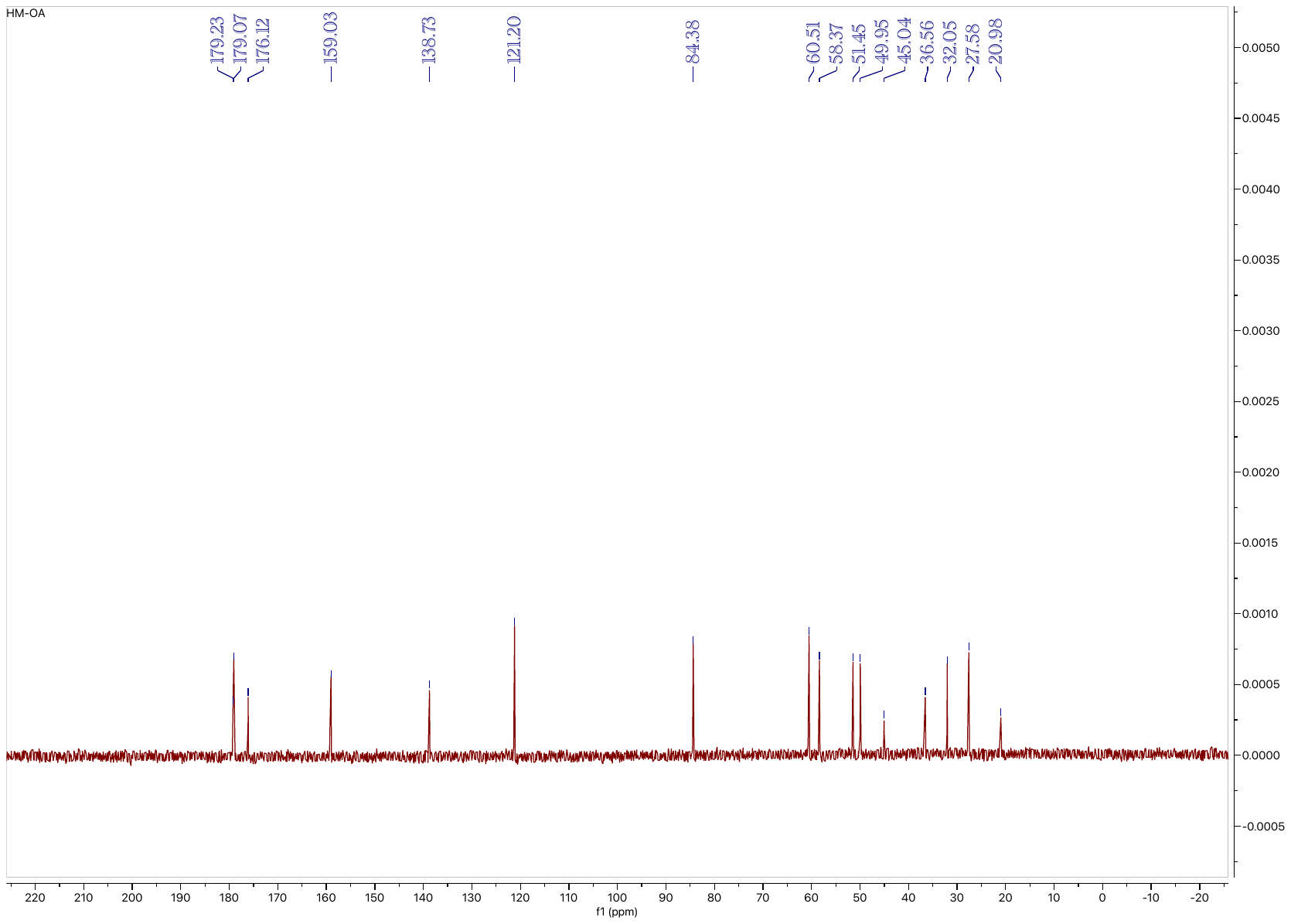

**Compound 13: ^1^H NMR (500 MHz, CDCl_3_)**

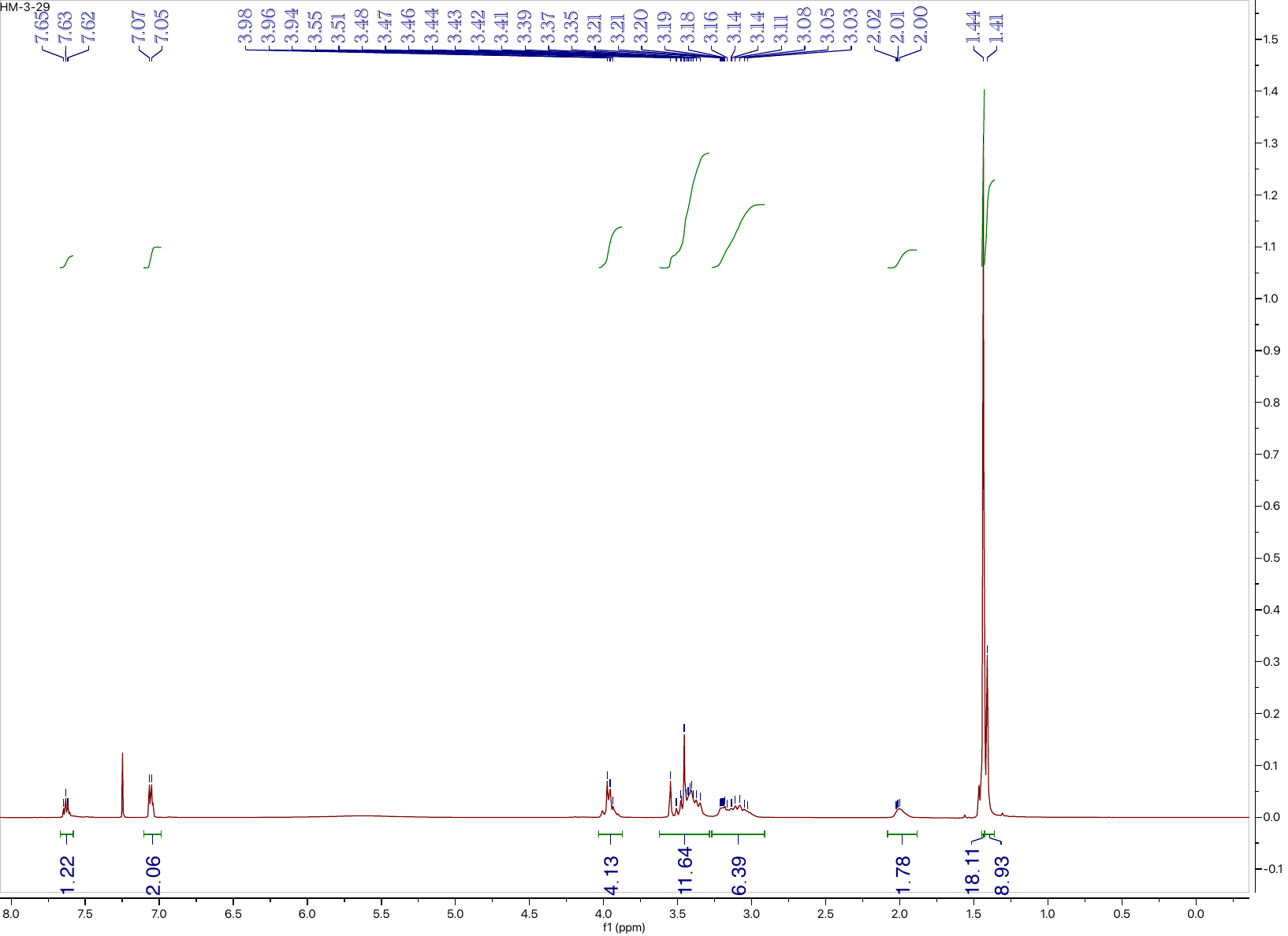

**Compound 13: ^13^C NMR (126 MHz, CDCl_3_)**

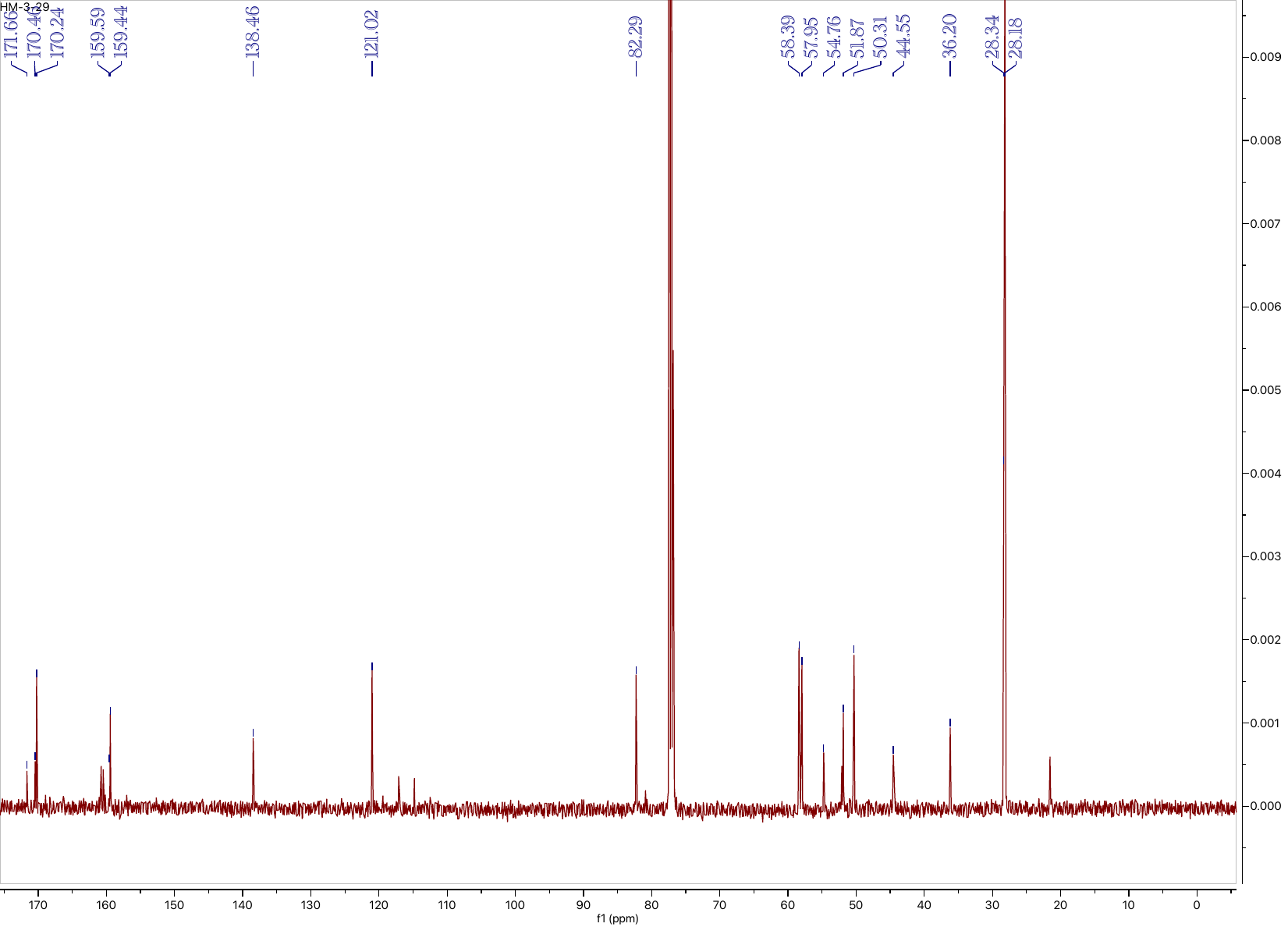

**Compound L3: ^1^H NMR (500 MHz, D_2_O)**

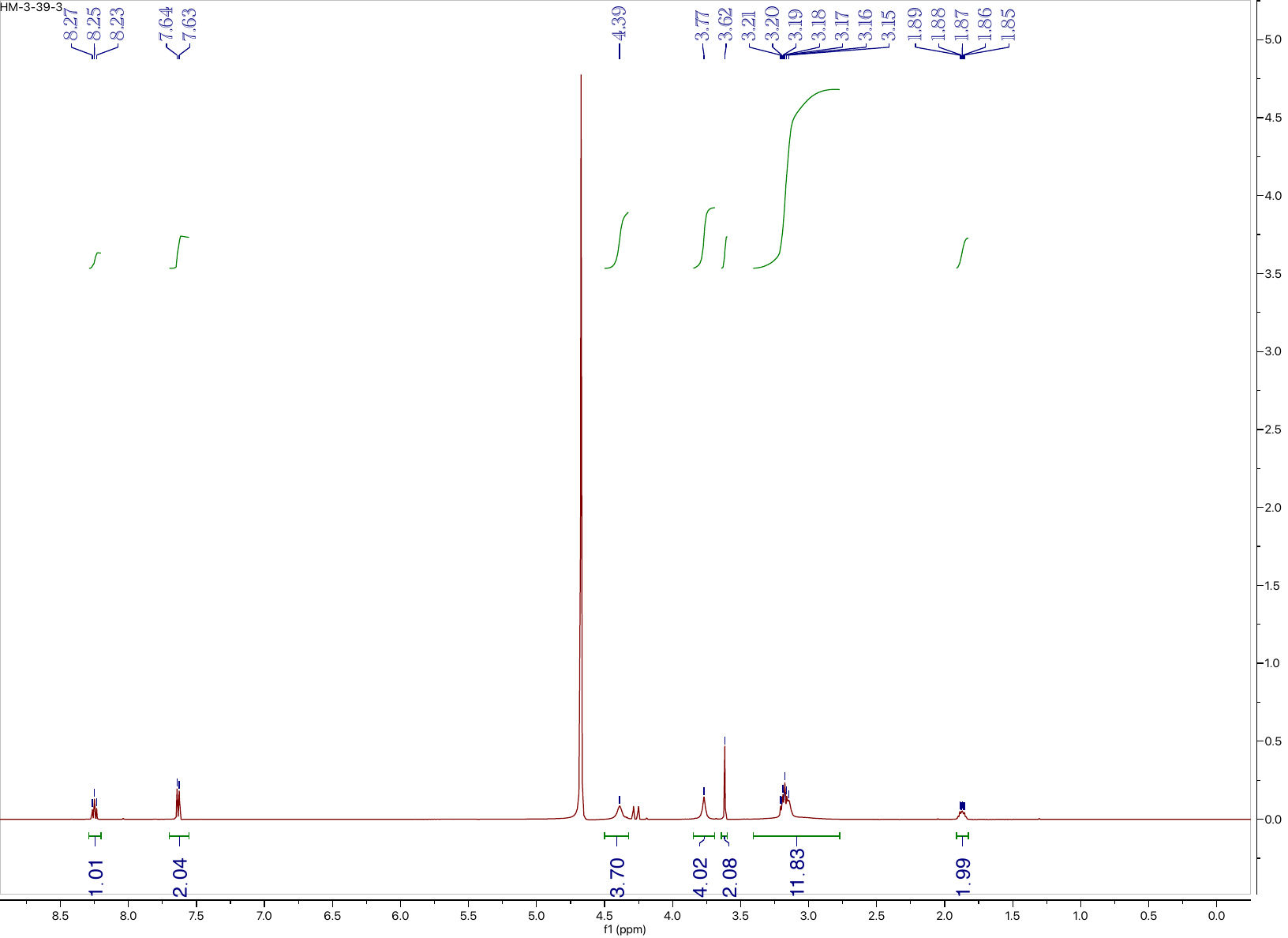

**Compound L3: ^13^C NMR (126 MHz, D_2_O)**

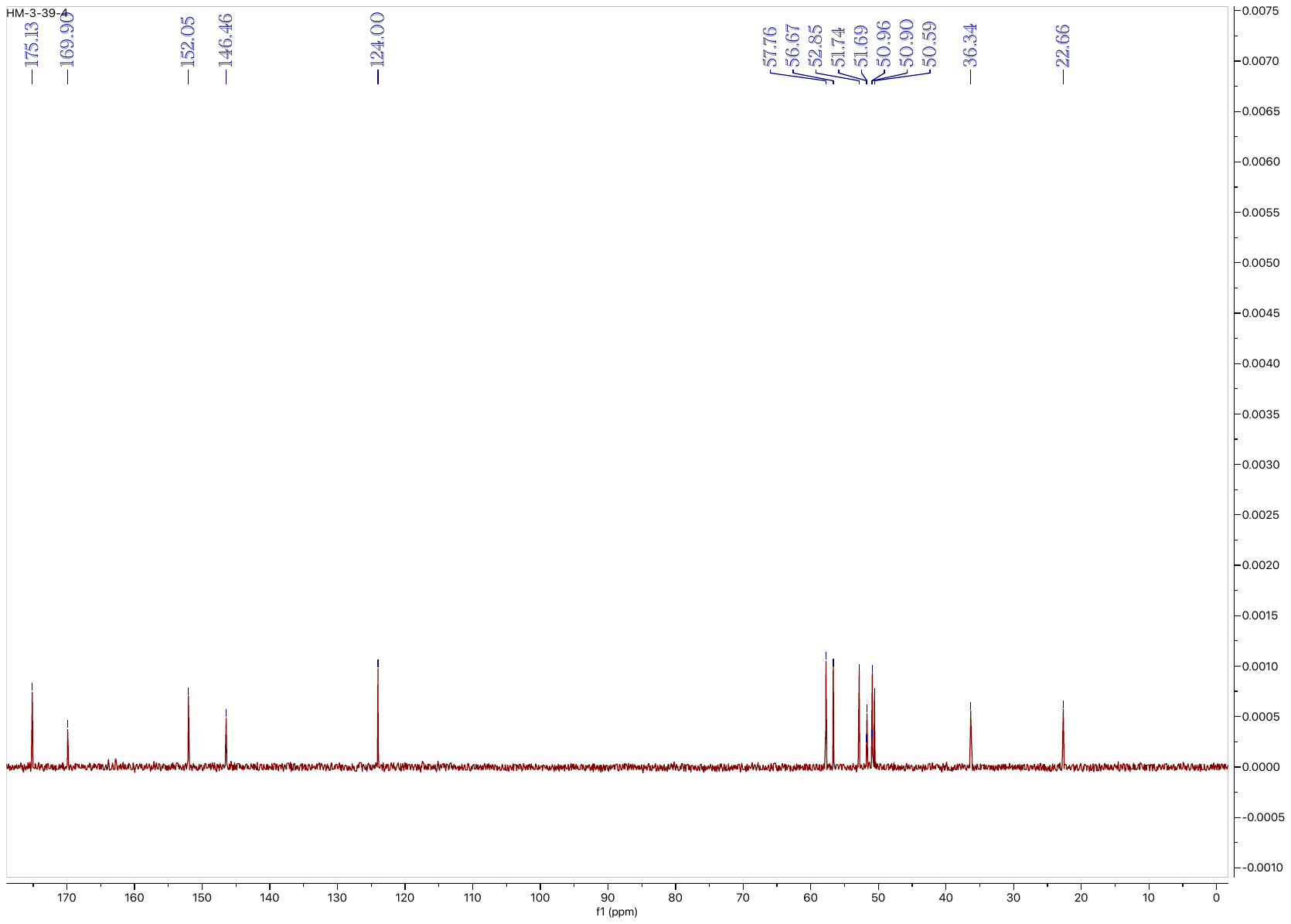

**Compound 14: ^1^H NMR (500 MHz, CDCl_3_)**

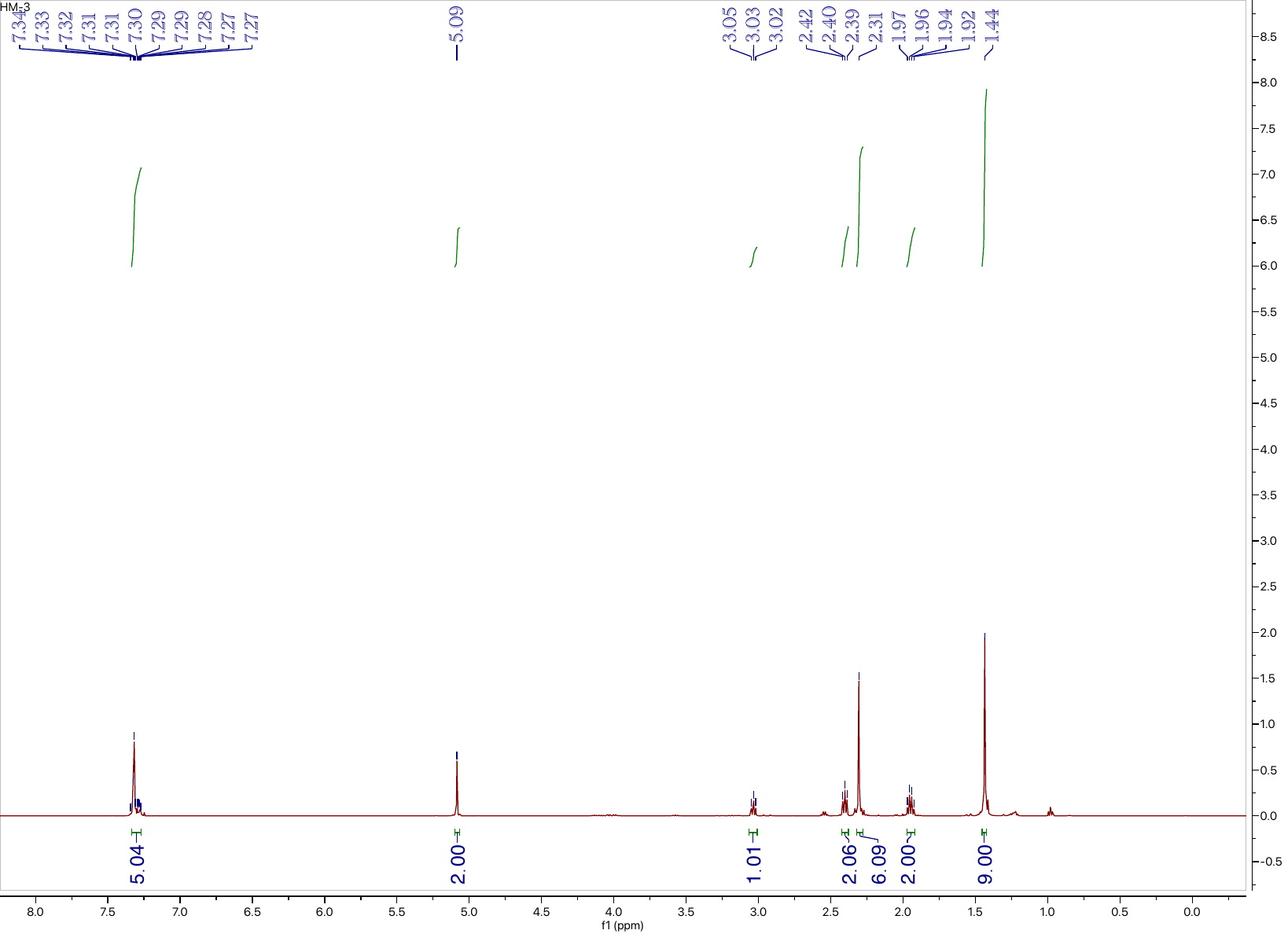

**Compound 14: ^13^C NMR (126 MHz, CDCl_3_)**

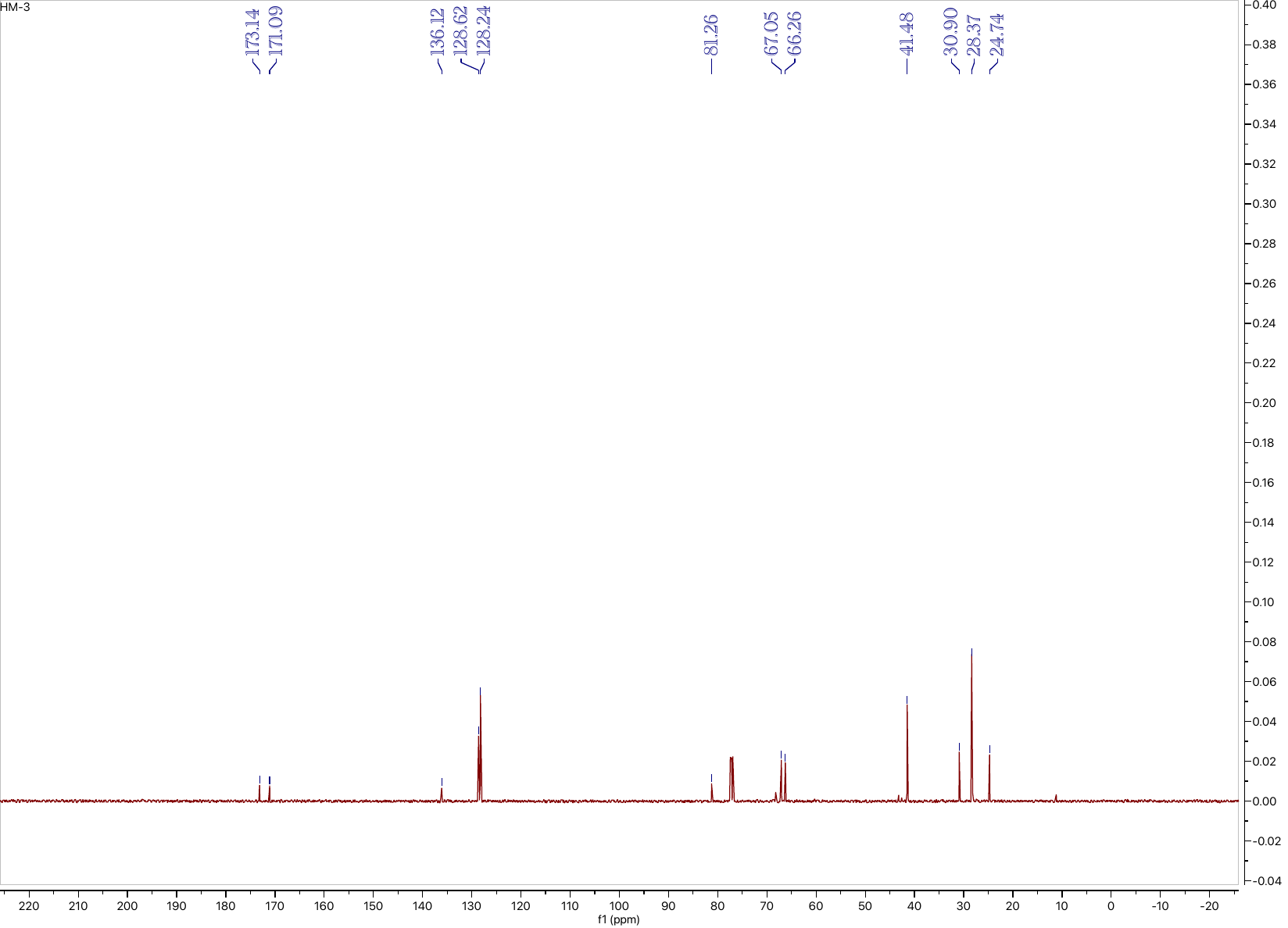

**Compound 15: ^1^H NMR (500 MHz, CDCl_3_)**

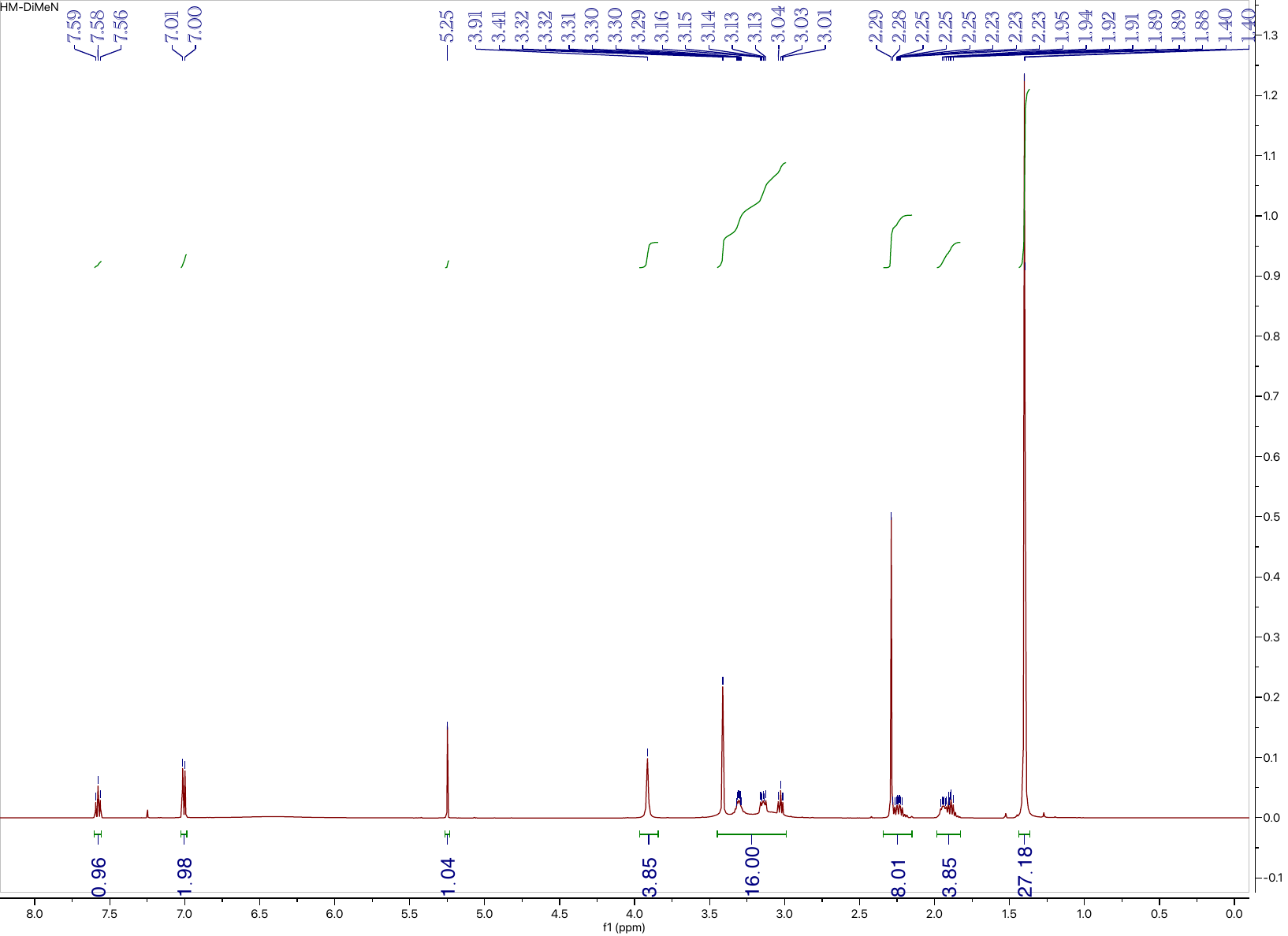

**Compound 15: ^13^C NMR (126 MHz, CDCl_3_)**

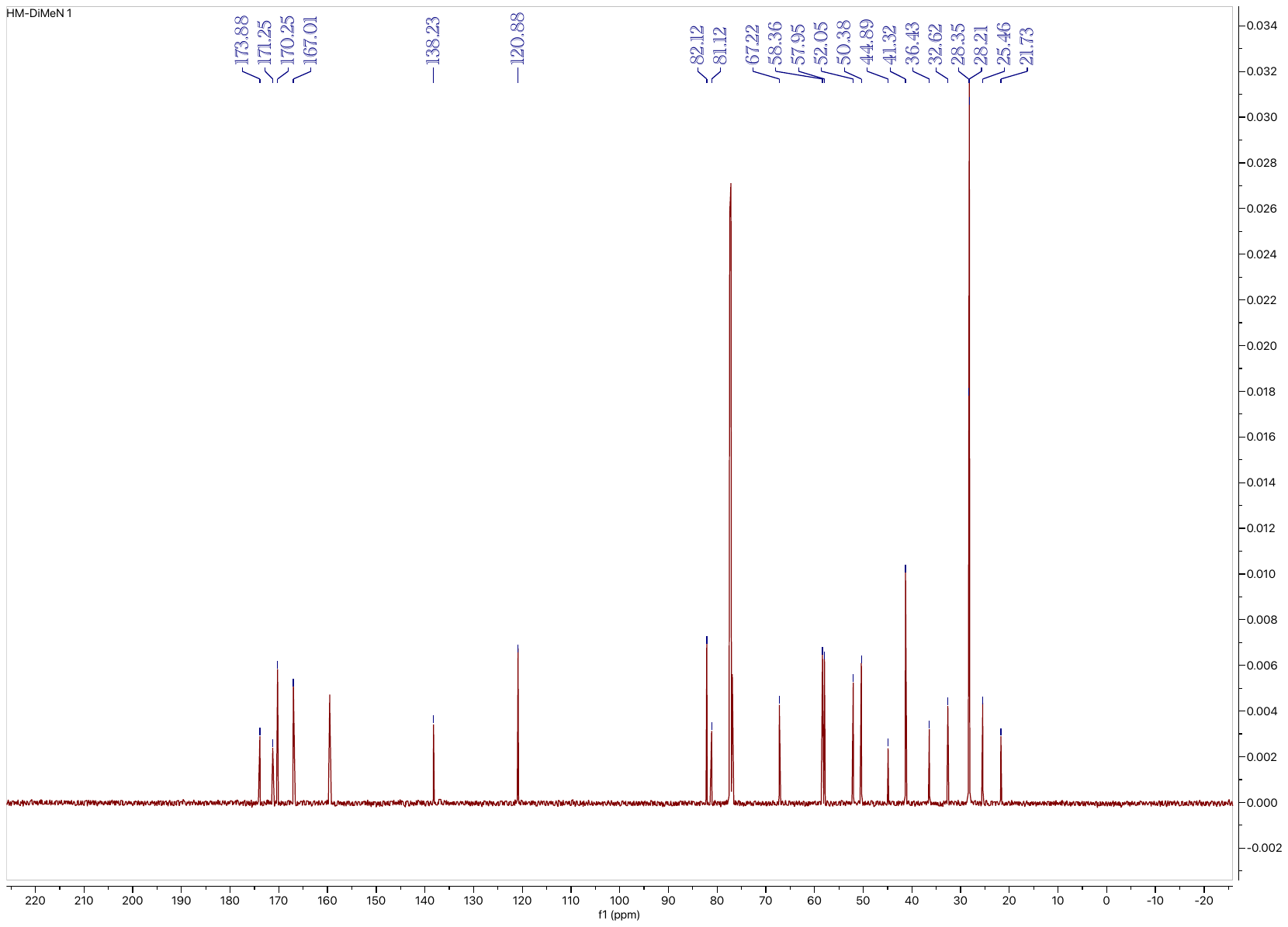

**Compound L4: ^1^H NMR (500 MHz, D_2_O)**

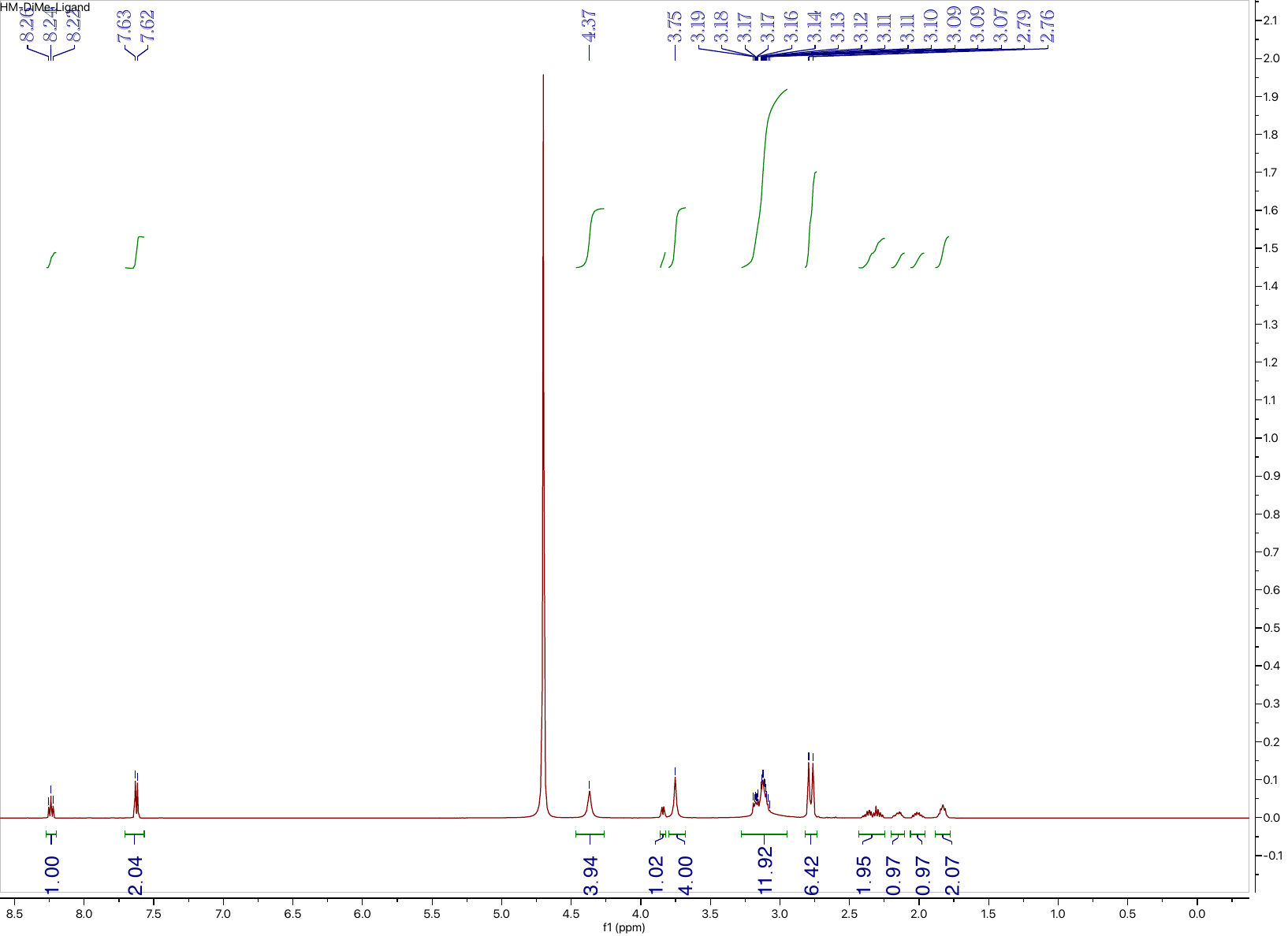

**Compound L4: ^13^C NMR (126 MHz, D_2_O)**

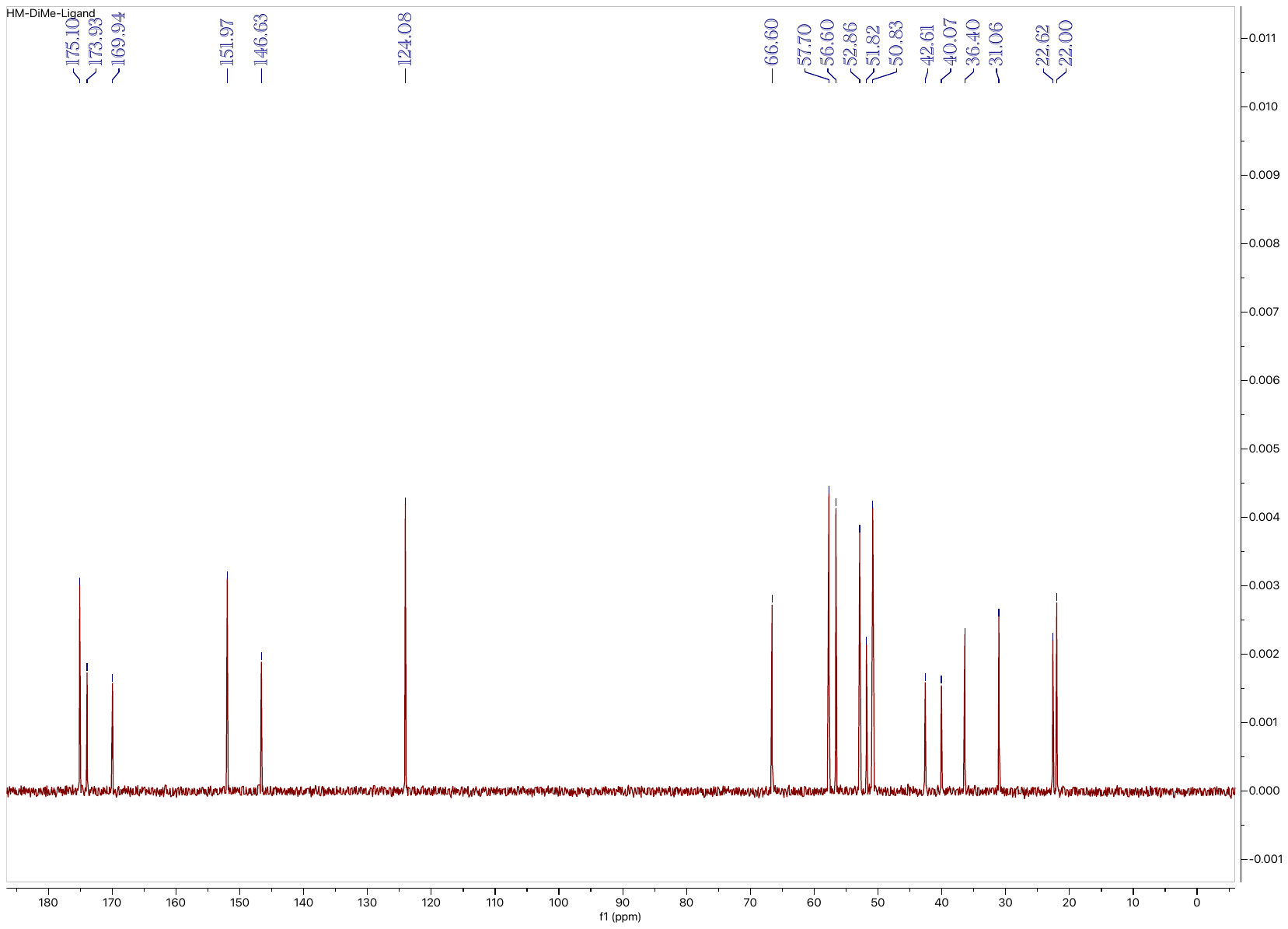
